# Machine learning models based on molecular descriptors to predict human and environmental toxicological factors in continental freshwater

**DOI:** 10.1101/2021.07.20.453034

**Authors:** Rémi Servien, Eric Latrille, Dominique Patureau, Arnaud Hélias

## Abstract

It is a real challenge for life cycle assessment practitioners to identify all relevant substances contributing to the ecotoxicity. Once this identification has been made, the lack of corresponding ecotoxicity factors can make the results partial and difficult to interpret. So, it is a real and important challenge to provide ecotoxicity factors for a wide range of compounds. Nevertheless, obtaining such factors using experiments is tedious, time-consuming, and made at a high cost. A modeling method that could predict these factors from easy-to-obtain information on each chemical would be of great value. Here, we present such a method, based on machine learning algorithms, that used molecular descriptors to predict two specific endpoints in continental freshwater for ecotoxicological and human impacts. The different tested machine learning algorithms show good performances on a learning database and the non-linear methods tend to outperform the linear ones. The cluster-then-predict approaches usually show the best performances which suggests that these predicted models must be derived for somewhat similar compounds. Finally, predictions were derived from the validated model for compounds with missing toxicity/ecotoxicity factors.

**Highlights:**

- Characterization factors (for human health and ecotoxicological impacts) were predicted using molecular descriptors.
- Several linear or non-linear machine learning methods were compared.
- The non-linear methods tend to outperform the linear ones using a train and test procedure. Cluster-then-predict approaches often show the best performances, highlighting their usefulness.
- This methodology was then used to derive characterization factors that were missing for more than a hundred chemicals in USEtox^®^.

**Graphical abstract:** 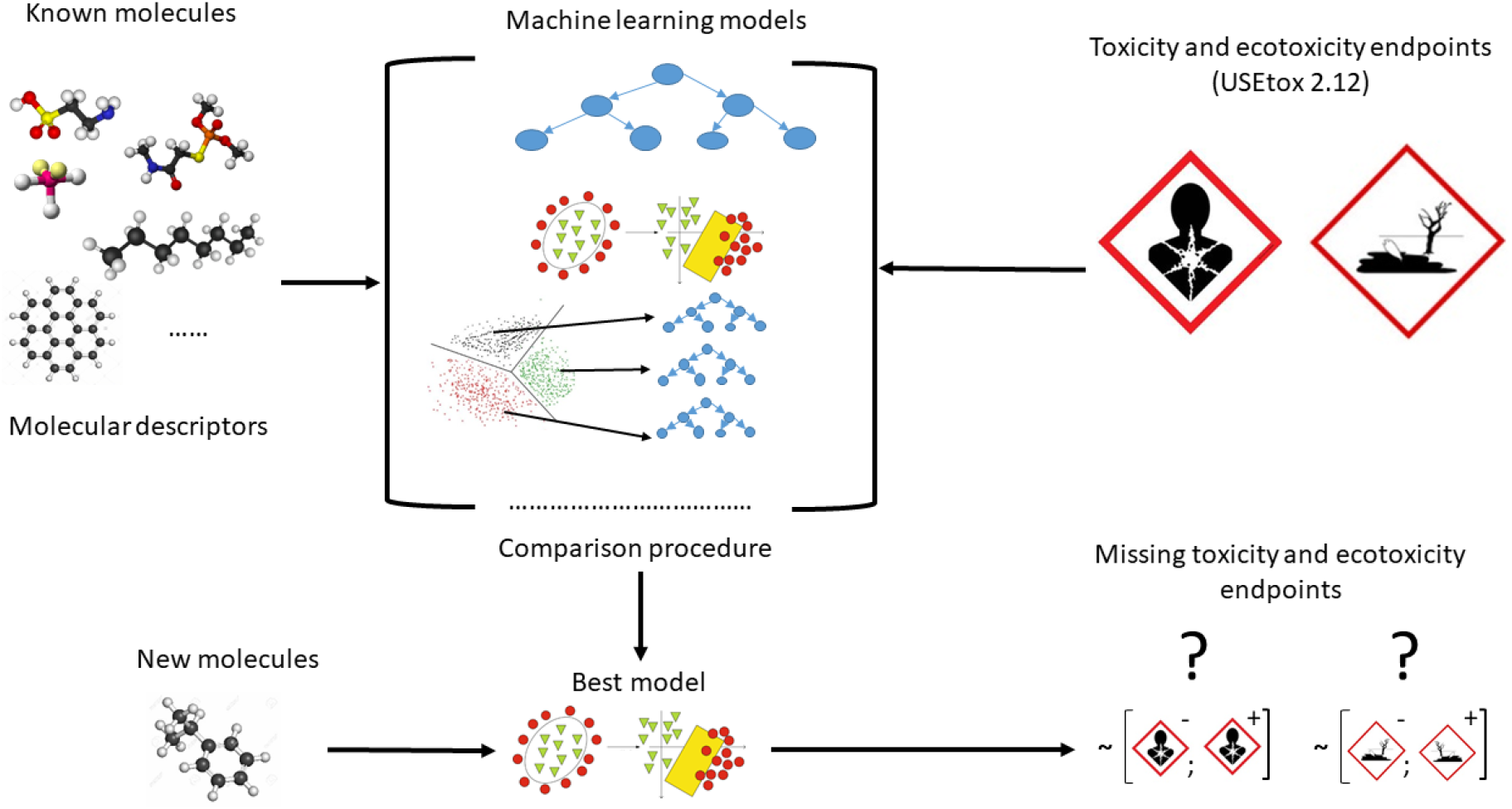

## Introduction

Recent legislations such as the Registration, Evaluation, Authorization and restriction of Chemicals (REACH) regulation in the EU requires that manufacturers of substances and formulators register to provide toxicological and ecotoxicological data for substances with volume higher than one metric ton per year. Furthermore, the U.S. Environmental Protection Agency (EPA) has more than 85,000 chemicals listed under the Toxic Substances Control Act (Hinds and Weller, 2016). Therefore, robust toxicological and ecotoxicological data are quickly needed to make informed decisions on how to regulate new chemicals. These data must also be coupled with environmental exposures and sources data, to better understand the impact on the environment.

To address the cause-effect relationships between the flow of molecules emitted by human activities and the consequences for ecosystems and humans, Life Cycle Assessment (LCA) offers a structured, operational, and standardized (Finkbeiner et al., 2006) methodological framework. Two main steps are at the core of this approach:

- Quantification of the masses of substances emitted into the environment through the Life Cycle Inventory (LCI). While it is possible to rely on databases that facilitate this inventory work for the background of the system under study, this task must nevertheless be carried out on a case-by-case basis to represent all the specificities of the foreground elements. This is the task of the LCA practitioner.
- Calculation of the impacts on ecosystems and human health of these emitted masses. Due to the complexity of environmental mechanisms, it is not possible to (re)model impact pathways on a case-by-case basis. Therefore, LCA uses characterization factors (CF) to assess the potential impacts of a compound. Concretely, if two compounds are emitted with the same mass, the one with the higher CFs will have the higher impact. Then, CFs are multiplied by the emitted masses of each compound to determine the impacts. CFs are not recalculated for each study but provided within a Life Cycle Impact Assessment (LCIA) method.

For a given impact, the LCIA method designer refers to the knowledge of the scientific community to model the mechanisms involved. For human toxicity and freshwater ecotoxicity, USEtox^®^ (Rosenbaum et al., 2008), was developed by life cycle initiative under the United Nations Environmental Programme (UNEP) and the Society for Environmental Toxicology and Chemistry (SETAC) (Henderson et al. 2011) to produce a transparent and consensus characterization model. USEtox^®^ is also used for the European Product Environmental Footprint (PEF) (Saouter et al., 2020). This model gathers in one single characterization factor the chemical fate, the exposure, and the effect for each of the several thousands of organic and inorganic compounds. Then, the USEtox^®^ model results can be extended to determine endpoint effects expressed as total (i.e. cancer and non-cancer) disability-adjusted life years (DALY) for human health impacts and potentially disappeared fraction of species (PDF) for ecotoxicological impacts. The PDF represents an increase in the fraction of species potentially disappearing as a consequence of emission in a compartment while the DALY represents an increase in adversely affected life years. These endpoints are now consensual at an international level (Verones et al., 2017).

If the structure of the USEtox^®^ multimedia model is always the same, to determine the CF of a molecule, numerous physicochemical parameters (such as solubility, hydrophobicity, degradability) and detailed toxicological and ecotoxicological data must be provided. For example, EC50 values (i.e. the effective concentration required to have a 50% effect) for at least three species from three different trophic levels are required for the ecotoxocological effect factor.

Over the past few decades, thousands of tests (in laboratory and field) have been carried out to evaluate the potential hazard effects of chemicals (He et al., 2017). Usually, toxicity testing has relied on *in vivo* animal models, which is extremely costly and time-consuming (Xia et al., 2008). In recent years, under societal pressures, there has been a significant paradigm shift in toxicity testing of chemicals from traditional *in vivo* tests to less expensive and higher throughput *in vitro* methods (National Research Council, 2007). However, it is still extremely difficult to test existing chemicals due to their large and ever increasing number, which leaves their impacts largely unknown. For example, in a recent study, Aemig et al. (2021) studied the potential impacts on Human health and aquatic environment of the release of 286 organic and inorganic micropollutants at the scale of France. One of their conclusion was that, due to a lack of characterization factors, these impacts could be assessed only for 1/3 of these molecules. That is why more computational models are needed to complement experimental approaches to decrease the experimental cost and determine the prioritization for those chemicals which may need further *in vivo* studies. Such models already exist, like QSAR models that are mostly linear models based on the chemical structure of compounds (Danish QSAR database (DTU, 2015), ECOSAR (Mayo-Bean et al., 2011), VEGA (Benfenati et al., 2013)) and are used to predict ecotoxicological data (LC50) needed for REACH for example. Recently, machine learning algorithms have been used to predict some midpoints based on molecular descriptors and environmental parameters (Marvuglia et al., 2014 and 2015; Song et al., 2017; Lysenko et al 2018) and a first review on this subject could be found in Wu and Wang (2018). After these first works, predictions of hazardous concentration 50% (HC50) based on 14 physicochemical characteristics (Hou et al., 2020a) or on 691 more various variables (Hou et al., 2020b) were carried out. Nevertheless, their input variables need some experiments and could be difficult to collect. This problem was tackled by Song et al. (2021) who predicted Lethal Concentration 50 (LC50) based on 2000 easy-to-obtain molecular descriptors. In the case of USEtox^®^, despite its wide use in LCA, it only offers characterization factors for approximately 3000 chemicals and even for this limited number of compounds, 19% of ecotoxicity CFs and 67% of human toxicity CFs are missing.

The objective of this article is thus to propose a new way of calculating CFs using machine learning approaches to solve the problem of nonlinearity that could affect a linear QSAR method. This makes it possible, when the CFs are not determined due to lack of time or lack of data, to propose values based solely on easily identifiable molecular descriptors. Here, the main differences with the above-cited methods are twofold: first, our input variables are only molecular descriptors that could be easily collected for any newly available compounds; second, our output variables are directly the CFs that are closer to the endpoints (DALY and PDF) than the HC50 or the LC50 (i.e. the acute aquatic toxicity experimental threshold). These two specific endpoints will be studied in the present paper through the emission of compounds in continental freshwater and will be named CF_ET_ for ecotoxicological impacts and CF_HT_ for human ones. To address this aim, we will test different methods (linear and non-linear) and assess their performances, to build a robust model that could predict CFs that are currently lacking.

## Materials & Methods

### USEtox^®^ database

The last version of the USEtox^®^ database was downloaded, namely the corrective release 2.12 (USEtox^®^, 2020). The whole USEtox^®^ 2.12 database contains 3076 compounds. The CFs were computed using the default landscape.

### TyPol database

We recently developed TyPol (Typology of Pollutants), a classification method based on statistical analyses combining several environmental parameters (i.e., sorption coefficient, degradation half-life, Henry constant) and an ecotoxicological parameter (bioconcentration factor BCF), and structural molecular descriptors (i.e., number of atoms in the molecule, molecular surface, dipole moment, energy of orbitals). Molecular descriptors are calculated using an *in silico* approach (combining Austin Model1 and Dragon software). In the present paper, we only extracted and used the molecular descriptors from the TyPol database, as this information could be easily collected for any new compound. The 40 descriptors included in the TyPol database have been selected based on a literature review on QSAR equations used to predict the main environmental processes as degradation, sorption, volatilization. These 40 descriptors were the ones most frequently used in the equations, meaning describing the best the behaviour of organic compounds in the environment. By consequence, even if no environmental parameters were directly incorporated as input in our model, some information that is directly linked to them were included in the 40 molecular descriptors. These descriptors are constitutional, geometric, topological, and quantum-chemical descriptors (see Table 1); 35 described the 2D-structure of the compound while the other five are linked to its 3D-structure. An important advantage of the unique use of molecular descriptors is that they are easily and quickly computable for not yet synthesized compounds. For more details, we refer the interested reader to Servien et al. (2014) and to Mamy et al. (2015) where the choice of the 40 molecular descriptors is described in details. Now, TyPol gathers 526 compounds, including pesticides, persistent chemicals, pharmaceuticals and their transformation products (Benoit et al. 2017, Traoré et al. 2018).

**Table 1.** List of the 40 molecular descriptors in TyPol

| Category | Molecular descriptors |  |  |
| --- | --- | --- | --- |
| Constitutional | Number of atoms | Number of non-H atoms | Number of hydrogen atoms |
|  | Number of chlorine atoms | Number of carbon atoms | Number of nitrogen atoms |
|  | Number of oxygen atoms | Number of phosphorus atoms | Number of sulfur atoms |
|  | Number of fluorine atoms | Number of circuits | Number of halogen atoms |
|  | Number of bonds | Number of non-H bonds | Number of double bonds |
|  | Number of triple bonds | Number of multiple bonds | Number of rotatable bonds |
|  | Number of aromatic bonds | Sum of conventional bond order | Number of rings |
|  | Molecular weight |  |  |
| Geometric | Connolly molecular surface area |  |  |
| Topological | Connectivity index of order 0 | Connectivity index of order 1 | Connectivity index of order 2 |
|  | Connectivity index of order 3 | Connectivity index of order 4 | Connectivity index of order 5 |
|  | Valence connectivity index of order 0 | Valence connectivity index of order 1 | Valence connectivity index of order 2 |
|  | Valence connectivity index of order 3 | Valence connectivity index of order 4 | Valence connectivity index of order 5 |
| Quantum-chemical | Polarizability | Electric dipole moment | HOMO energy |
|  | LUMO energy | Total energy |  |

### Machine learning methods

To predict the CFs using the molecular descriptors we used three modeling methods combined. The first method is a linear well-known prediction method namely the Partial Least Squares (PLS) (Wold, 1985). It finds the multidimensional directions in the observable variable (molecular descriptor) space that explains the maximum multidimensional variance direction in the predicted variable (CF) space. That provides a linear regression model based on the observable variables to predict the predicted variable. We also chose to compare two machine learning methods adapted to non-linear problems: the random forest (Breiman 2001) and the support vector machines (SVM) (Drucker et al. 1996). Random forests are a machine learning method, for classification or, in our case, regression, that operate by constructing a multitude of decision trees that uses a random subset of the training data and limits the number of variables used at each split and outputting the mean prediction (regression) of the individual trees. SVM constructs a hyperplane or set of hyperplanes in a high- or infinite-dimensional space in which the problem is linearly separable.

These choices allowed us to compare several ideas. The PLS is a simple linear method that will not exhibit good performances if the underlying relationship is not linear. The SVM and RF methods are well-known non-linear machine learning algorithms that used to show good results in this kind of problem (Hou et al., 2020a).

All the models were computed in the freeware R (R core team, 2019). The PLS has been computed using the package mixOmics (Rohart et al., 2017), the random forests using the package randomForest (Liaw et al., 2002), and the SVM using the package e1071 (Meyer et al., 2019). These 3 modeling methods have some parameters that needed to be fixed: the number of latent components for the PLS (fixed using the tune.pls function), the number of variables randomly sampled as candidates at each split for the random forests (selected using the tune.randomForest function) and, for the SVM, the gamma parameter of the radial kernel and the cost of constraints violation (using the tune.svm function). All these different tune functions are based on cross-validation (i.e. a training/test procedure to find the best value for the parameters) using default function values.

### Clustering-based model

A recent popular way to make predictions is to use a cluster-then-predict approach. That is, clustering is used for pre-classification which is to arrange a given collection of input patterns into natural meaningful clusters. Then, the clustering results are used to construct a predictor in each cluster. The main idea of the cluster-then-predict approach is that if the clustering performs well, the prediction will be easier by modeling only similar compounds. If a new compound with no CF_ET_ and/or CF_HT_ is investigated, the clustering can easily be applied to it before the prediction model itself. The cluster-then-predict approach has already been applied with success in various domains such as sentiment prediction (Sony et al., 2015), finance (Tsai et al., 2014), chemometrics (Minh Maï Le et al., 2018). So we decided to use the clustering given by the TyPol application (more details in Servien et al., 2014). Note that the TyPol clustering has already been shown relevant on various occasion: in combination with mass spectrometry to categorize tebuconazole products in soil (Storck et al., 2016), to explore the potential environmental behaviour of putative chlordecone transformation products (Benoit et al., 2017) or to classify pesticides with similar environmental behaviors (Traore et al., 2018; Mamy et al., 2021). So, the clustering procedure of TyPol was applied on the whole database of 526 compounds using the 40 molecular descriptors. This approach provided us a global clustering based on all the available information contained in the TyPol database. It is based on PLS, hierarchical clustering and an optimal choice of the number of clusters and is detailed in Servien et al. (2014). The obtained clustering is given in Supplementary Figure S1 and relies on 5 different clusters. The Supplementary Figure S2 represents this clustering restricted to the common molecules between TyPol and USEtox^®^. We could see that, as the cluster 5 is only constituted of one compound, the cluster-then-predict-models cannot be applied.

Based on this clustering, we then defined three other competing methods. For these methods, a different model (with different parameters) was derived for the compounds in each cluster. Consequently, six different models were calibrated and tested for each CF prediction: global PLS, global SVM, global random forest, cluster-then-PLS, cluster-then-SVM and cluster-then-Random Forest.

### Comparison procedure of the models

To assess the performances of the different models we used the following procedure:

1. Split each cluster (the whole dataset if the model is global, only the data lying in the dedicated cluster if that is a cluster-then-predict model) between a training set (85% of the dataset) and a test set (15%) (Pareto principle). The test set is not used for any step of the procedure (such as the imputation of the missing data, the calibration of the parameters…).
2. Imputation of the NA (Not Available, *i.e.* missing) values (less than 1%) in the descriptor matrix using the NIPALS algorithm (Wold, 1985).
3. Tune the parameters and train the specific models by performing cross-validation on the training set. We have 3 global models to train (PLS, random forest, and SVM) and the cluster-then-test models (PLS, random forest and SVM for each cluster).
4. Test the different models on the test set. Compute the absolute error.
5. Back to step 1.

The whole algorithm was repeated 200 times. All the performances were compared in terms of absolute error. The absolute error is the absolute difference between the prediction and the true value. It has been shown to be the most natural and unambiguous measure of error (Willmott et Matsuura, 2005) and is chosen to be easily comparable to the assumed error on the experimental CFs (2-3 logs, see Rosenbaum, 2008). For each cluster, we chose the model with the lowest median absolute error.

### Predictions

Then, the best model was calibrated and computed on the whole cluster. Finally, it was applied to the compounds, according to their clusters, with a CF_ET_ (or a CF_HT_) equals to NA to provide a prediction. For the compounds in cluster 5, this best model cannot be a cluster-then-predict one and, by consequence, is a global one. To assess the robustness of our prediction we derived a 95% prediction interval for each prediction. The type of model and its corresponding parameters were fixed during this process, according to the best model of the cluster. For example, if the best model of cluster 1 was the random forest approach, random forest models are used with the parameters optimized during the previous step. Then, we performed a leave-one-out bootstrap on the dataset that was used to compute the model (the whole dataset if the model is global, only the data lying in the dedicated cluster if that is a cluster-then-predict model) and a new model was computed on this leave-one-out sample. A prediction was carried for each leave-one-out model (*i.e*. n-1 models if n is the number of compounds of the, eventually global, cluster) and the 2.5% and 97.5% quantile of these predictions were computed and considered as the prediction interval (Hou et al., 2020a). The whole modeling process is summarized in the following Figure 1.

**Figure 1.**
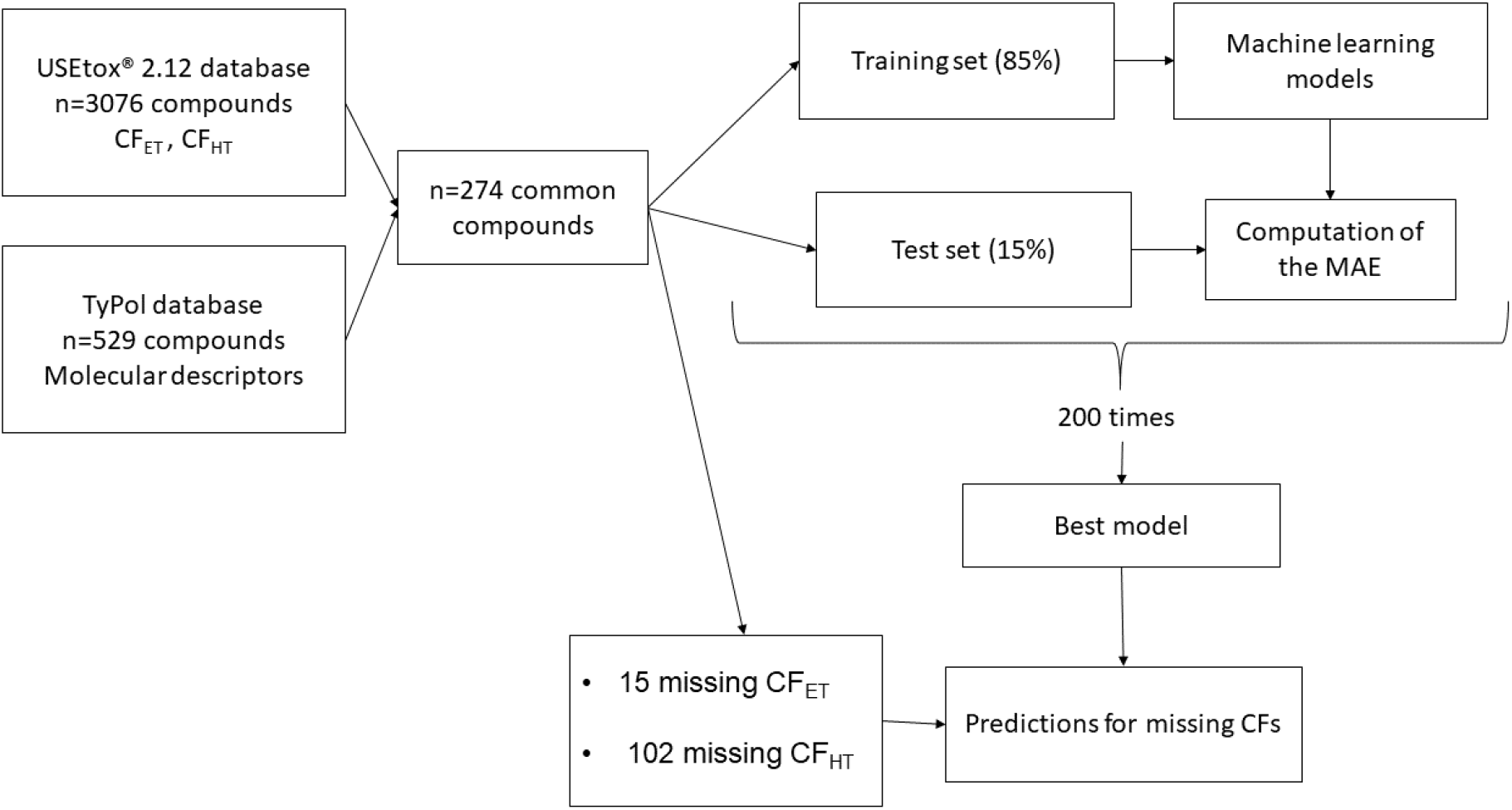
Schematic representation of the modeling procedure adopted in the paper.

The five molecular descriptors contributing the most to the prediction were then derived for each chosen model to assess the differences between models and to interpret their relevance. For a random forest model, these descriptors are calculated using variable permutations (Breiman, 2001), for the SVM they are the descriptors with the higher coefficients in absolute value.

## Results

### Descriptive analysis of the intersection of the TyPol and the USEtox^®^ databases

As the objective of this proof-of-concept study was to predict USEtox^®^ CF_ET_ and CF_HT_ using the molecular descriptors contained in TyPol, we could only use the compounds that are present in both databases. This resulted in 274 compounds that are detailed in Table S1 in supplementary material and the range of their CF_ET_ and CF_HT_ values are summarized in the boxplots in Figures 2 and S3. Note that for the 274 common compounds there are 15 NA values for the CF_ET_ and 102 for the CF_HT_.

**Figure 2.**
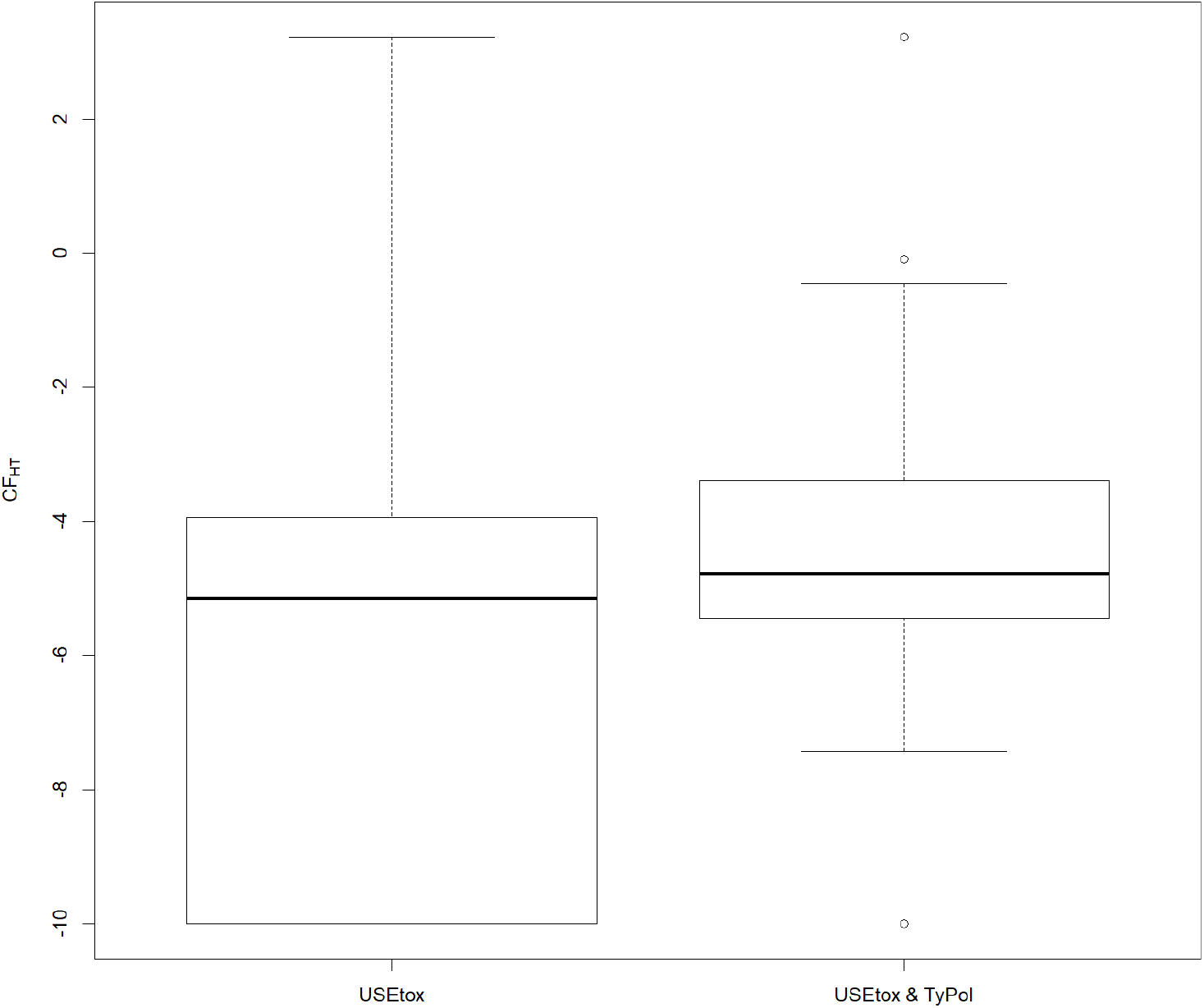
Boxplots of the CF_HT_ for the USEtox^®^ database and the common molecules between the USEtox^®^ and the TyPol databases. This CF_HT_ is equal to log_10_((DALY+ε).kg^-1^). The ε is needed as some values of the DALY are exactly equal to zero. ε has been chosen equal to 1e-10 to be below the minimum of the USEtox^®^ database (5e-9).

We could see on these two figures that the common compounds present higher CF_ET_ and CF_HT_ values than the one of the complete USEtox^®^ database: it focuses on the more dangerous compounds as their boxplots are above the USEtox^®^ counterparts and they cover the whole order of magnitude of the CFs of the USEtox^®^ database.

### Clustering of the compounds

The global Typol clustering of Supplementary Figure S1 with only the common compounds is plotted in Supplementary Figure S2 and the boxplots of each molecular descriptor per cluster are given in Supplementary Figure S4 with different indicators in Table S2. We could see that they are clustered in 5 groups with different sizes (respectively 33 compounds in the first black cluster, 122 compounds in the second red cluster, 91 compounds in the third green cluster, 27 compounds in the fourth blue cluster, and one compound in the fifth brown cluster). Cluster 1 grouped compounds with a high number of aromatic bonds, double bonds, rotatable bonds, and multiple bonds. Cluster 2 is an intermediate one between clusters 1 and 3, with less extreme values. Cluster 3 is made of compounds with the lowest molecular mass. Cluster 4 gathered compounds presenting a high number of halogens, rings, and circuits. The unique compound in the fifth cluster is erythromycin (highest molecular mass and number of H and C, lowest number of rings) and, obviously, no cluster-then-predict model could be built for this cluster

As a first analysis of the clustering given by TyPol, we could see in Figure 3 below the boxplots of the CF_ET_ and CF_HT_ within the 5 clusters.

**Figure 3.**
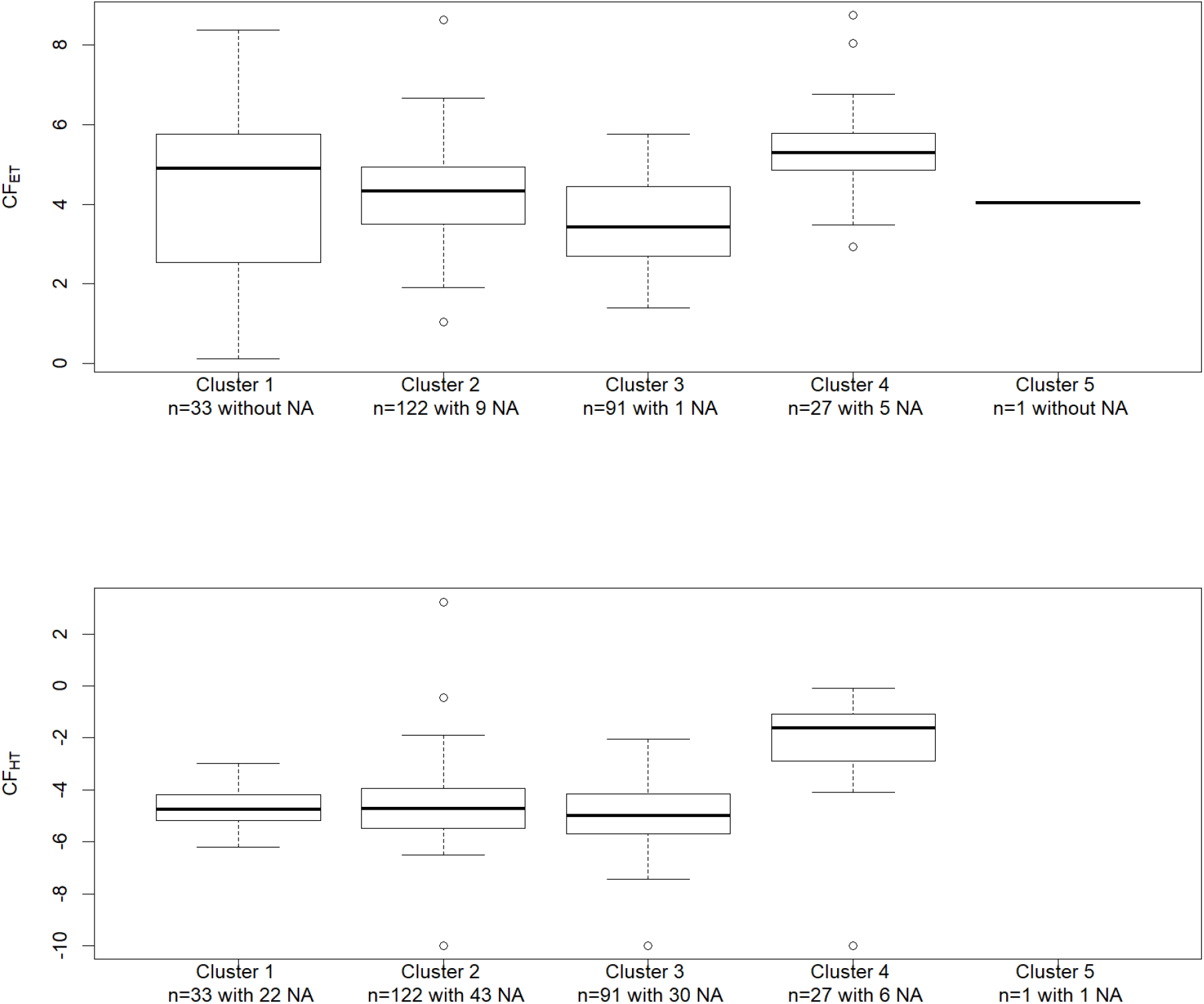
Boxplot by cluster for the CF_ET_ and CF_HT_ values. Note that the unique compound of Cluster 5 has no CF_HT_ value. The size of the clusters and the numbers of NA are gathered in the legend.

The predictions will be made difficult for the CF_ET_ of cluster 1 as it covers a wide range whereas it includes a relatively small number of compounds. On the contrary, cluster 3 covers a small range with no extreme values and includes a high number of compounds, for this cluster the cluster-then-predict approach could produce interesting results.

### Performances of the machine learning methods

#### For the prediction of the CF_ET_

The methodology described in the previous section was applied to our dataset and gave the results gathered in Figure 4 for each cluster.

**Figure 4.**
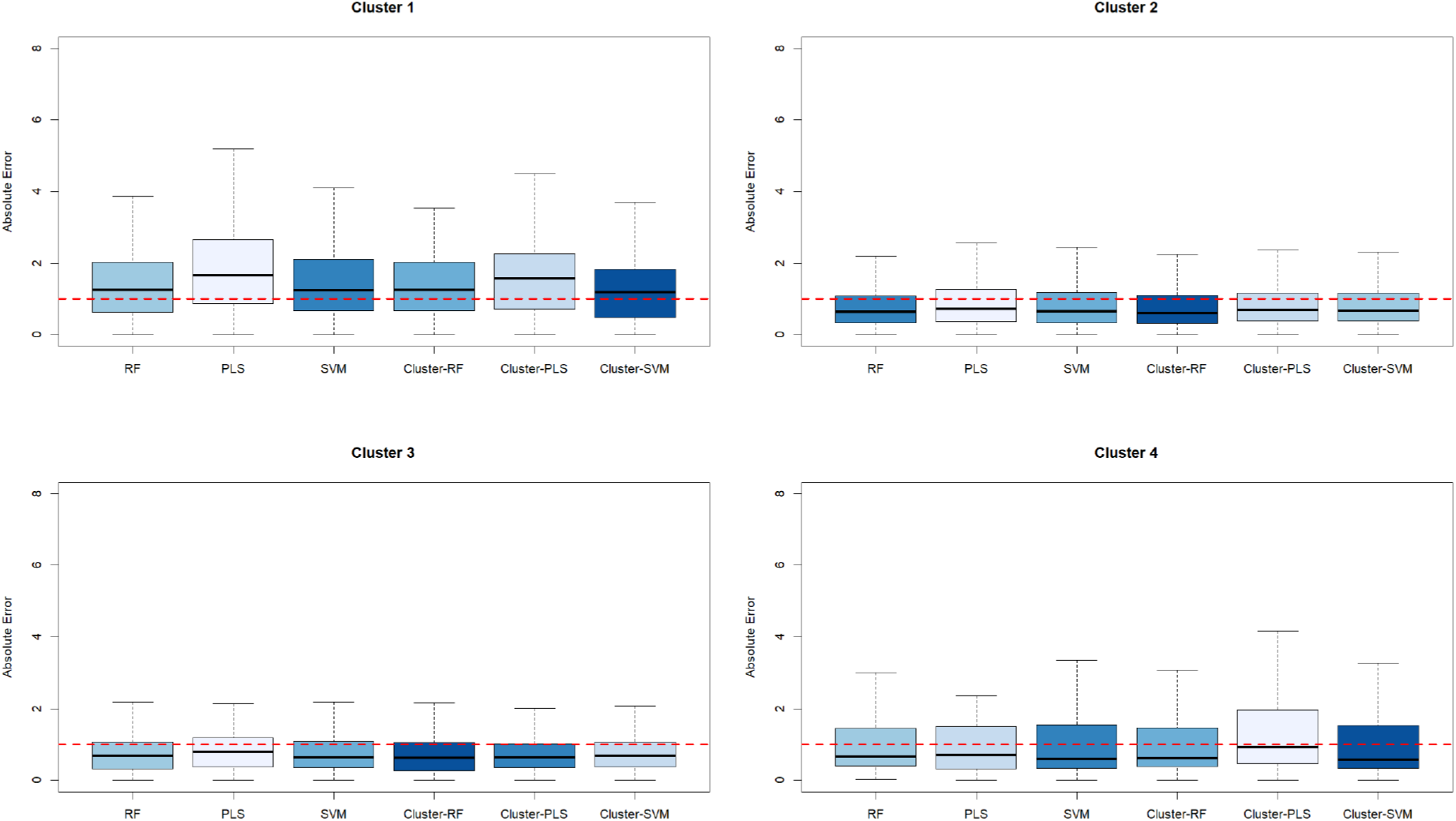
Performances of the different methods (RF: random forest, PLS: partial least squares, SVM: support vector machines) in terms of the log of the absolute error of the CF_ET_ with respect to the different clusters for 200 repetitions. In each cluster, the models are coloured from dark blue (best) to clear blue (worst) according to their median of the absolute error. The red dotted line represents an absolute error of 1 log that is considered as acceptable (Rosebaum et al. 2008; Douziech et al., 2019).

The performances were not similar from one cluster to another. For example, performances of all methods for cluster 1 were very poor (median absolute error above 1) whereas performances for cluster 4 seemed good despite its smallest size (median absolute error around 0.6). Therefore, a future prediction of an unknown compound which lies in cluster 1 will be less reliable than in other clusters. Note that we could not test this in the next section as no NA value is present in this cluster 1.

The cluster-then-predict methods seemed more appropriate in each cluster. The cluster-then-RF approach had the best performances (with a global median absolute error equals to 0.64 and the best performances on clusters 2 and 3), even if there was not a big difference between the different methods. The cluster-then-SVM was also the best method for the two clusters 1 and 4. The linear methods (PLS and cluster-then-PLS) had higher absolute errors but were competitive. The individual predictions of the best method in each cluster are reported in Figure S5.

#### Prediction for the CF_HT_

Let us recall that we have more NA values for the CF_HT_ (102) than for the CF_ET_ (15). The performances of the methods are illustrated in the following figure.

**Figure 5.**
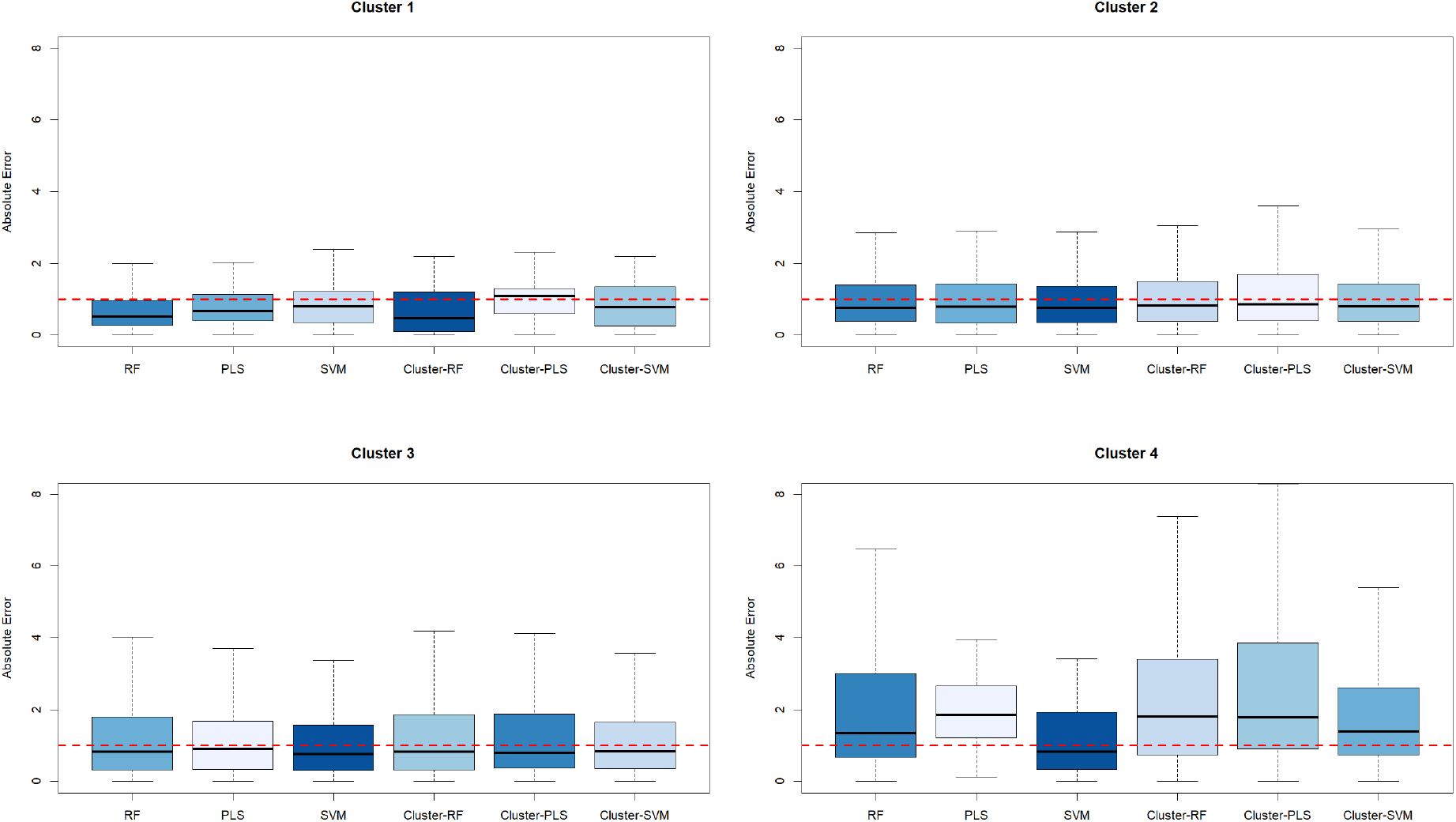
Performances of the different methods (RF: random forest, PLS: partial least squares, SVM: support vector machines) in terms of the log of the absolute error of the CF_HT_ with respect to the different clusters for 200 repetitions. In each cluster, the models are coloured from dark blue (best) to clear blue (worst) according to their median of the absolute error. The red dotted line represents an absolute error of 1 log that is considered as acceptable (Rosebaum et al. 2008; Douziech et al., 2019).

We observed that, despite its small size (11 compounds), the CF_HT_ of the first cluster were well predicted (with the best performance for the cluster-then-RF approach). It could be explained by the small range of the CF_HT_ values of this cluster, as illustrated on the boxplot in Figure 3. The performances of all the methods were comparable on clusters 2 and 3 where the best method was the SVM. Cluster 4 seemed to be the most difficult to predict: all the methods had their worst results on this cluster and, if the SVM had an acceptable median absolute error of 0.82, all the medians of the other methods were above 1.3. Global performances of the different methods were given in Supplementary Figure S6. Note that, as for CF_ET_, the linear methods based on PLS were outperformed by the other ones.

### Best model predictions

#### Best models for the CF_ET_

Using our methodology, we could exhibit a median absolute error of 0.62 log for the prediction of the CF_ET_ on the whole dataset using the best models. If we looked closer on Table S3, we could see that the median of our estimations is below 0.6 log except for cluster 1 (above 1 log).

Then we calibrated the best models on the whole dataset of each cluster: a cluster-then-predict approach using SVM for clusters 1 and 4 and using random forest for clusters 2 and 3. To compare the different models in each cluster and give an idea of what were the important molecular descriptors we provided the five most important molecular descriptors for each cluster in the Table S4. We could see in this table that the important molecular descriptors strongly differ from one cluster to another.

Then the models were used to predict the missing CF_ET_ of the common compounds between USEtox^®^ and TyPol databases. These values were by consequence new estimations of the CF_ET_ for compounds on which we had no information. The prediction intervals were relatively small: less than 0.5 log10 in a log scale which highlighted the robustness of the estimation. They are given in Table S5. No NA value was present in cluster 1 with no prediction for this cluster. For cluster 2 gathering molecules with intermediate molecular mass, 9 CF_ET_ values were predicted for various kinds of compounds. One value concerns the antibiotic sulfamethazine and its value is quite near to the one of sulfamethoxazole and sulfadiazine of the same sulphonamide antibiotic family constituted of the sulphonamide group (−S(=O)_2_-NR_2_R_3_). Cluster 3 grouped compounds with the lowest molecular mass and the lowest median CF_ET_ like ibuprofen, phthalates, cresol constituted of monoaromatic ring substituted with methyl, carboxylic groups. The CF_ET_ prediction for acetylsalicylic acid seemed coherent with the value of the nearest compounds (herbicides mecoprop) of this group. Cluster 4 gathered compounds with the highest median CF_ET_ and that presented a high number of rings halogenated or not, like PAH and hormones. The 5 CF_ET_ predicted concerned 4 PAHs and 1 hormone. By comparison to the 2 other PAHs present in this cluster, the 4 predicted CF_ET_ were quite similar and higher. Concerning the prediction for the hormone, the CF_ET_ was intermediate between the CF_ET_ of the 3 other hormones in the cluster. It seems that all these 5 predicted values were very closed, falling near the median value of this cluster.

#### Best models for the CF_HT_

The best models had a global median of 0.75 log for the prediction of the CF_HT_. We could see on the results gathered on Table S6 that the best performances are for cluster 1 and that they are comparable for the other clusters.

Then, the global SVM model was calibrated and computed on the whole dataset. It was used to predict the compound of clusters 2, 3, 4, and 5. Let us recall that there was a single molecule in cluster 5 and, as it has a NA value for its CF_HT_, the best global model (SVM) was used. For cluster 1, a cluster-then-RF model was computed. The most important descriptors of these two models are gathered in the Table S7 and, as for CF_ET_, were strongly different between the different best models.

Then, this model was used to predict the CF_HT_ value for the 102 common compounds without a CF_HT_ value. These predictions are reported in Supplementary Table S8. As for the CF_ET_, the small width of the prediction interval (less than a log10 in a log scale) highlighted the robustness of the approach even with a relatively small number like estimations made for compounds that lie in cluster 1. In this cluster 1, CF_HT_ for a phthalate (DEHP) was already known, but the one for diisodecyl and diisononyl phthalate was predicted with value in the same range. The 3 cyclines (tetracycline, aureomycin, and oxytetracycline) grouped in cluster 1, presented also similar predicted CF_HT_. This was also the case for triclosan and triclocarban in cluster 2. Similar predicted and known CF_HT_ were found for four herbicides from the substituted urea family (linuron, diuron, monolinuron, isoproturon) in cluster 3. Cluster 4 gathered a small number of molecules but with the highest median CF_HT_, the predicted CF_HT_ of the organochlorine insecticide isodrin was similar to another congener of the same family, aldrin.

## Discussion

It is a real and important challenge to provide characterization factors for a wide range of compounds. Obviously, it is expected that these new calculated factors have an acceptable margin of error. As reported in UNEP/SETAC (2019), it is commonly assumed that the uncertainty of the characterization factors can vary by approximately 2-3 orders of log-magnitude (Rosenbaum et al. 2008) or significantly higher (up to 7 orders) if all sources of uncertainty are considered (Douziech et al. 2019). The results obtained in the previous section are very promising as they are below the level of uncertainty commonly assumed and as they are based on molecular descriptors that could be easily obtained for each compound without ecotoxicity factor. Based on this fact we could already provide 15 new CF_ET_ and 102 new CF_HT_ for the common molecules between USEtox^®^ and TyPol without a previous value.

The idea of predicting ecotoxicity characterization factors for chemicals using machine learning algorithms has already been used (Hou et al., 2020a and 2020b). But, here, our findings go further. Indeed, we show that we could directly obtain accurate estimations of endpoint values from easy-to-obtain molecular descriptors. This will open the door to the fast characterization of each new unknown compound that appears, including transformation products. We also show that the cluster-then-predict approach can give better performances than the approach without the clustering step. This local (*i.e*. cluster-then-predict) approach confirms that local models could be an efficient prediction method when heterogeneity of data generates nonlinear relations between the response and the explanatory variables (Lesnoff et al., 2020).

Across the clusters and models, there is a general trend that the non-linear models tend to outperform the linear ones. This suggests that a linear model is not fully adequate to capture the complexity of the relationship between the molecular descriptors and the CFs. However, the use of linear model for e.g. a QSAR is likely due to the ease of interpreting its coefficients, while interpretation is much more challenging for machine learning approaches such as random forest or SVM. Thus, the advantages or drawbacks of linear/non-linear approaches must be balanced according to the final goal of each study. Here, as the main goal is to calculate the most accurate CFs, non-linear models seem more suited. We must also mention that a new emerging field is developing tools needed to help making black-box models (e.g. random forest) more interpretable (Bénard et al., 2021).

The difficult interpretability of the machine learning models used in this study can thus be viewed as a limitation. On another side, even if we already had an acceptable number of compounds in our training datasets, the model accuracies would benefit of the inclusion of new compounds. These compounds could be carefully chosen to improve the models where there is a clear need (i.e. where the performances of the models are not good enough), for example in the cluster 1 for CF_ET_ or in the cluster 4 for CF_HT_.

One of the interests of USEtox^®^ and its three-step structure (fate - exposure - effect) is that it can be adapted to some specific contexts (a more accurate and spatialized fate model, a different exposure…) while keeping the steps that are not modified. However, these adaptations of USEtox^®^ are not widely used and are reserved for advanced users. Our approach does not allow this, with a direct one-step estimation of CFs. It was designed to provide default CF values for molecules where information is missing. We have chosen to directly predict the CF by simplicity, as the first tests revealed that doing three models (for the three steps) and then calculating the CFs produced less accurate results. It would however be an interesting perspective to estimate only some of the stages by these learning approaches and to combine them with stages modelled in a classical way in USEtox^®^

## Conclusion

This paper presents a modeling method to derive characterization factors from easily obtainable molecular descriptors. The results presented here show that models that can handle non-linearity and that could be adapted to a small number of compounds (using the cluster-then-predict approaches) are the best suited. The cluster-then-predict approaches could also be more accepted by the users, as they allow to consider mainly the compounds similar to the one under investigation. By consequence, the missing characterization factors, as well as those of new molecules, could now be quickly estimated with an overall good precision as performed in Servien et al. (2021). More generally, one of the key factors in the evaluation of toxicity and ecotoxicity in LCA lies in the construction of the characterization factors: a task requiring a large amount of data and a consequent investment of time. The use of machine learning allows us to go beyond these constraints and to propose a new methodology in the LCA framework. This makes it possible to obtain characterization factor values in a fast and simple way, which can be used as long as conventionally established CFs are not available.

## Supporting information

Scripts and data

## Acknowledgements

The authors are grateful to Pierre Benoit, Laure Mamy, and Virginie Rossard for their work on TyPol. Version 6 of this preprint has been peer-reviewed and recommended by Peer Community In Ecotoxicology and Environmental Chemistry (https://doi.org/10.24072/pci.ecotoxenvchem.100001).

## Supplementary materials

Supplementary material associated with this article can be found, in the online version. Scripts and data used are provided.

## Data, script and code availability

Data, script and code are provided as supplementary materials of this preprint (https://doi.org/10.1101/2021.07.20.453034)

