## Supplemental Material for "Machine learning models based on molecular descriptors to predict human and environmental toxicological factors in continental freshwater"

### 1. Supplemental Figures

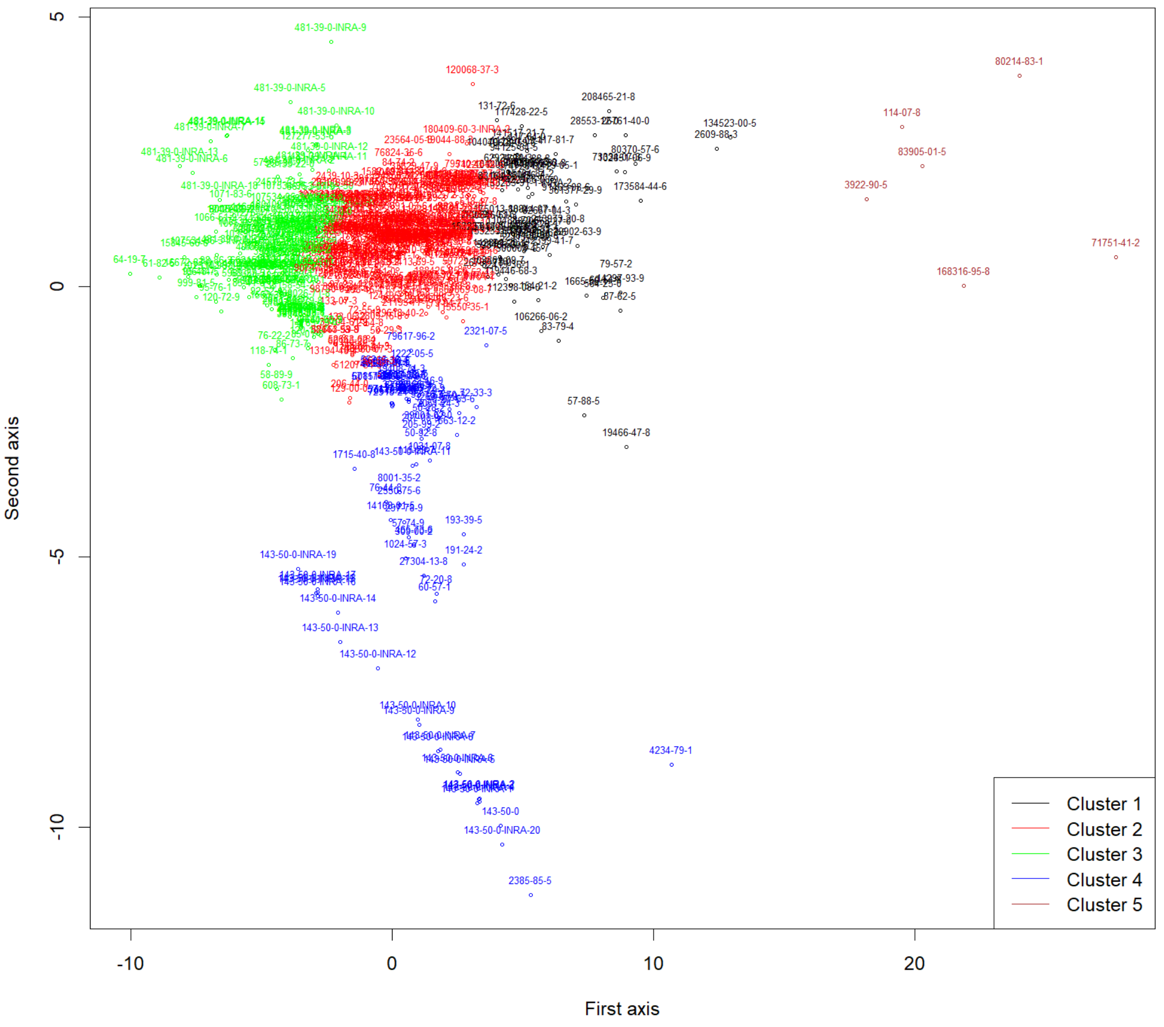


**Figure S1**- Clustering based on molecular descriptors produced by TyPol on the 526 molecules of the database. We represent here the two first axes of the PLS and the five different clusters in different colours.

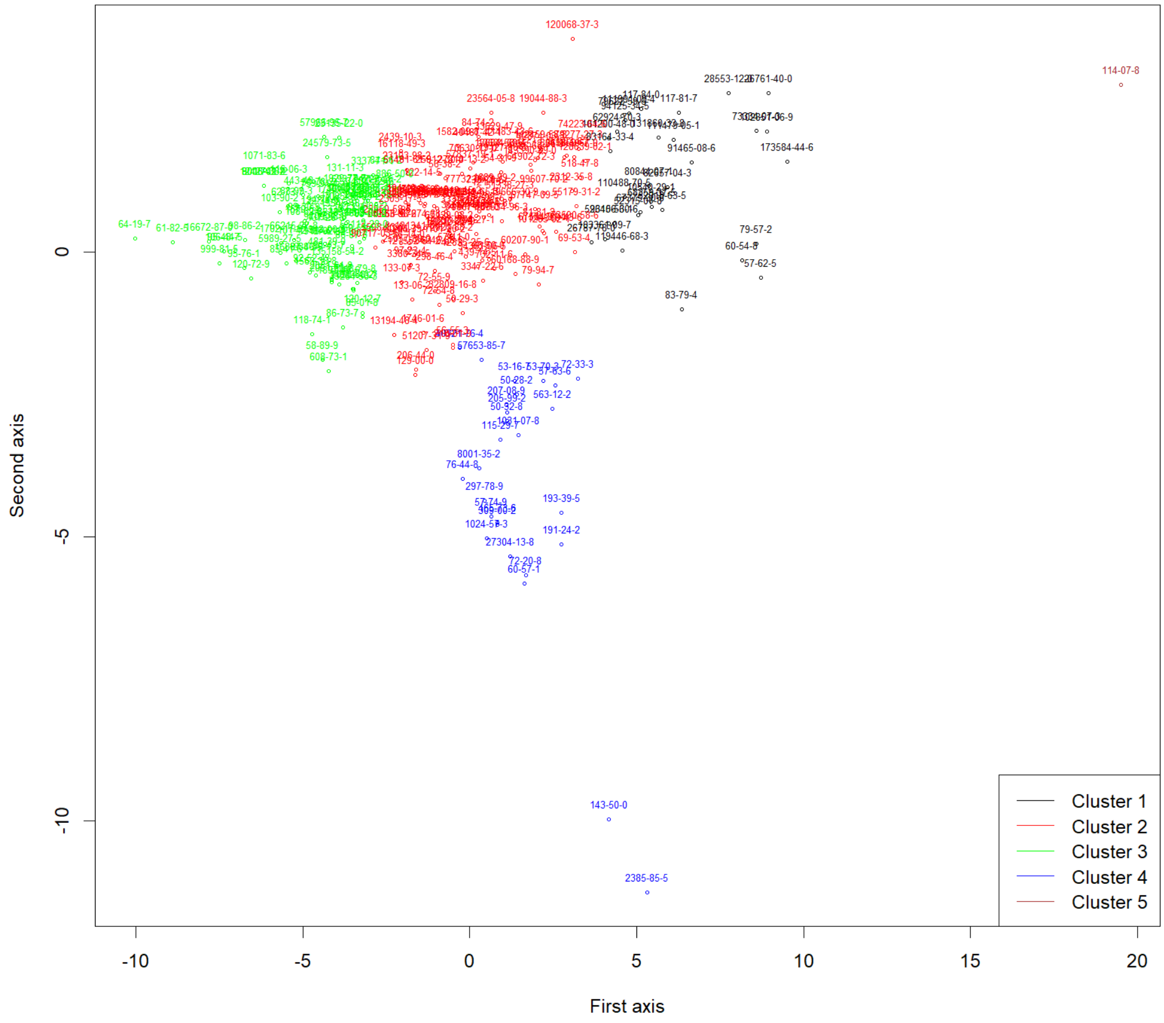


**Figure S2**- Focus on the 274 common molecules of TyPol & USEtox. The cluster 5 in brown is reduced to a single molecule so the cluster-then-predict methodology cannot be applied for it.

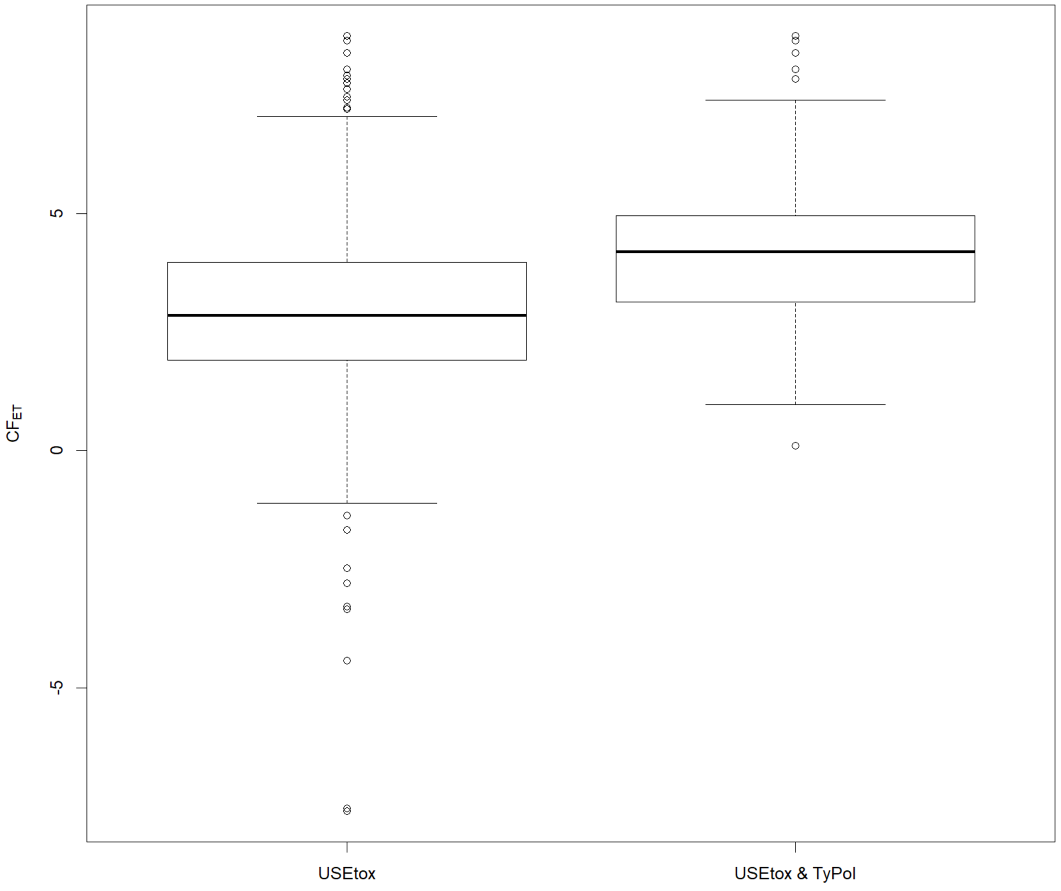


**Figure S3**- Boxplots of the $CF_{ET}$ for the USEtox® database and the common molecules between the USEtox® and the TyPol databases. This $CF_{ET}$ is equal to the $log_{10}(PDF.m^3.d.kg^{-1})$.

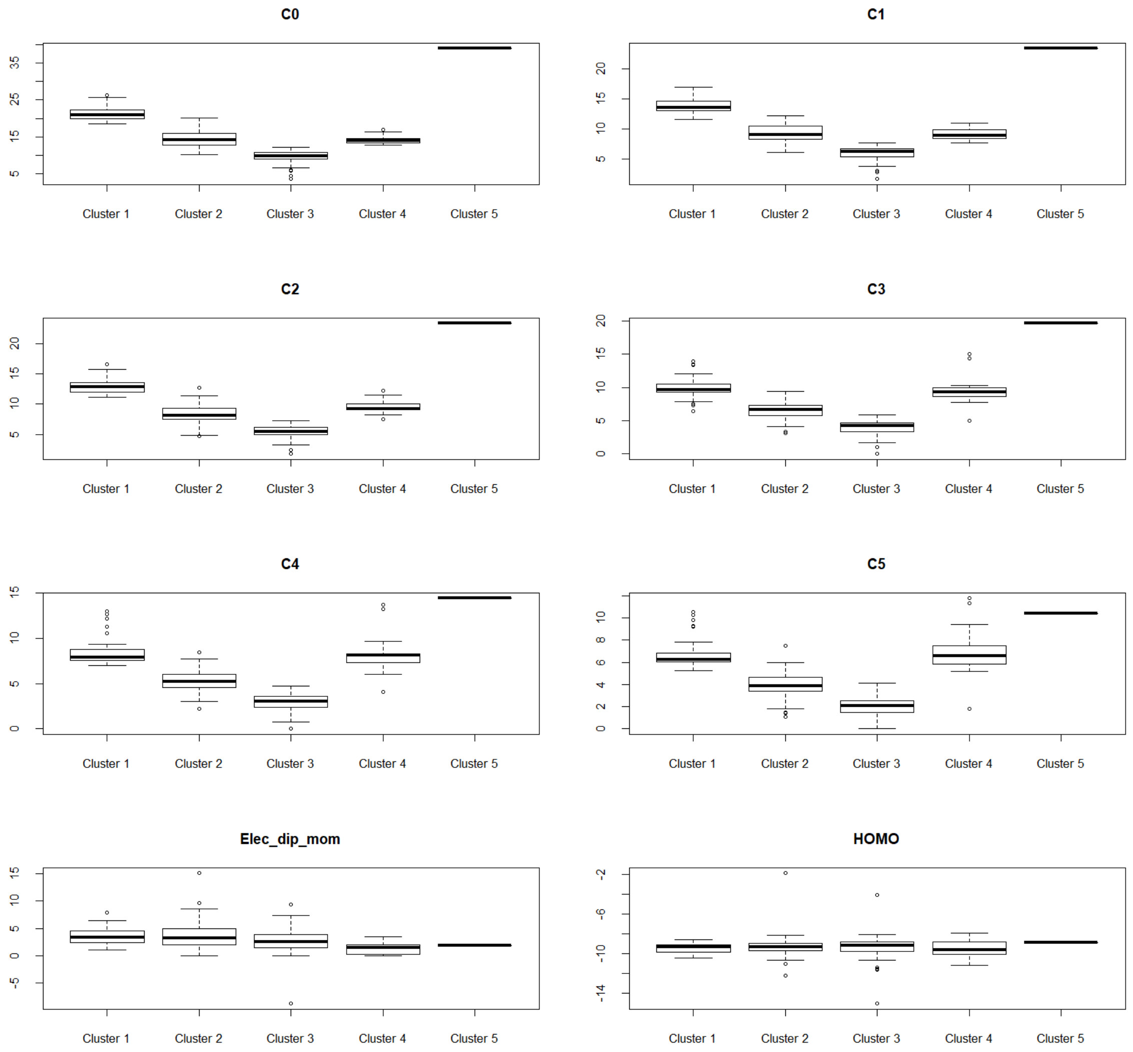


**Figure S4** – Boxplots of the 40 molecular descriptors for the clustering given by TyPol on the common compounds of TyPol & USEtox.

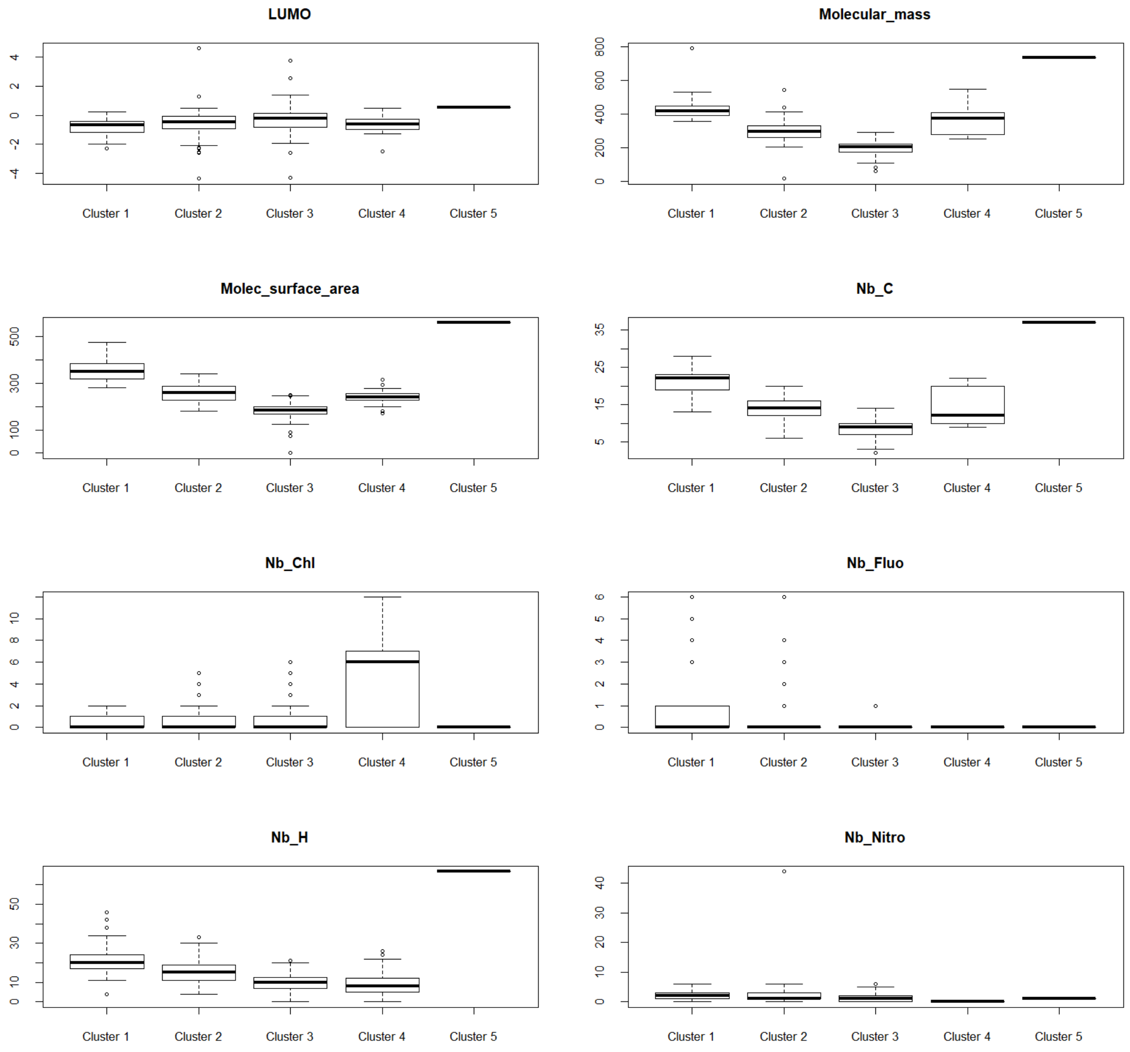


**Figure S4** (continued)

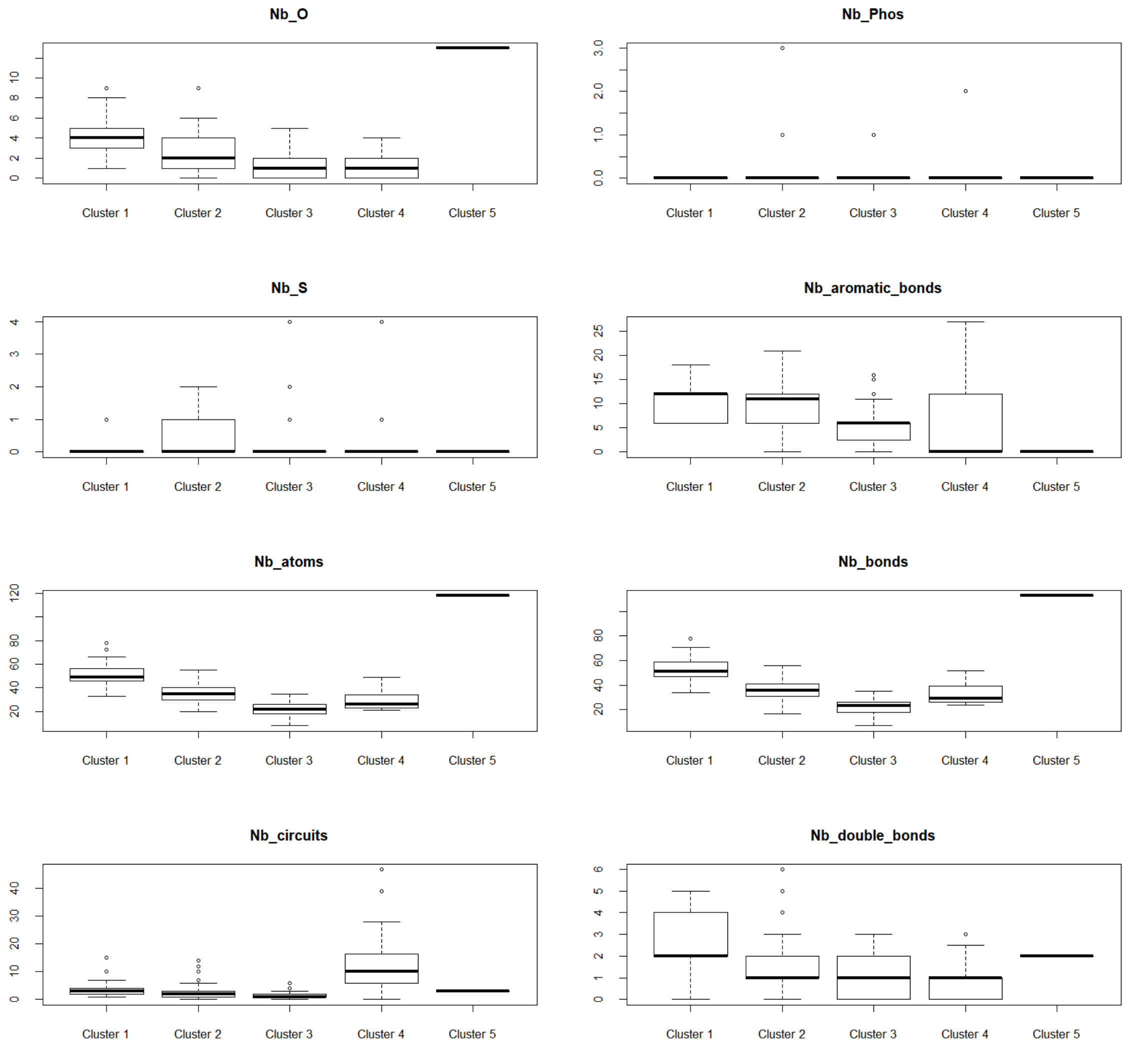


**Figure S4** (continued)

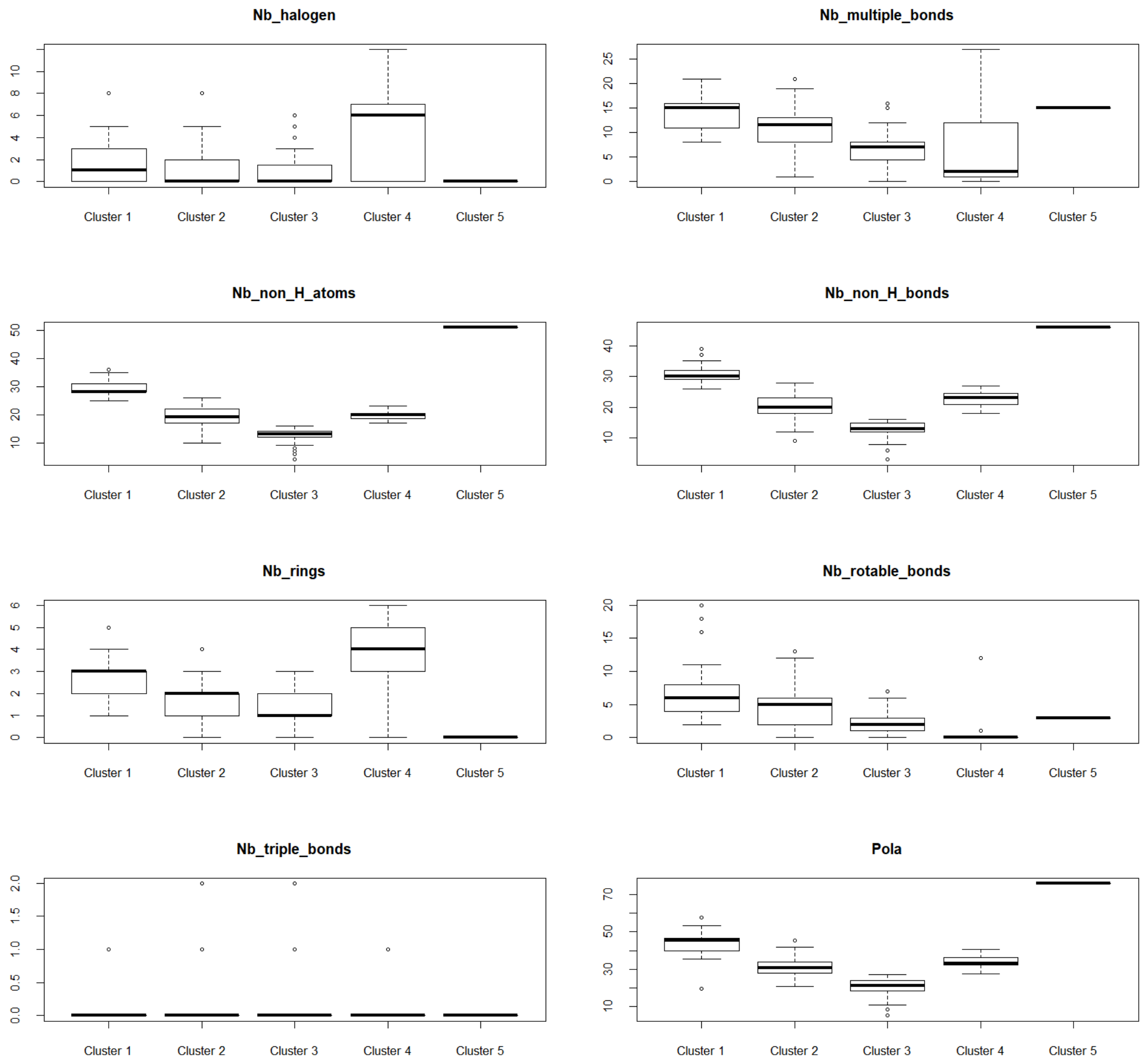


**Figure S4** (continued)

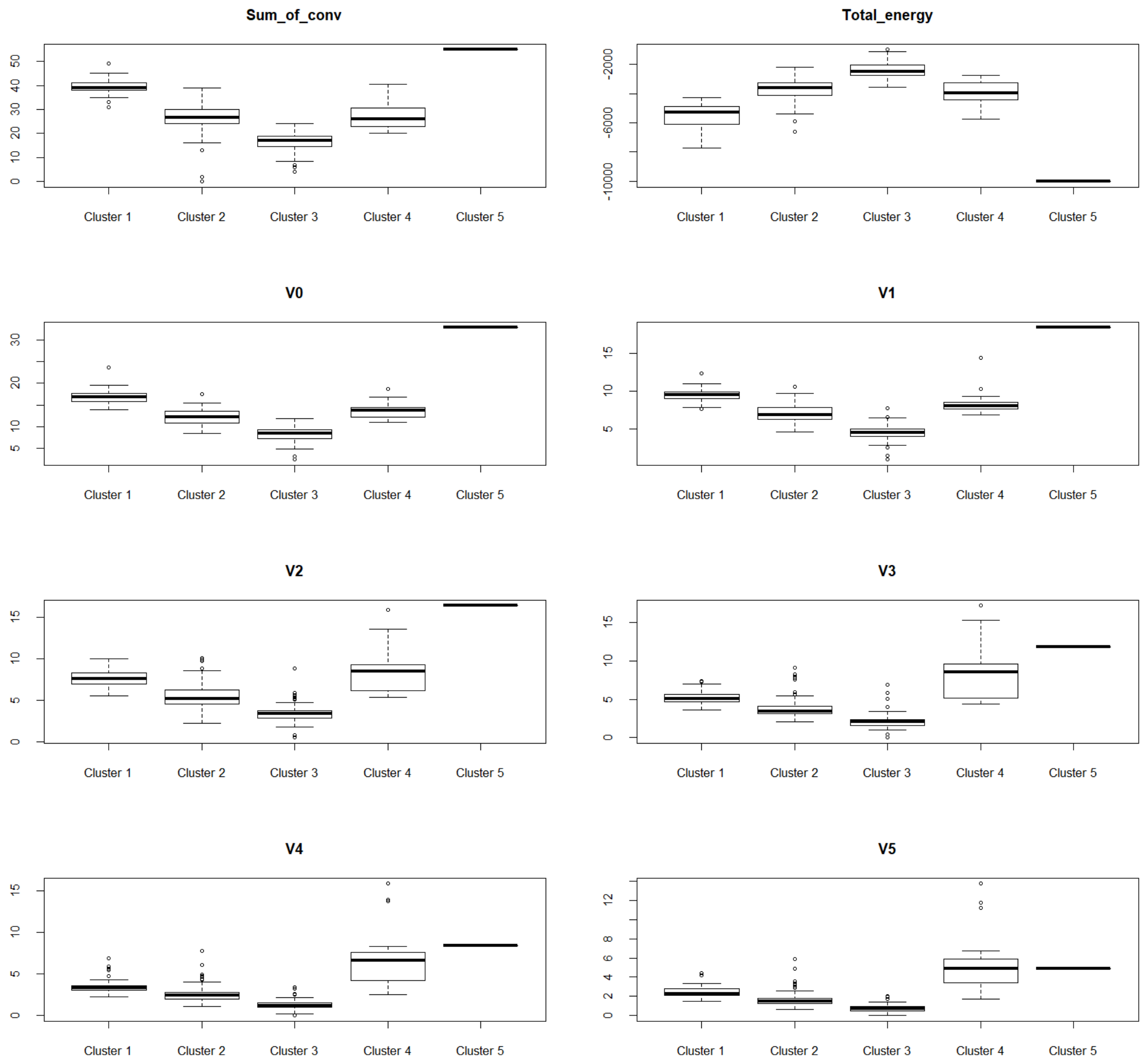


**Figure S4** (continued)

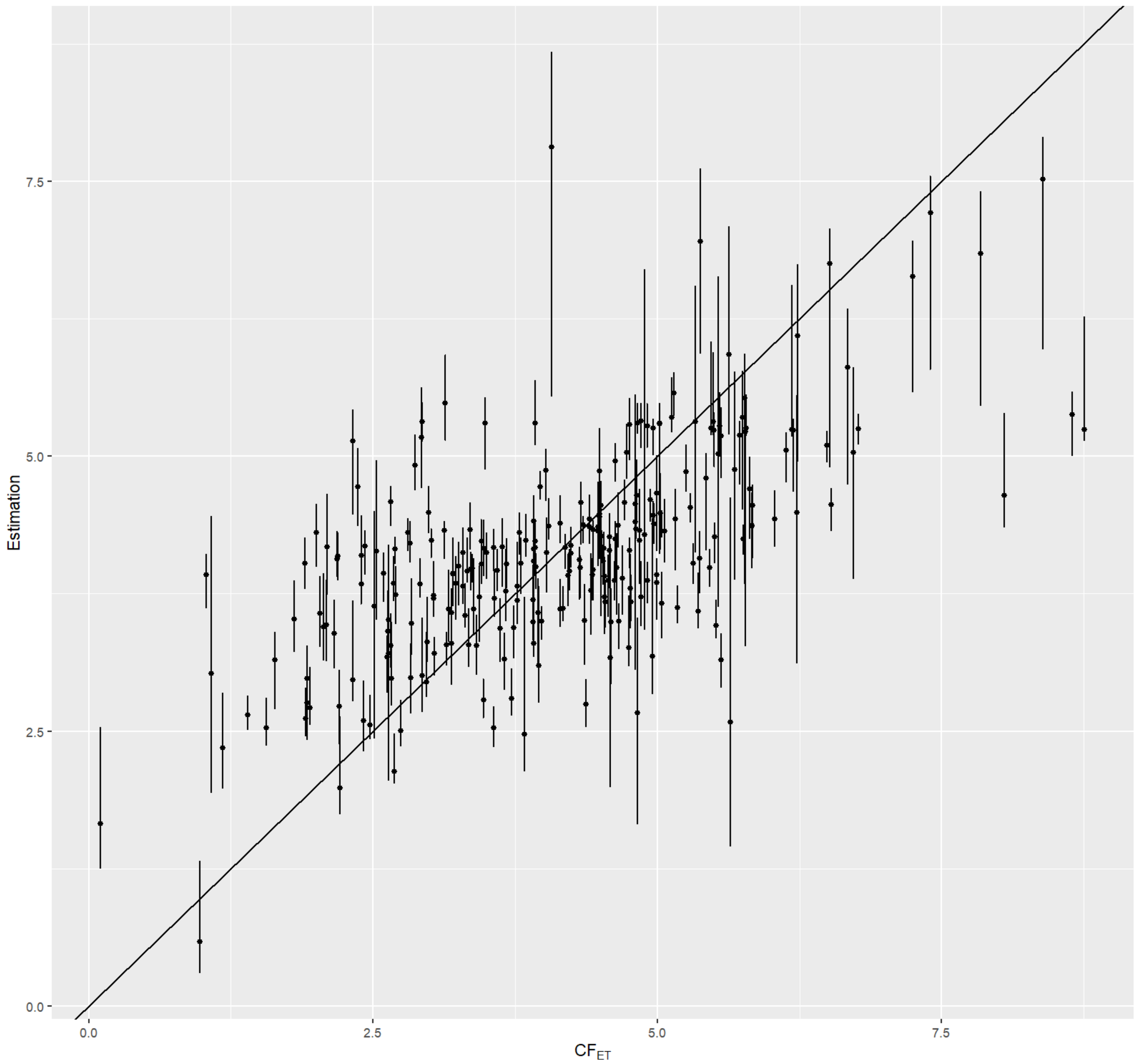


**Figure S5**- Estimation of $CF_{ET}$ according to the value in Usetox®. The estimation is the median of the estimations made using the best method of the cluster during the comparison procedure. The bar represents the 5% and the 95% quantiles of these individual estimations.

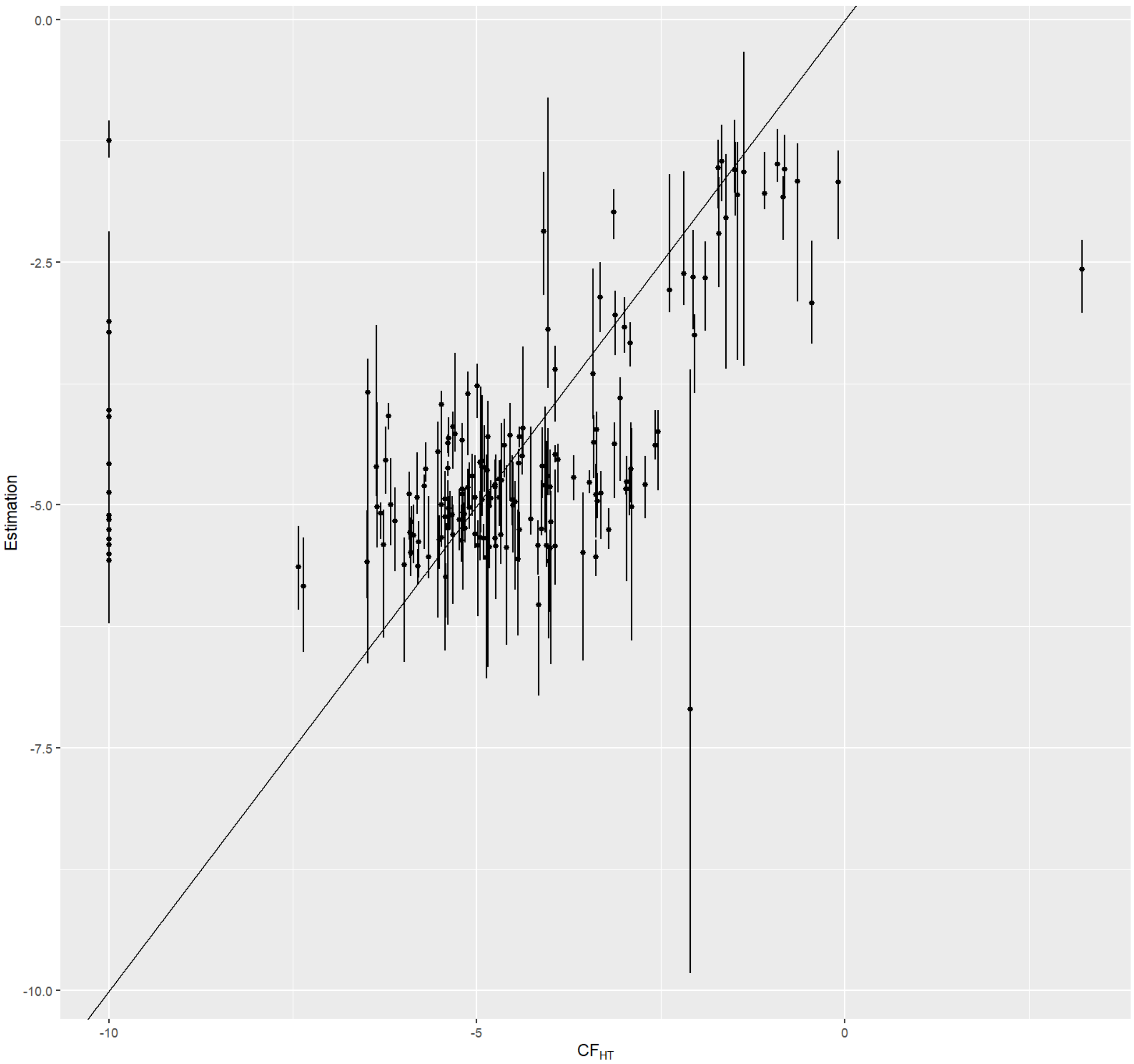


**Figure S6**- Estimation of $CF_{HT}$ according to the value in Usetox®. The estimation is the median of the estimations made using the best method of the cluster during the comparison procedure. The bar represents the 5% and the 95% quantiles of these individual estimations.

### 2. Supplemental Tables

**Table S1-** CAS number and name of the 274 common compounds between TyPol and USEtox databases and their associated $CF_{ET}$ and $CF_{HT}$ values. NA means that the there is no value in USEtox for this compound.

| CAS | Name | $CF_{HT}$ | $CF_{ET}$ | Cluster |
|---|---|---|---|---|
| 101-20-2 | Triclocarban | NA | 6.79E+05 | 2 |
| 101-21-3 | Chlorpropham | 9.60E-06 | 2.74E+03 | 3 |
| 101-42-8 | Fenuron | NA | 1.39E+03 | 3 |
| 101200-48-0 | Tribenuron-methyl | 1.80E-05 | 3.39E+02 | 1 |

| | | | | |
|---|---|---|---|---|
| 101205-02-1 | Cycloxydim | NA | 1.56E+02 | 2 |
| 1024-57-3 | Heptachlor epoxide | 0.81 | 3.17E+05 | 4 |
| 102851-06-9 | tau-Fluvalinate | NA | 4.28E+05 | 1 |
| 103-90-2 | Acetamide, n-(4-hydroxyphenyl) | 1.00E-06 | 4.33E+01 | 3 |
| 1031-07-08 | Endosulfan sulfate | NA | 1.05E+05 | 4 |
| 103361-09-7 | Flumioxazin | NA | 2.38E+05 | 1 |
| 104-40-5 | P-nonylphenol | NA | 3.24E+04 | 2 |
| 10540-29-1 | Tamoxifen | NA | 4.40E+05 | 1 |
| 105512-06-9 | Clodinafop-propargyl | NA | 1.39E+04 | 2 |
| 106-44-5 | P-cresol | NA | 5.51E+02 | 3 |
| 1071-83-6 | Glyphosate | 4.30E-07 | 1.60E+02 | 3 |
| 107534-96-3 | Tebuconazole | 2.00E-05 | 3.43E+04 | 2 |
| 108-62-3 | Metaldehyde (tetramer) | NA | 1.23E+02 | 3 |
| 110488-70-5 | Dimethomorph | NA | 1.37E+03 | 1 |
| 111479-05-1 | Propaquizafop | NA | 6.71E+04 | 1 |
| 111991-09-4 | Nicosulfuron | NA | 3.25E+02 | 1 |
| 114-07-08 | Erythromycin | NA | 1.07E+04 | 5 |
| 114369-43-6 | Fenbuconazole | 2.80E-05 | 5.87E+04 | 2 |
| 115-29-7 | Endosulfan | 8.10E-05 | 2.97E+05 | 4 |
| 116-06-03 | Aldicarb | 0.00028 | 2.35E+04 | 3 |
| 117-81-7 | Di-(2-ethylhexyl)-phthalate (DEHP) | 4.10E-06 | 1.61E+02 | 1 |
| 117-84-0 | Di(n-octyl) phthalate | NA | 1.51E+01 | 1 |
| 118-74-1 | Hexachlorobenzene | 0.0091 | 5.13E+04 | 3 |
| 119446-68-3 | Difenoconazole | NA | 6.43E+04 | 1 |
| 0120-12-7 | Anthracene | 0.0029 | 1.51E+05 | 3 |
| 120-72-9 | Indole | 0 | 2.95E+03 | 3 |
| 120068-37-3 | Fipronil | 0.00089 | 1.08E+06 | 2 |
| 121-75-5 | Malathion | 5.80E-07 | 3.11E+04 | 2 |
| 1214-39-7 | 1h-purin-6-amine, n-(phenylmethyl)- | NA | 5.01E+02 | 2 |
| 121552-61-2 | Cga 219417 (cyprodinil) | NA | 1.40E+04 | 2 |
| 122-14-5 | Fenitrothion | 8.80E-05 | 9.87E+04 | 2 |

| | | | | |
|---|---|---|---|---|
| 122-34-9 | Simazine | 7.50E-05 | 3.89E+04 | 3 |
| 12427-38-2 | Maneb | 1.20E-05 | 3.44E+04 | 3 |
| 128639-02-1 | Carfentrazone-ethyl | NA | 1.17E+05 | 2 |
| 129-00-0 | Pyrene | 0.00047 | 6.47E+05 | 2 |
| 131-11-3 | Dimethylphthalate (DMP) | NA | 8.35E+01 | 3 |
| 131341-86-1 | Fludioxonil | NA | 4.94E+04 | 2 |
| 131860-33-8 | Azoxystrobin | NA | 3.85E+04 | 1 |
| 13194-48-4 | O-ethyl s,s-dipropyl phosphorodithioate | 0.00049 | 1.06E+05 | 2 |
| 133-06-02 | Captan | 7.60E-06 | 4.24E+04 | 2 |
| 133-07-03 | Folpet | 4.80E-06 | 5.58E+05 | 2 |
| 135158-54-2 | Cga 245704 | NA | 9.02E+03 | 3 |
| 13684-56-5 | Desmedipham | NA | 4.23E+04 | 2 |
| 13684-63-4 | Phenmedipham | 1.30E-06 | 2.10E+04 | 2 |
| 137-26-8 | Thiram | 1.20E-05 | 2.90E+05 | 3 |
| 138261-41-3 | Imidacloprid | 6.80E-06 | 1.60E+03 | 2 |
| 140-66-9 | P-(1,1,3,3-tetramethylbutyl)phenol | NA | 1.74E+04 | 2 |
| 142459-58-3 | Fluthiamide | NA | 8.71E+04 | 2 |
| 143-50-0 | Kepone | 0.042 | 5.95E+05 | 4 |
| 143390-89-0 | Bas 490f | 1.20E-06 | 8.18E+04 | 2 |
| 14698-29-4 | Oxolinic acid | 6.50E-06 | 1.09E+05 | 2 |
| 148-79-8 | Thiabendazole | 3.70E-06 | 1.70E+04 | 3 |
| 15299-99-7 | N,n-diethyl-2-(1-naphthalenyloxy)propanamide | 1.90E-06 | 1.96E+03 | 2 |
| 15307-86-5 | Diclofenac | 0.00043 | 9.72E+02 | 2 |
| 15545-48-9 | Chlortoluron | NA | 1.34E+03 | 3 |
| 1563-38-8 | Carbofuran phenol | NA | 2.57E+03 | 3 |
| 1563-66-2 | Carbofuran | 1.00E-04 | 5.61E+04 | 2 |
| 15687-27-1 | Ibuprofen | 0 | 1.17E+02 | 3 |
| 1570-64-5 | 2-methyl-4-chlorophenol | NA | 3.64E+03 | 3 |
| 1582-09-08 | Trifluralin | 9.30E-05 | 5.38E+04 | 2 |
| 15972-60-8 | Alachlor | NA | 3.81E+04 | 2 |

| | | | | |
|---|---|---|---|---|
| 16118-49-3 | Carbetamide | NA | 1.08E+03 | 2 |
| 16672-87-0 | Ethephon | 1.40E-05 | 6.80E+02 | 3 |
| 1689-84-5 | Bromoxynil | 8.80E-06 | 8.23E+03 | 3 |
| 1689-99-2 | Bromoxynil octanoate | 5.10E-06 | 9.27E+04 | 2 |
| 1698-60-8 | Chloridazon | NA | 4.65E+03 | 3 |
| 1702-17-6 | 3,6-dichloropicolinic acid | NA | 4.55E+02 | 3 |
| 173584-44-6 | Dpx-mp062 | NA | 7.78E+04 | 1 |
| 1746-01-06 | 2,3,7,8-TetraCDD | 1.70E+03 | 4.72E+06 | 2 |
| 1746-81-2 | Monolinuron | NA | 9.65E+03 | 3 |
| 1897-45-6 | Chlorothalonil | 1.00E-05 | 5.72E+05 | 3 |
| 19044-88-3 | Oryzalin | 3.10E-06 | 1.10E+05 | 2 |
| 191-24-2 | Benzo[g,h,i]perylene | 0.00073 | NA | 4 |
| 1912-24-9 | Atrazine | 5.40E-05 | 4.37E+04 | 3 |
| 1918-00-9 | Dicamba | 6.30E-06 | 9.43E+02 | 3 |
| 1918-02-1 | Picloram | 2.00E-06 | 1.59E+03 | 3 |
| 1918-16-7 | Propachlor | 4.40E-06 | 3.72E+04 | 3 |
| 1929-77-7 | Vernolate | 1.50E-05 | 2.20E+03 | 3 |
| 193-39-5 | Indeno[1,2,3-cd]-pyrene | 0.019 | NA | 4 |
| 19666-30-9 | Oxadiazon | 0.00075 | 3.20E+05 | 2 |
| 205-99-2 | Benzo[b]fluoranthene | 0.081 | NA | 4 |
| 2050-68-2 | PCB-15 | NA | 2.74E+04 | 3 |
| 2051-60-7 | PCB-1 | NA | 2.05E+03 | 3 |
| 2051-61-8 | PCB-2 | NA | 1.55E+03 | 3 |
| 206-44-0 | Fluoranthene | 0.001 | 5.70E+04 | 2 |
| 207-08-09 | Benzo[k]fluoranthene | 0.035 | NA | 4 |
| 21087-64-9 | Metribuzin | 4.20E-06 | 4.73E+03 | 3 |
| 21725-46-2 | Cyanazine | 0.00043 | 4.28E+04 | 3 |
| 218-01-09 | Chrysene | 0.013 | NA | 2 |
| 22071-15-4 | Ketoprofen | 0 | NA | 2 |
| 2303-16-4 | Diallate | 0.00021 | 2.25E+03 | 3 |
| 2303-17-5 | Triallate | 4.30E-05 | 9.34E+03 | 2 |

| | | | | |
|---|---|---|---|---|
| 23103-98-2 | Pirimicarb | 6.90E-06 | 8.24E+02 | 2 |
| 2312-35-8 | Propargite | 1.00E-04 | 7.21E+04 | 2 |
| 23135-22-0 | Oxamyl | 1.10E-05 | 8.09E+03 | 3 |
| 23564-05-08 | Thiophanate-methyl | 4.70E-06 | 3.64E+03 | 2 |
| 2385-85-5 | Mirex | 0.024 | 8.59E+02 | 4 |
| 23950-58-5 | Pronamide | 3.70E-05 | 2.15E+03 | 2 |
| 197143 | Dodine | 4.40E-07 | 8.51E+03 | 2 |
| 24579-73-5 | Propamocarb | 1.40E-06 | 8.27E+01 | 3 |
| 25057-89-0 | Bentazone | 3.30E-06 | 1.00E+02 | 2 |
| 25812-30-0 | Gemfibrozil | 3.60E-05 | NA | 2 |
| 26225-79-6 | Ethofumesate | NA | 1.96E+03 | 2 |
| 26761-40-0 | Diisodecyl phthalate | NA | 1.30E+00 | 1 |
| 26787-78-0 | Amoxicillin | NA | 5.28E+06 | 1 |
| 27304-13-8 | Oxychlordane | NA | 7.16E+04 | 4 |
| 27314-13-2 | Norflurazon | 4.10E-06 | 2.54E+04 | 2 |
| 28553-12-0 | Diisononyl phthalate | NA | 9.50E+00 | 1 |
| 2921-88-2 | Chloropyrifos | 0.0012 | 3.12E+06 | 2 |
| 297-78-9 | Isobenzan | 0 | 8.14E+04 | 4 |
| 298-46-4 | Carbamazepine | 6.30E-06 | 3.90E+02 | 2 |
| 3060-89-7 | Metobromuron | NA | 6.72E+02 | 3 |
| 309-00-2 | Aldrin | 0.033 | 1.34E+05 | 4 |
| 32809-16-8 | Procymidone | 3.30E-06 | 4.51E+02 | 2 |
| 330-54-1 | Diuron | 1.80E-05 | 3.00E+04 | 3 |
| 330-55-2 | Linuron | 9.90E-05 | 9.93E+04 | 3 |
| 33284-50-3 | PCB-7 | NA | 2.21E+04 | 3 |
| 333-41-5 | Diazinon | 0.00042 | 9.26E+04 | 2 |
| 3337-71-1 | Asulam | 2.20E-06 | 1.08E+02 | 3 |
| 3347-22-6 | Dithianone | 1.40E-05 | 2.12E+04 | 2 |
| 33629-47-9 | Butralin | NA | 9.85E+04 | 2 |
| 3380-34-5 | 5-chloro-2-(2,4-dichlorophenoxy)phenol | NA | 6.60E+04 | 2 |
| 34014-18-1 | Tebuthiuron | 4.10E-06 | 6.35E+03 | 3 |

| | | | | |
|---|---|---|---|---|
| 34123-59-6 | Isoproturon | NA | 5.78E+04 | 3 |
| 34256-82-1 | Acetochlor | NA | 3.38E+04 | 2 |
| 34883-43-7 | 2,4'-dichlorobiphenyl | NA | 2.52E+04 | 3 |
| 35554-44-0 | Imazalil base | 2.50E-05 | 8.14E+03 | 2 |
| 36734-19-7 | Rovral (Iprodione) | 2.30E-05 | 3.11E+04 | 2 |
| 3739-38-6 | M-phenoxybenzoic acid | NA | 2.31E+02 | 2 |
| 39148-24-8 | Fosetyl-aluminium | 3.30E-07 | 7.45E+02 | 2 |
| 40321-76-4 | 1,2,3,7,8-pentachlorodibenzo-p-dioxin | NA | 5.71E+08 | 4 |
| 40487-42-1 | Pendimethalin | 1.60E-06 | 2.29E+05 | 2 |
| 41394-05-02 | Metamitron | NA | 2.49E+02 | 3 |
| 41483-43-6 | Bupirimate | NA | 8.41E+03 | 2 |
| 41859-67-0 | Bezafibrate | 3.00E-05 | 6.43E+02 | 2 |
| 42835-25-6 | Flumequine | NA | 4.33E+03 | 2 |
| 42874-03-03 | Oxyfluorfen | 0.002 | 3.19E+04 | 2 |
| 439-14-5 | Diazepam | 0 | NA | 2 |
| 443-48-1 | Metronidazole | 3.80E-06 | 8.07E+01 | 3 |
| 465-73-6 | Isodrin | NA | 6.08E+05 | 4 |
| 481-39-0 | 5-hydroxy-1,4-naphthoquinone | NA | 4.60E+04 | 3 |
| 50-28-2 | Estradiol | 0 | 1.12E+08 | 4 |
| 50-29-3 | p,p'-DDT | 0.0065 | 1.39E+05 | 2 |
| 50-32-8 | Benzo[a]pyrene | 0.032 | 8.44E+03 | 4 |
| 50-78-2 | Acetylsalicylic acid | 0 | NA | 3 |
| 51-03-6 | Piperonyl butoxide | 1.80E-05 | 2.06E+04 | 2 |
| 51207-31-9 | 2,3,7,8-TetraCDF | NA | 4.45E+08 | 2 |
| 51218-45-2 | Metolachlor | 3.30E-06 | 3.35E+04 | 2 |
| 51338-27-3 | Diclofop-methyl | NA | 6.48E+04 | 2 |
| 51481-61-9 | Cimetidine | 0 | NA | 2 |
| 518-47-8 | Fluorescein sodium | NA | 1.09E+01 | 2 |
| 52315-07-08 | Cypermethrin | 1.10E-05 | 2.51E+07 | 1 |
| 52645-53-1 | Permethrin | 4.10E-06 | 5.88E+05 | 1 |
| 52888-80-9 | Prosulfocarb | NA | 1.55E+04 | 2 |

| | | | | |
|---|---|---|---|---|
| 52918-63-5 | Deltamethrin | 2.00E-05 | 1.72E+06 | 1 |
| 53-16-7 | Estrone | NA | 1.18E+04 | 4 |
| 53-70-3 | Dibenz(a,h)anthracene | 0.14 | 3.05E+03 | 4 |
| 53112-28-0 | Pyrimethanil | NA | 1.70E+03 | 3 |
| 54-31-9 | Furosemide | 3.70E-06 | NA | 2 |
| 55179-31-2 | Bitertanol | 9.30E-05 | 8.11E+03 | 2 |
| 55219-65-3 | Triadimenol | 1.50E-05 | 2.85E+03 | 2 |
| 55335-06-03 | Triclopyr | NA | 2.43E+03 | 3 |
| 555-37-3 | Neburon | NA | 2.68E+04 | 2 |
| 5598-13-0 | Chlorpyrifos methyl | 0.0012 | 3.64E+05 | 2 |
| 56-38-2 | Parathion | 0.00011 | 3.40E+06 | 2 |
| 56-55-3 | Benz[a]anthracene | 0.0086 | 6.77E+05 | 2 |
| 563-12-2 | Ethion | 0.0013 | 1.05E+05 | 4 |
| 57-41-0 | Phenytoin | 3.30E-05 | NA | 2 |
| 57-62-5 | Aureomycin | NA | 4.33E+02 | 1 |
| 57-63-6 | Ethinyl estradiol | 0.0079 | 1.57E+06 | 4 |
| 57-68-1 | Sulfamethazine | 1.20E-06 | NA | 2 |
| 57-74-9 | Chlordane | 0.12 | 9.17E+04 | 4 |
| 57653-85-7 | 1,2,3,6,7,8-hexachlorodibenzo-p-dioxin | NA | 1.52E+06 | 4 |
| 57837-19-1 | Metalaxyl | 1.60E-06 | 4.78E+02 | 2 |
| 57966-95-7 | Cymoxanil | NA | 5.45E+03 | 3 |
| 58-08-2 | Caffeine | 0 | 3.49E+04 | 3 |
| 58-14-0 | Pyrimethamine | 0 | 2.98E+03 | 2 |
| 58-89-9 | Gamma-HCH (lindane) | 0.0012 | 1.44E+05 | 3 |
| 5915-41-3 | Terbuthylazine | NA | 2.36E+05 | 3 |
| 5989-27-5 | D-limonene | 4.80E-06 | 1.45E+02 | 3 |
| 60-51-5 | Dimethoate | 1.10E-05 | 8.95E+03 | 3 |
| 60-54-8 | Tetracycline | NA | 1.25E+02 | 1 |
| 60-57-1 | Dieldrin | 0.15 | 3.10E+05 | 4 |
| 60168-88-9 | Fenarimol | 0.00012 | 1.73E+04 | 2 |
| 60207-90-1 | Propiconazole | 4.10E-05 | 1.11E+04 | 2 |

| | | | | |
|---|---|---|---|---|
| 608-73-1 | 1,2,3,4,5,6-hexachlorocyclohexane | 0.00077 | 6.99E+04 | 3 |
| 61-82-5 | Amitrole | 7.00E-05 | 4.90E+02 | 3 |
| 61213-25-0 | Flurochloridone | NA | 1.05E+04 | 2 |
| 62-73-7 | Dichlorvos | 0.00041 | 3.62E+05 | 3 |
| 62924-70-3 | Flumetralin | NA | 4.81E+05 | 1 |
| 63-25-2 | Carbaryl | 9.50E-05 | 2.29E+04 | 3 |
| 64-19-7 | Acetic acid | NA | 2.50E+01 | 3 |
| 64902-72-3 | Chlorsulfuron | 7.80E-06 | 6.12E+03 | 2 |
| 66215-27-8 | Cyromazine | 2.10E-05 | 1.56E+03 | 3 |
| 66246-88-6 | Penconazole | 0.00013 | 8.39E+03 | 2 |
| 67129-08-02 | Metazachlor | NA | 3.72E+03 | 2 |
| 67375-30-8 | alpha-Cypermethrin | 1.40E-05 | 1.75E+07 | 1 |
| 67564-91-4 | Fenpropimorph | NA | 5.89E+03 | 2 |
| 67747-09-05 | Prochloraz | 0.0027 | 1.96E+05 | 2 |
| 68-35-9 | Sulfadiazine | NA | 5.87E+03 | 2 |
| 68359-37-5 | Cyfluthrin | 3.80E-05 | 2.44E+08 | 1 |
| 69-53-4 | Ampicillin | NA | 1.53E+02 | 2 |
| 69377-81-7 | Fluroxypyr | NA | 1.46E+03 | 3 |
| 70630-17-0 | Metalaxyl-M | NA | 1.08E+03 | 2 |
| 7085-19-0 | Mecoprop | 3.80E-05 | 4.31E+02 | 3 |
| 709-98-8 | Propanil | 1.20E-05 | 2.07E+05 | 3 |
| 72-20-8 | Endrin | 0.019 | 5.90E+06 | 4 |
| 72-33-3 | Mestranol | 0 | NA | 4 |
| 72-54-8 | DDD | 0.35 | 1.36E+06 | 2 |
| 72-55-9 | p,p'-DDE | 0.0042 | 3.51E+05 | 2 |
| 723-46-6 | Sulfamethoxazole | 1.30E-06 | 2.35E+03 | 2 |
| 731-27-1 | Tolyfluanide | NA | 1.80E+05 | 2 |
| 732-11-6 | Phosmet | 2.20E-05 | 6.91E+05 | 2 |
| 73334-07-03 | Iopromide | 6.40E-07 | 1.20E+01 | 1 |
| 73590-58-6 | Omeprazole | 1.30E-05 | NA | 2 |
| 738-70-5 | Trimethoprim | 7.50E-06 | 4.98E+02 | 2 |

| | | | | |
|---|---|---|---|---|
| 74070-46-5 | Aclonifen | NA | 3.31E+05 | 2 |
| 74223-64-6 | Metsulfuron-methyl | 1.60E-06 | 1.07E+04 | 2 |
| 759-94-4 | Eptc | 6.20E-06 | 8.54E+02 | 3 |
| 76-44-8 | Heptachlor | 0.021 | 6.73E+04 | 4 |
| 77732-09-03 | Oxadixyl | NA | 7.93E+01 | 2 |
| 79-57-2 | Oxytetracylcine | NA | 6.81E+03 | 1 |
| 79-94-7 | 2,2-bis(4-hydroxy-3,5-dibromophenyl)propane | NA | 3.09E+04 | 2 |
| 79127-80-3 | Fenoxycarb | NA | 1.65E+04 | 2 |
| 79277-27-3 | Harmony | 3.10E-05 | 6.43E+04 | 2 |
| 79622-59-6 | Fluazinam | NA | 3.45E+05 | 1 |
| 80-05-7 | 4,4'-Isopropylidenediphenol | 3.00E-06 | 4.18E+03 | 2 |
| 8001-35-2 | Toxaphene | 0.23 | 5.27E+05 | 4 |
| 8018-01-7 | Mancozeb | 5.80E-06 | 2.63E+04 | 3 |
| 80844-07-01 | Etofenprox | 0.0011 | 2.11E+02 | 1 |
| 81-81-2 | Warfarin | 0.0011 | 2.70E+02 | 2 |
| 81777-89-1 | Clomazone | NA | 3.89E+03 | 2 |
| 82558-50-7 | Isoxaben | 1.80E-05 | 2.72E+04 | 2 |
| 82657-4-3 | Bifenthrin | 0.00034 | 3.29E+06 | 1 |
| 83-79-4 | Rotenone | 0.00012 | 2.16E+05 | 1 |
| 83164-33-4 | Diflufenican | NA | 8.48E+02 | 1 |
| 0834-12-8 | Ametryne | NA | 3.80E+04 | 3 |
| 84-66-2 | Diethylphthalate (DEP) | 3.70E-08 | 2.11E+02 | 3 |
| 84-74-2 | Dibutylphthalate (DBP) | 3.20E-07 | 3.16E+03 | 2 |
| 85-01-8 | Phenanthrene | 0.00039 | 8.21E+03 | 3 |
| 85-41-6 | Phthalimide | NA | 4.21E+02 | 3 |
| 85-68-7 | Butyl benzyl phthalate | 7.70E-07 | 2.83E+03 | 2 |
| 86-50-0 | Methyl azinphos | 8.40E-05 | 2.69E+05 | 2 |
| 86-73-7 | Fluorene | 7.70E-05 | 1.80E+03 | 3 |
| 86-87-3 | Naphthaleneacetic acid | 0 | 6.43E+01 | 3 |
| 87-51-4 | Indole-3-acetic acid | 0 | 4.60E+02 | 3 |
| 87-86-5 | Pentachlorophenol | 0.00038 | 4.53E+04 | 3 |

| | | | | |
|---|---|---|---|---|
| 87392-12-9 | S-Metolachlor | NA | 5.72E+04 | 2 |
| 87674-68-8 | Dimethenamid | NA | 7.02E+04 | 2 |
| 88-99-3 | O-phthalic acid | NA | 2.60E+02 | 3 |
| 886-50-0 | Terbutryn | 0.00063 | 3.22E+04 | 3 |
| 88671-89-0 | Myclobutanil | 6.30E-06 | 1.49E+04 | 2 |
| 90-43-7 | 2-Phenylphenol | 4.10E-06 | 4.55E+03 | 3 |
| 9006-42-2 | Metiram | 4.90E-07 | 1.03E+03 | 3 |
| 90717-03-06 | Quinmerac | NA | 2.49E+02 | 2 |
| 91465-08-06 | Lambda-cyhalothrin | NA | 6.93E+07 | 1 |
| 92-52-4 | Biphenyl | 6.80E-07 | 1.10E+03 | 3 |
| 93106-60-6 | Enrofloxacin | NA | 1.69E+06 | 1 |
| 94-74-6 | 2-Methyl-4-chlorophenoxyacetic acid | 6.80E-05 | 9.40E+02 | 3 |
| 94-75-7 | 2-(2,4-dichlorophenoxy)acetic acid | 1.60E-05 | 4.30E+02 | 3 |
| 94-82-6 | 2,4-DB | 9.50E-06 | 6.92E+02 | 3 |
| 94125-34-5 | Prosulfuron | NA | 9.07E+04 | 1 |
| 94361-06-05 | Cyproconazole | NA | 2.30E+03 | 2 |
| 95-48-7 | o-cresol | 5.40E-07 | 2.96E+02 | 3 |
| 95-76-1 | 3,4-Dichloroaniline | NA | 5.24E+03 | 3 |
| 97-23-4 | Phenol,2,2'-methylenebis 4-chloro | NA | 3.02E+04 | 2 |
| 98-86-2 | Acetophenone | 4.50E-08 | 3.63E+01 | 3 |
| 99-30-9 | 2,6-dichloro-4-nitroaniline | 1.00E-05 | 8.25E+03 | 3 |
| 99607-70-2 | Cloquintocet-mexyl | NA | 7.00E+03 | 2 |
| 999-81-5 | Chlormequat chloride | 0 | 8.83E+01 | 3 |

**Table S2** - Summary of the descriptors included in the whole TyPol database (in the first three columns) and for the 274 compounds common between TyPol and UseTox databases (in the last three columns)

| Descriptors | TyPol | | | TyPol & UseTox | | |
|---|---|---|---|---|---|---|
| | Min global | Max global | Nb NA (%) | Min commun | Max commun | Nb NA (%) |
| Connectivity index chi-0 | 3.58 | 44.67 | 1.09 | 3.58 | 38.96 | 1.46 |

| | | | | | | |
|---|---|---|---|---|---|---|
| Connectivity index chi-1 | 1.73 | 29.5 | 0.18 | 1.73 | 23.43 | 0 |
| Connectivity index chi-2 | 1.73 | 27.87 | 1.09 | 1.73 | 23.46 | 1.46 |
| Connectivity index chi-3 | 0 | 24.49 | 0.18 | 0 | 19.66 | 0 |
| Connectivity index chi-4 | 0 | 20.34 | 1.09 | 0 | 14.47 | 1.46 |
| Connectivity index chi-5 | 0 | 16.52 | 1.09 | 0 | 11.81 | 1.46 |
| Electric dipole moment | -8.8 | 24.14 | 0.18 | -8.8 | 15.19 | 0 |
| HOMO energy | -15.04 | -0.26 | 0.18 | -15.04 | -1.81 | 0 |
| LUMO energy | -9.96 | 8.47 | 0.18 | -4.38 | 4.63 | 0 |
| Molecular mass | 16 | 873.2 | 0 | 16 | 791.12 | 0 |
| Molecular surface area (Connolly) | 0 | 698.85 | 0 | 0 | 560.27 | 0 |
| Number of Carbon atoms | 2 | 48 | 0 | 2 | 37 | 0 |
| Number of Chlorine atoms | 0 | 12 | 0 | 0 | 12 | 0 |
| Number of Fluorine atoms | 0 | 6 | 0 | 0 | 6 | 0 |
| Number of Hydrogen atoms | 0 | 116 | 0 | 0 | 67 | 0 |
| Number of Nitrogen atoms | 0 | 44 | 0 | 0 | 44 | 0 |
| Number of Oxygen atoms | 0 | 15 | 0 | 0 | 13 | 0 |
| Number of Phosphorus atoms | 0 | 3 | 0 | 0 | 3 | 0 |
| Number of Sulfur atoms | 0 | 4 | 0 | 0 | 4 | 0 |
| Number of aromatic bonds | 0 | 27 | 0.18 | 0 | 27 | 0 |

| | | | | | | |
|---|---|---|---|---|---|---|
| Number of atoms | 8 | 134 | 0 | 8 | 118 | 0 |
| Number of bonds | 4 | 140 | 0.18 | 7 | 113 | 0 |
| Number of circuits | 0 | 47 | 0.18 | 0 | 47 | 0 |
| Number of double bonds | 0 | 10 | 0.18 | 0 | 6 | 0 |
| Number of halogen atoms | 0 | 12 | 0 | 0 | 12 | 0 |
| Number of multiple bonds | 0 | 27 | 0.18 | 0 | 27 | 0 |
| Number of non-H atoms | 4 | 62 | 0 | 4 | 51 | 0 |
| Number of non-H bonds | 2 | 68 | 0.18 | 3 | 46 | 0 |
| Number of rings | 0 | 7 | 0.18 | 0 | 6 | 0 |
| Number of rotatable bonds | 0 | 28 | 0.18 | 0 | 20 | 0 |
| Number of triple bonds | 0 | 3 | 0.18 | 0 | 2 | 0 |
| Polarizability | 5.13 | 94.85 | 0.18 | 5.13 | 75.99 | 0 |
| Sum of conventional bond order | 0 | 74 | 0.18 | 0 | 55 | 0 |
| Total energy | -11625.4 | 5030.03 | 0.18 | -10037.4 | -952.91 | 0 |
| Valence connectivity index chi-0 | 2.36 | 38.22 | 1.09 | 2.36 | 32.94 | 1.46 |
| Valence connectivity index chi-1 | 0.93 | 22.86 | 1.09 | 0.93 | 18.49 | 1.46 |
| Valence connectivity index chi-2 | 0.52 | 18.96 | 1.09 | 0.52 | 16.47 | 1.46 |
| Valence connectivity index chi-3 | 0 | 17.47 | 1.09 | 0 | 17.29 | 1.46 |

| | | | | | | |
|---|---|---|---|---|---|---|
| Valence connectivity index chi-4 | 0 | 15.94 | 1.09 | 0 | 15.9 | 1.46 |

**Table S3**- Quantiles for the different methods and clusters for the predictions of the $CF_{ET}$. These values are used to obtain the Figure 4. The best models are in bold for each cluster.

| quantiles | RF | PLS | SVM | Cluster-RF | Cluster-SVM | Cluster-PLS |
|---|---|---|---|---|---|---|
| | | | **Cluster 1** | | | |
| 0% | 0,00 | 0,00 | 0,00 | 0,00 | **0,00** | 0,00 |
| 25% | 0,63 | 0,86 | 0,66 | 0,66 | **0,48** | 0,71 |
| 50% | 1,25 | 1,67 | 1,23 | 1,25 | **1,18** | 1,57 |
| 75% | 2,02 | 2,65 | 2,12 | 2,03 | **1,84** | 2,26 |
| 100% | 3,87 | 6,51 | 4,10 | 3,53 | **9,45** | 8,69 |
| | | | **Cluster 2** | | | |
| 0% | 0,00 | 0,00 | 0,00 | **0,00** | 0,00 | 0,00 |
| 25% | 0,34 | 0,37 | 0,33 | **0,32** | 0,38 | 0,38 |
| 50% | 0,63 | 0,71 | 0,66 | **0,60** | 0,67 | 0,68 |
| 75% | 1,09 | 1,27 | 1,17 | **1,09** | 1,16 | 1,17 |
| 100% | 3,73 | 4,40 | 4,69 | **3,68** | 5,75 | 5,44 |
| | | | **Cluster 3** | | | |
| 0% | 0,00 | 0,00 | 0,00 | **0,00** | 0,00 | 0,00 |
| 25% | 0,31 | 0,36 | 0,34 | **0,27** | 0,37 | 0,35 |
| 50% | 0,66 | 0,79 | 0,64 | **0,62** | 0,67 | 0,63 |
| 75% | 1,06 | 1,18 | 1,08 | **1,05** | 1,06 | 1,02 |
| 100% | 2,56 | 2,15 | 2,67 | **2,79** | 3,39 | 3,90 |
| | | | **Cluster 4** | | | |
| 0% | 0,01 | 0,00 | 0,01 | 0,00 | **0,00** | 0,00 |
| 25% | 0,40 | 0,31 | 0,33 | 0,38 | **0,34** | 0,46 |
| 50% | 0,65 | 0,70 | 0,59 | 0,61 | **0,57** | 0,92 |
| 75% | 1,45 | 1,49 | 1,55 | 1,46 | **1,53** | 1,96 |
| 100% | 3,98 | 6,11 | 5,08 | 3,56 | **3,98** | 12,44 |

**Table S4**- The five most important molecular descriptors for each best model for each cluster for the $CF_{ET}$. Descriptors are listed from top to bottom in decreasing order of importance.

| Cluster 1: cluster-then-SVM model | Cluster 2: cluster-then-RF model | Cluster 3: cluster-then-RF model | Cluster 4: cluster-then-SVM model |
|---|---|---|---|

| | | | |
|---|---|---|---|
| HOMO energy | Number of Chlorine atoms | Number of triple bonds | Number of double bonds |
| Molecular surface area | Number of halogen atoms | Molecular mass | Number of Nitrogen atoms |
| Number of Sulfur atoms | Number of Oxygen atoms | Number of Phosphorus atoms | HOMO energy |
| Connectivity index chi-5 | Molecular mass | Number of Oxygen atoms | Number of triple bonds |
| Connectivity index chi-3 | Number of bonds | Number of halogen atoms | Electric dipole moment |

**Table S5**- Predicted $CF_{ET}$ for the common compounds of the two databases with NA $CF_{ET}$ in USEtox. The unit is the USEtox one.

| CAS | Name | Cluster | Predicted $CF_{ET}$ | Lower bound of the prediction intervals | Upper bound of the prediction intervals |
|---|---|---|---|---|---|
| 191-24-2 | Benzo[g,h,i]perylene | 4 | 176978 | 164318 | 187562 |
| 193-39-5 | Indeno[1,2,3-cd]-pyrene | 4 | 176978 | 164318 | 187562 |
| 205-99-2 | Benzo[b]fluoranthene | 4 | 176846 | 164198 | 187499 |
| 207-08-9 | Benzo[k]fluoranthene | 4 | 176896 | 164244 | 187523 |
| 218-01-9 | Chrysene | 2 | 25996 | 14315 | 28110 |
| 22071-15-4 | Ketoprofen | 2 | 5318 | 4395 | 6023 |
| 25812-30-0 | Gemfibrozil | 2 | 13174 | 12189 | 16126 |
| 439-14-5 | Diazepam | 2 | 4687 | 4545 | 7272 |
| 50-78-2 | Acetylsalicylic acid | 3 | 451 | 399 | 542 |
| 51481-61-9 | Cimetidine | 2 | 5371 | 4304 | 5863 |
| 54-31-9 | Furosemide | 2 | 23463 | 20978 | 30042 |
| 57-41-0 | Phenytoin | 2 | 4109 | 2941 | 4368 |
| 57-68-1 | Sulfamethazine | 2 | 5177 | 4826 | 6983 |
| 72-33-3 | Mestranol | 4 | 178155 | 165342 | 188398 |

| | | | | | |
|---|---|---|---|---|---|
| 73590-58-6 | Omeprazole | 2 | 6781 | 4992 | 7587 |

**Table S6**- Quantiles for the different methods and clusters for the predictions of the $CF_{HT}$. These values are used to obtain the Figure 3. The best models are in bold for each cluster.

| quantiles | RF | PLS | SVM | Cluster-RF | Cluster-SVM | Cluster-PLS |
|---|---|---|---|---|---|---|
| | | | | **Cluster 1** | | |
| 0% | 0,00 | 0,02 | 0,00 | **0,00** | 0,03 | 0,02 |
| 25% | 0,26 | 0,40 | 0,36 | **0,11** | 0,25 | 0,61 |
| 50% | 0,52 | 0,67 | 0,80 | **0,46** | 0,76 | 1,09 |
| 75% | 0,96 | 1,13 | 1,22 | **1,19** | 1,34 | 1,30 |
| 100% | 2,26 | 5,82 | 2,39 | **2,18** | 2,20 | 4,97 |
| | | | | **Cluster 2** | | |
| 0% | 0,00 | 0,01 | **0,00** | 0,00 | 0,00 | 0,00 |
| 25% | 0,38 | 0,34 | **0,36** | 0,38 | 0,38 | 0,40 |
| 50% | 0,75 | 0,79 | **0,75** | 0,81 | 0,79 | 0,85 |
| 75% | 1,40 | 1,43 | **1,37** | 1,50 | 1,43 | 1,69 |
| 100% | 7,23 | 9,90 | **13,33** | 7,79 | 8,55 | 10,20 |
| | | | | **Cluster 3** | | |
| 0% | 0,00 | 0,00 | **0,00** | 0,00 | 0,00 | 0,00 |
| 25% | 0,32 | 0,34 | **0,30** | 0,32 | 0,36 | 0,37 |
| 50% | 0,81 | 0,90 | **0,75** | 0,82 | 0,83 | 0,79 |
| 75% | 1,80 | 1,69 | **1,57** | 1,87 | 1,65 | 1,87 |
| 100% | 5,49 | 5,61 | **5,99** | 6,22 | 8,18 | 8,92 |
| | | | | **Cluster 4** | | |
| 0% | 0,00 | 0,12 | **0,00** | 0,01 | 0,00 | 0,00 |
| 25% | 0,67 | 1,21 | **0,34** | 0,73 | 0,74 | 0,90 |
| 50% | 1,34 | 1,85 | **0,82** | 1,79 | 1,39 | 1,78 |
| 75% | 2,99 | 2,65 | **1,92** | 3,39 | 2,61 | 3,85 |
| 100% | 8,87 | 8,72 | **9,45** | 9,01 | 9,79 | 14,77 |

**Table S7**- Five most important molecular descriptors for each best model for each cluster for the $CF_{HT}$. Descriptors are listed from top to bottom in decreasing order of importance.

| Cluster 1 : cluster-then-RF model | Cluster 2, 3, 4 and 5: SVM model |
|---|---|
| Number of Fluorine atoms | Number of halogen atoms |
| Connectivity index chi-5 | Electric dipole moment |
| Connectivity index chi-1 | Number of double bonds |
| Number of circuits | Number of Chloride atoms |

| Number of rings | Number of Oxygen atoms |
|---|---|

**Table S8**- Predicted $CF_{HT}$ for the common compounds without a $CF_{HT}$ value. The predicted $CF_{HT}$ are rounded at two decimal digits (in USEtox unit).

| CAS | Name | Cluster | Predicted $CF_{HT}$ | Lower bound for the prediction intervals | Upper bound for the prediction intervals |
|---|---|---|---|---|---|
| 101-20-2 | Triclocarban | 2 | 2.3E-04 | 2.0E-04 | 2.4E-04 |
| 101-42-8 | Fenuron | 3 | 2.5E-05 | 1.9E-05 | 3.2E-05 |
| 101205-02-1 | Cycloxydim | 2 | 8.6E-06 | 6.9E-06 | 1.2E-05 |
| 102851-06-9 | tau-Fluvalinate | 1 | 4.7E-05 | 3.1E-05 | 1.1E-04 |
| 1031-07-08 | Endosulfan sulfate | 4 | 1.4E-03 | 1.1E-03 | 1.6E-03 |
| 103361-09-7 | Flumioxazin | 1 | 2.1E-05 | 1.7E-05 | 2.7E-05 |
| 104-40-5 | P-nonylphenol | 2 | 3.7E-06 | 3.2E-06 | 5.3E-06 |
| 10540-29-1 | Tamoxifen | 1 | 1.0E-04 | 2.3E-05 | 1.1E-04 |
| 105512-06-9 | Clodinafop-propargyl | 2 | 4.9E-05 | 4.3E-05 | 5.6E-05 |
| 106-44-5 | P-cresol | 3 | 1.7E-06 | 1.2E-06 | 2.1E-06 |
| 108-62-3 | Metaldehyde (tetramer) | 3 | 5.5E-06 | 4.8E-06 | 6.7E-06 |
| 110488-70-5 | Dimethomorph | 1 | 2.0E-05 | 1.5E-05 | 2.8E-05 |
| 111479-05-1 | Propaquizafop | 1 | 2.9E-05 | 1.8E-05 | 6.2E-05 |
| 111991-09-4 | Nicosulfuron | 1 | 2.7E-05 | 1.9E-05 | 2.9E-05 |
| 114-07-08 | Erythromycin | 5 | 1.8E-04 | 1.5E-04 | 2.2E-04 |
| 117-84-0 | Di(n-octyl) phthalate | 1 | 9.9E-06 | 7.7E-06 | 2.4E-05 |
| 119446-68-3 | Difenoconazole | 1 | 3.3E-05 | 2.0E-05 | 4.3E-05 |
| 1214-39-7 | 1h-purin-6-amine, n-(phenylmethyl) | 2 | 6.2E-06 | 5.5E-06 | 7.7E-06 |
| 121552-61-2 | Cga 219417 (Cyprodinil) | 2 | 2.0E-05 | 1.8E-05 | 2.4E-05 |
| 128639-02-1 | Carfentrazone-ethyl | 2 | 8.3E-05 | 6.6E-05 | 9.9E-05 |
| 131-11-3 | Dimethylphthalate (DMP) | 3 | 2.0E-06 | 1.9E-06 | 2.2E-06 |
| 131341-86-1 | Fludioxonil | 2 | 1.7E-05 | 1.5E-05 | 2.0E-05 |
| 131860-33-8 | Azoxystrobin | 1 | 7.0E-05 | 3.6E-05 | 8.2E-05 |

| | | | | | |
|---|---|---|---|---|---|
| 135158-54-2 | Cga 245704 | 3 | 2.1E-06 | 2.0E-06 | 2.5E-06 |
| 13684-56-5 | Desmedipham | 2 | 1.0E-05 | 9.9E-06 | 1.2E-05 |
| 140-66-9 | P-(1,1,3,3-tetramethylbutyl)phenol | 2 | 3.9E-06 | 3.2E-06 | 5.4E-06 |
| 142459-58-3 | Fluthiamide | 2 | 2.9E-05 | 2.3E-05 | 3.3E-05 |
| 15545-48-9 | Chlortoluron | 3 | 6.4E-06 | 5.7E-06 | 7.2E-06 |
| 1563-38-8 | carbofuran phenol | 3 | 3.0E-06 | 2.5E-06 | 3.8E-06 |
| 1570-64-5 | 2-methyl-4-chlorophenol | 3 | 9.8E-07 | 7.9E-07 | 1.3E-06 |
| 15972-60-8 | Alachlor | 2 | 7.0E-06 | 6.4E-06 | 9.3E-06 |
| 16118-49-3 | Carbetamide | 2 | 2.5E-06 | 2.3E-06 | 2.9E-06 |
| 1698-60-8 | Chloridazon | 3 | 6.4E-06 | 5.9E-06 | 7.3E-06 |
| 1702-17-6 | 3,6-dichloropicolinic acid | 3 | 5.4E-06 | 4.9E-06 | 6.1E-06 |
| 173584-44-6 | Dpx-mp062 | 1 | 4.6E-05 | 2.7E-05 | 1.0E-04 |
| 1746-81-2 | Monolinuron | 3 | 4.8E-06 | 4.4E-06 | 5.4E-06 |
| 2050-68-2 | PCB-15 | 3 | 7.5E-05 | 5.8E-05 | 8.6E-05 |
| 2051-60-7 | PCB-1 | 3 | 1.4E-05 | 1.2E-05 | 1.7E-05 |
| 2051-61-8 | PCB-2 | 3 | 1.4E-05 | 1.1E-05 | 1.6E-05 |
| 26225-79-6 | Ethofumesate | 2 | 6.1E-06 | 5.5E-06 | 7.4E-06 |
| 26761-40-0 | Diisodecyl phthalate | 1 | 1.2E-05 | 8.1E-06 | 4.0E-05 |
| 26787-78-0 | Amoxicillin | 1 | 1.5E-05 | 1.2E-05 | 2.3E-05 |
| 27304-13-8 | Oxychlordane | 4 | 4.6E-02 | 4.1E-02 | 4.9E-02 |
| 28553-12-0 | Diisononyl phthalate | 1 | 1.5E-05 | 1.0E-05 | 4.5E-05 |
| 3060-89-7 | Metobromuron | 3 | 8.5E-06 | 7.9E-06 | 9.2E-06 |
| 33284-50-3 | PCB-7 | 3 | 5.2E-05 | 4.1E-05 | 6.0E-05 |
| 33629-47-9 | Butralin | 2 | 2.3E-06 | 2.2E-06 | 2.8E-06 |
| 3380-34-5 | 5-chloro-2-(2,4-dichlorophenoxy)phenol | 2 | 2.2E-04 | 1.8E-04 | 2.4E-04 |
| 34123-59-6 | Isoproturon | 3 | 3.2E-06 | 2.8E-06 | 3.9E-06 |
| 34256-82-1 | Acetochlor | 2 | 6.8E-06 | 6.1E-06 | 9.1E-06 |

| | | | | | |
|---|---|---|---|---|---|
| 34883-43-7 | 2,4'-dichlorobiphenyl | 3 | 4.8E-05 | 3.8E-05 | 5.5E-05 |
| 3739-38-6 | M-phenoxybenzoic acid | 2 | 6.5E-04 | 5.2E-04 | 7.3E-04 |
| 40321-76-4 | 1,2,3,7,8-pentachlorodibenzo-p-dioxin | 4 | 1.3E-02 | 1.1E-02 | 1.4E-02 |
| 41394-05-02 | Metamitron | 3 | 4.0E-06 | 3.6E-06 | 4.7E-06 |
| 41483-43-6 | Bupirimate | 2 | 3.6E-06 | 3.4E-06 | 4.4E-06 |
| 42835-25-6 | Flumequine | 2 | 1.1E-05 | 9.9E-06 | 1.3E-05 |
| 465-73-6 | Isodrin | 4 | 1.7E-02 | 1.4E-02 | 1.8E-02 |
| 481-39-0 | 5-hydroxy-1,4-naphthoquinone | 3 | 2.8E-06 | 2.4E-06 | 3.2E-06 |
| 51207-31-9 | 2,3,7,8-TetraCDF | 2 | 3.4E-03 | 2.8E-03 | 3.8E-03 |
| 51338-27-3 | Diclofop-methyl | 2 | 6.7E-05 | 5.9E-05 | 7.4E-05 |
| 518-47-8 | Fluorescein sodium | 2 | 2.3E-04 | 2.0E-04 | 2.6E-04 |
| 52888-80-9 | Prosulfocarb | 2 | 4.6E-06 | 4.1E-06 | 6.1E-06 |
| 53-16-7 | Estrone | 4 | 7.1E-05 | 5.9E-05 | 9.1E-05 |
| 53112-28-0 | Pyrimethanil | 3 | 8.3E-06 | 6.9E-06 | 1.0E-05 |
| 55335-06-03 | Triclopyr | 3 | 2.4E-05 | 2.2E-05 | 2.8E-05 |
| 555-37-3 | Neburon | 2 | 1.1E-05 | 9.8E-06 | 1.2E-05 |
| 57-62-5 | Aureomycin | 1 | 2.9E-05 | 2.0E-05 | 8.7E-05 |
| 57653-85-7 | 1,2,3,6,7,8-hexachlorodibenzo-p-dioxin | 4 | 3.0E-02 | 2.3E-02 | 3.2E-02 |
| 57966-95-7 | Cymoxanil | 3 | 4.2E-06 | 3.9E-06 | 4.5E-06 |
| 5915-41-3 | Terbuthylazine | 3 | 5.9E-06 | 5.3E-06 | 6.8E-06 |
| 60-54-8 | Tetracycline | 1 | 2.9E-05 | 2.1E-05 | 8.3E-05 |
| 61213-25-0 | Flurochloridone | 2 | 7.4E-05 | 6.5E-05 | 8.6E-05 |
| 62924-70-3 | Flumetralin | 1 | 3.4E-05 | 2.0E-05 | 3.9E-05 |
| 64-19-7 | Acetic acid | 3 | 4.7E-06 | 3.2E-06 | 5.4E-06 |
| 67129-08-02 | Metazachlor | 2 | 1.4E-05 | 1.3E-05 | 1.6E-05 |
| 67564-91-4 | Fenpropimorph | 2 | 1.1E-05 | 9.4E-06 | 1.6E-05 |

| | | | | | |
|---|---|---|---|---|---|
| 68-35-9 | Sulfadiazine | 2 | 2.6E-06 | 2.4E-06 | 3.0E-06 |
| 69-53-4 | Ampicillin | 2 | 1.2E-05 | 1.1E-05 | 1.4E-05 |
| 69377-81-7 | Fluroxypyr | 3 | 1.8E-05 | 1.6E-05 | 2.1E-05 |
| 70630-17-0 | Metalaxyl-M | 2 | 2.9E-06 | 2.6E-06 | 3.7E-06 |
| 731-27-1 | Tolyfluanide | 2 | 2.9E-05 | 2.5E-05 | 3.1E-05 |
| 74070-46-5 | Aclonifen | 2 | 7.6E-06 | 7.1E-06 | 8.7E-06 |
| 77732-09-03 | Oxadixyl | 2 | 2.6E-06 | 2.4E-06 | 3.0E-06 |
| 79-57-2 | Oxytetracylcine | 1 | 2.5E-05 | 2.0E-05 | 8.0E-05 |
| 79-94-7 | 2,2-bis(4-hydroxy-3,5-Dibromophenyl)propane | 2 | 8.0E-04 | 6.0E-04 | 8.9E-04 |
| 79127-80-3 | Fenoxycarb | 2 | 8.1E-06 | 7.6E-06 | 9.8E-06 |
| 79622-59-6 | Fluazinam | 1 | 4.7E-05 | 2.8E-05 | 6.0E-05 |
| 81777-89-1 | Clomazone | 2 | 1.4E-06 | 1.2E-06 | 1.8E-06 |
| 83164-33-4 | Diflufenican | 1 | 5.7E-05 | 2.9E-05 | 6.4E-05 |
| 834-12-8 | Ametryne | 3 | 6.7E-06 | 6.3E-06 | 7.9E-06 |
| 85-41-6 | Phthalimide | 3 | 1.6E-06 | 1.4E-06 | 1.9E-06 |
| 87392-12-9 | S-Metolachlor | 2 | 8.6E-06 | 7.7E-06 | 1.2E-05 |
| 87674-68-8 | Dimethenamid | 2 | 9.8E-06 | 8.9E-06 | 1.2E-05 |
| 88-99-3 | O-phthalic acid | 3 | 1.9E-06 | 1.8E-06 | 2.1E-06 |
| 90717-03-06 | Quinmerac | 2 | 4.5E-06 | 4.3E-06 | 5.2E-06 |
| 91465-08-06 | Lambda-cyhalothrin | 1 | 6.8E-05 | 2.9E-05 | 9.0E-05 |
| 93106-60-6 | Enrofloxacin | 1 | 1.9E-05 | 1.4E-05 | 2.6E-05 |
| 94125-34-5 | Prosulfuron | 1 | 2.9E-05 | 1.9E-05 | 3.3E-05 |
| 94361-06-05 | Cyproconazole | 2 | 1.4E-05 | 1.4E-05 | 1.7E-05 |
| 95-76-1 | 3,4-dichloroaniline | 3 | 5.0E-06 | 4.0E-06 | 5.9E-06 |
| 97-23-4 | Phenol,2,2'-methylenebis 4-chloro | 2 | 1.1E-04 | 9.0E-05 | 1.2E-04 |
| 99607-70-2 | Cloquintocet-mexyl | 2 | 2.0E-05 | 1.8E-05 | 2.4E-05 |
