## Supplementary material for "Machine learning models based on molecular descriptors to predict human and environmental toxicological factors in continental freshwater": Scripts and data: ReadMe.txt

###script.R
It gathers all the script needed to find the results of the paper, with the R software. 
Warning : some of the computations could last several hours.
Needs the 3 other files as input.
Produce the plots and the tables of the paper as output.

##TyPol_2020_07_23.Rdata
Extraction from the TyPol database, contains all the molecular descriptor.

##usetox.csv
Contains the data used in the paper from the USEtox database.
The first column is the CAS number of the compound.
The second column is the DALY of the compound.
The third column is the PDF of the compound.

##usetox_name.csv
Correspondance between the CAS number of the compound of the usetox database and their name.
