## Supplementary material for "Machine learning models based on molecular descriptors to predict human and environmental toxicological factors in continental freshwater": Scripts and data: script.R

```
rm(list = ls())

#load of the TyPol dataset
load("TyPol_2020_07_23.Rdata")
nrow(results$X) 

#First results of subsection 3.1
Typol_sum<-matrix(0,ncol=6,nrow=40)
rownames(Typol_sum)<-colnames(results$X)
colnames(Typol_sum)<-c("Min global","Max global","Nb NA (%)","Min commun","Max commun","Nb NA commun")
Typol_sum[,1]<-apply(results$X,2,min,na.rm=T)
Typol_sum[,2]<-apply(results$X,2,max,na.rm=T)
Typol_sum[,3]<-colSums(is.na(results$X))/nrow(results$X)*100

#load of usetox database
usetox<-read.csv("usetox.csv", header=F,sep=";",na.strings =c("n/a"))
usetox<-usetox[1:3077,]
colnames(usetox)<-c("CAS","DALY","PDF")
rownames(usetox)<-usetox$CAS

descriptors<-results$X
colnames(descriptors)<-c("C0","C1","C2","C3","C4","C5","Elec_dip_mom","HOMO","LUMO",  "Molecular_mass" ,"Molec_surface_area", "Nb_C", "Nb_Chl"   ,       "Nb_Fluo"     , "Nb_H"    ,      "Nb_Nitro"         , "Nb_O"           , "Nb_Phos"       ,"Nb_S"            ,"Nb_aromatic_bonds"         , "Nb_atoms"                   ,"Nb_bonds"                  ,"Nb_circuits"                ,"Nb_double_bonds"           , "Nb_halogen"           ,"Nb_multiple_bonds"         ,"Nb_non_H_atoms"             ,"Nb_non_H_bonds"            , "Nb_rings"                   ,"Nb_rotable_bonds"          ,"Nb_triple_bonds"            ,"Pola"                   ,"Sum_of_conv"    ,"Total_energy"                     ,"V0"  ,"V1" , "V2"  ,"V3", "V4"  ,"V5" )

#we keep the common compounds
common_U_X_2101<-intersect(rownames(usetox),rownames(descriptors)) 
usetox_keep<-usetox[common_U_X_2101,]
descriptors_keep<-descriptors[common_U_X_2101,]

Typol_sum[,4]<-apply(descriptors_keep,2,min,na.rm=T)
Typol_sum[,5]<-apply(descriptors_keep,2,max,na.rm=T)
Typol_sum[,6]<-colSums(is.na(descriptors_keep))/nrow(descriptors_keep)*100
Typol_sum<-round(Typol_sum,digits=2)

write.csv(Typol_sum,file="Typol_summary.csv")

#load of the name of the common compounds
usetox_name<-read.csv("usetox_name.csv", header=T,sep=";",na.strings =c("n/a"))
colnames(usetox_name)[1]<-"CAS"
usetox_keep[,3]<-round(usetox_keep[,3],digits=1)
common_name<-merge(usetox_name,usetox_keep,by="CAS")
common_name$DALY<-signif(common_name$DALY,digits=2)

write.csv(common_name,file="Common_name.csv")

sum(is.na(common_name$DALY))
sum(is.na(common_name$PDF))

#boxplots
boxplot(log10(usetox$PDF),log10(usetox_keep$PDF),names=c("USEtox","USEtox & TyPol"),ylab=expression(CF[ET]))
dev.print(png, file = "boxplot_PDF.png", width = 1600, height = 1500, res = 150)

eps<-1e-10
boxplot(log10(usetox$DALY+eps),log10(usetox_keep$DALY+eps),names=c("USEtox","USEtox & TyPol"),ylab=expression(CF[HT]))
dev.print(png, file = "boxplot_DALY.png", width = 1600, height = 1500, res = 150)

#TyPol clustering graph
library(mixOmics)

head(results$analyse$plsComp) ##coordinate of the individuals
classif_commun_axes<-results$analyse$plsComp[common_U_X_2101,]
rownames(results$analyse$classes_mol)<-results$analyse$classes_mol[,1]
classes_commun<-results$analyse$classes_mol[common_U_X_2101,]

#modifications of the colors for the graph
classes_commun<-cbind(classes_commun,classes_commun[,2])
colnames(classes_commun)[3]<-"Graph_color"
classes_commun[classes_commun[,2]=="1",3]<-"black"
classes_commun[classes_commun[,2]=="2",3]<-"red"
classes_commun[classes_commun[,2]=="3",3]<-"green"
classes_commun[classes_commun[,2]=="4",3]<-"blue"
classes_commun[classes_commun[,2]=="5",3]<-"brown"

plot(0,0,type="n",xlim=c(min(classif_commun_axes[,1]),max(classif_commun_axes[,1])),
     ylim=c(min(classif_commun_axes[,2]),max(classif_commun_axes[,2])),xlab="First axis",ylab="Second axis")
points(classif_commun_axes[,1],classif_commun_axes[,2],cex=0.4,col=classes_commun[,3])
text(classif_commun_axes[,1],classif_commun_axes[,2],labels=rownames(classif_commun_axes),pos=3,cex=.5,col=classes_commun[,3])
legend("bottomright",col=c("black","red","green","blue","brown"),lty=rep(1,5),legend=c("Cluster 1","Cluster 2","Cluster 3","Cluster 4","Cluster 5"))
dev.print(png, file = "clusters_common.png", width = 1600, height = 1500, res = 150)

#graph for all the compounds
#colors
head(results$analyse$classes_mol)
classes_tout<-cbind(results$analyse$classes_mol,results$analyse$classes_mol[,2])
colnames(classes_tout)[3]<-"Graph_color"

classes_tout[classes_tout[,2]=="1",3]<-"black"
classes_tout[classes_tout[,2]=="2",3]<-"red"
classes_tout[classes_tout[,2]=="3",3]<-"green"
classes_tout[classes_tout[,2]=="4",3]<-"blue"
classes_tout[classes_tout[,2]=="5",3]<-"brown"

plot(0,0,type="n",xlim=c(min(results$analyse$plsComp[,1]),max(results$analyse$plsComp[,1])),
     ylim=c(min(results$analyse$plsComp[,2]),max(results$analyse$plsComp[,2])),xlab="First axis",ylab="Second axis")
points(results$analyse$plsComp[,1],results$analyse$plsComp[,2],cex=0.4,col=classes_tout[,3])
text(results$analyse$plsComp[,1],results$analyse$plsComp[,2],labels=rownames(results$analyse$plsComp),pos=3,cex=.5,col=classes_tout[,3])
legend("bottomright",col=c("black","red","green","blue","brown"),lty=rep(1,5),legend=c("Cluster 1","Cluster 2","Cluster 3","Cluster 4","Cluster 5"))
dev.print(png, file = "clusters_tout.png", width = 1600, height = 1500, res = 150)

#boxplots PDF DALY per cluster
classes_PDF_DALY<-merge(usetox_keep,classes_commun,by="row.names")
colnames(classes_PDF_DALY)[6]<-"Cluster"
par(mfrow=c(2,1))
par(mgp=c(3,1.5,0))
boxplot(log10(PDF)~Cluster,data=classes_PDF_DALY,ylab=expression(CF[ET]),names=c("Cluster 1 \n n=33 without NA","Cluster 2 \n n=122 with 9 NA","Cluster 3 \n n=91 with 1 NA","Cluster 4 \n n=27 with 5 NA","Cluster 5 \n n=1 without NA"))
boxplot(log10(DALY+eps)~Cluster,data=classes_PDF_DALY,ylab=expression(CF[HT]),names=c("Cluster 1 \n n=33 with 22 NA","Cluster 2 \n n=122 with 43 NA","Cluster 3 \n n=91 with 30 NA","Cluster 4 \n n=27 with 6 NA","Cluster 5 \n n=1 with 1 NA"))
dev.print(png, file = "boxplot_PDF_DALY_clusters.png", width = 1600, height = 1500, res = 150)

#boxplots of molecular descriptors per clusters
colnames(classes_commun)<-c("CAS","Gpe","col")
boxxx<-merge(descriptors_keep,classes_commun,by="row.names")
par(mfrow=c(4,2))
for (i in 2:9)
{boxplot(boxxx[,i]~boxxx$Gpe,main=colnames(boxxx)[i],names=c("Cluster 1","Cluster 2","Cluster 3","Cluster 4","Cluster 5"))}
dev.print(png, file = "boxplot_desc1.png", width = 1600, height = 1500, res = 150)
for (i in 10:17)
{boxplot(boxxx[,i]~boxxx$Gpe,main=colnames(boxxx)[i],names=c("Cluster 1","Cluster 2","Cluster 3","Cluster 4","Cluster 5"))}
dev.print(png, file = "boxplot_desc2.png", width = 1600, height = 1500, res = 150)
for (i in 18:25)
{boxplot(boxxx[,i]~boxxx$Gpe,main=colnames(boxxx)[i],names=c("Cluster 1","Cluster 2","Cluster 3","Cluster 4","Cluster 5"))}
dev.print(png, file = "boxplot_desc3.png", width = 1600, height = 1500, res = 150)
for (i in 26:33)
{boxplot(boxxx[,i]~boxxx$Gpe,main=colnames(boxxx)[i],names=c("Cluster 1","Cluster 2","Cluster 3","Cluster 4","Cluster 5"))}
dev.print(png, file = "boxplot_desc4.png", width = 1600, height = 1500, res = 150)
for (i in 34:41)
{boxplot(boxxx[,i]~boxxx$Gpe,main=colnames(boxxx)[i],names=c("Cluster 1","Cluster 2","Cluster 3","Cluster 4","Cluster 5"))}
dev.print(png, file = "boxplot_desc5.png", width = 1600, height = 1500, res = 150)

table(classes_PDF_DALY$Cluster)

Nb_PDF<-rep(0,5)
Nb_DALY<-rep(0,5)
for (i in 1:nrow(classes_PDF_DALY))
{ if (!(is.na(classes_PDF_DALY$DALY[i]))){Nb_DALY[classes_PDF_DALY$Cluster[i]]<-Nb_DALY[classes_PDF_DALY$Cluster[i]]+1}
  if (!(is.na(classes_PDF_DALY$PDF[i]))){Nb_PDF[classes_PDF_DALY$Cluster[i]]<-Nb_PDF[classes_PDF_DALY$Cluster[i]]+1}
}
Nb_PDF
sum(Nb_PDF)
Nb_DALY
sum(Nb_DALY)

#Results of Section 3.2
#independant script from above
rm(list = ls())

set.seed(150720)
#load of the datasets
usetox<-read.csv("usetox.csv", header=F,sep=";",na.strings =c("n/a"))
usetox<-usetox[1:3077,]
colnames(usetox)<-c("CAS","DALY","PDF")
rownames(usetox)<-usetox$CAS

load("TyPol_2020_07_23.Rdata")
descriptors<-results$X
colnames(descriptors)<-c("C0","C1","C2","C3","C4","C5","Elec_dip_mom","HOMO","LUMO",  "Molecular_mass" ,"Molec_surface_area", "Nb_C", "Nb_Chl"   ,       "Nb_Fluo"     , "Nb_H"    ,      "Nb_Nitro"         , "Nb_O"           , "Nb_Phos"       ,"Nb_S"            ,"Nb_aromatic_bonds"         , "Nb_atoms"                   ,"Nb_bonds"                  ,"Nb_circuits"                ,"Nb_double_bonds"           , "Nb_halogen"           ,"Nb_multiple_bonds"         ,"Nb_non_H_atoms"             ,"Nb_non_H_bonds"            , "Nb_rings"                   ,"Nb_rotable_bonds"          ,"Nb_triple_bonds"            ,"Pola"                   ,"Sum_of_conv"    ,"Total_energy"                     ,"V0"  ,"V1" , "V2"  ,"V3", "V4"  ,"V5" )

#keep only common compounds
common_U_X_2101<-intersect(rownames(usetox),rownames(descriptors)) 
usetox_keep<-usetox[common_U_X_2101,]
descriptors_keep<-descriptors[common_U_X_2101,]

head(results$analyse$classes_mol) ##classif
rownames(results$analyse$classes_mol)<-results$analyse$classes_mol[,1]
classes_commun<-results$analyse$classes_mol[common_U_X_2101,]

#create a dataset without NA for train/test
length(usetox_keep$DALY)##274
sum(is.na(usetox_keep$DALY)==F)##172 
usetox_keep_DALY_sans_NA<-usetox_keep[is.na(usetox_keep$DALY)==F,]

table(classes_commun[,2])
usetox_keep_DALY_sans_NA_c<-merge(usetox_keep_DALY_sans_NA,classes_commun,by="row.names")
table(usetox_keep_DALY_sans_NA_c$V2) 

#train/test procedure
library(randomForest)
library(e1071)

epsilon<-1e-10
Error<-matrix(NA,ncol=7)
colnames(Error)<-c("RF_pred","PLS_pred","SVM_pred","Gpe_RF_pred","Gpe_SVM_pred","Gpe_PLS_pred","Gpe")
nbboot<-200
if (!file.exists("Error_DALY.rds")) ###could be very long 
{for (tt in (1:nbboot))
{ print(tt)  
#Train/test
#15% in each cluster : 1 in cluster 1, 12 in 2, 9 in 3, 3 in 4
  gpe1<-usetox_keep_DALY_sans_NA_c[usetox_keep_DALY_sans_NA_c$V2=="1",]
  gpe2<-usetox_keep_DALY_sans_NA_c[usetox_keep_DALY_sans_NA_c$V2=="2",]
  gpe3<-usetox_keep_DALY_sans_NA_c[usetox_keep_DALY_sans_NA_c$V2=="3",]
  gpe4<-usetox_keep_DALY_sans_NA_c[usetox_keep_DALY_sans_NA_c$V2=="4",]
  sample1<-sample(c(1:nrow(gpe1)),size=1)
  sample2<-sample(c(1:nrow(gpe2)),size=12)
  sample3<-sample(c(1:nrow(gpe3)),size=9)
  sample4<-sample(c(1:nrow(gpe4)),size=3)
  Valid1<-gpe1[sample1,]
  Valid2<-gpe2[sample2,]
  Valid3<-gpe3[sample3,]
  Valid4<-gpe4[sample4,]

#we create the test set
  usetox_keep_DALY_test<-rbind(Valid1,Valid2,Valid3,Valid4)
  rownames(usetox_keep_DALY_test)<-usetox_keep_DALY_test$CAS
  usetox_keep_DALY_test<-usetox_keep_DALY_test[,c(2,3,4,6)]
  colnames(usetox_keep_DALY_test)[4]<-"Cluster"

#train set
#we remove the compounds that are in the test set
  usetox_keep_DALY_train<-usetox_keep_DALY_sans_NA
  for (i in (1:nrow(usetox_keep_DALY_test)))
    {usetox_keep_DALY_train<-usetox_keep_DALY_train[-which(usetox_keep_DALY_train$CAS==usetox_keep_DALY_test$CAS[i]),]}

#put descriptors and PDF in the same matrix
  train_total<-merge(usetox_keep_DALY_train,descriptors,by="row.names")
  test_total<-merge(usetox_keep_DALY_test,descriptors,by="row.names")
  train_total$log10DALY<-log10(train_total$DALY+epsilon)
  test_total$log10DALY<-log10(test_total$DALY+epsilon)

#modelisation
#1. Nipals for NA in train
  pls_nipals_train<-nipals(X=train_total[,5:44], ncomp = 10, reconst = TRUE, max.iter = 500, tol = 1e-09)
  data_nipals<-cbind(pls_nipals_train$rec,train_total$log10DALY)
  colnames(data_nipals)[ncol(data_nipals)]<-"log10DALY"
  rownames(data_nipals)<-train_total$Row.names

  ff<-as.formula(log10DALY~C0+C1+C2+C3+C4+C5+Elec_dip_mom+HOMO+LUMO+Molecular_mass+Molec_surface_area+Nb_C+Nb_Chl+Nb_Fluo+Nb_H+Nb_Nitro+Nb_O+Nb_Phos+Nb_S+Nb_aromatic_bonds+Nb_atoms+Nb_bonds+Nb_circuits+Nb_double_bonds+Nb_halogen+Nb_multiple_bonds+Nb_non_H_atoms+Nb_non_H_bonds+Nb_rings+Nb_rotable_bonds+Nb_triple_bonds+Pola+Sum_of_conv+Total_energy+V0+V1+V2+V3+V4+V5)
#2. rf
  tune.RF<-tune.randomForest(x=pls_nipals_train$rec,y=train_total$log10DALY,mtry=c(3:30))
  tune.RF$best.parameters 
  model_RF_DALY<-randomForest(ff,data=data_nipals,importance=TRUE,na.action=na.roughfix,mtry=as.integer(tune.RF$best.parameters))

#pls
  model_pre_pls_DALY<-pls(X=pls_nipals_train$rec,Y=train_total$log10DALY,ncomp=10)
  tune.pls <- perf(model_pre_pls_DALY, validation = 'Mfold', folds=10,criterion = 'all', test.keepX=40,measure="MSE")
  model_pls_DALY<-pls(X=pls_nipals_train$rec,Y=train_total$log10DALY,ncomp=which.min(tune.pls$MSEP))

#svm
  tune_svm<-tune.svm(x=pls_nipals_train$rec,y=train_total$log10DALY, gamma = 10^(-3:3), cost = 10^(-3:3))
  model_svm_DALY<-svm(x=pls_nipals_train$rec,y=train_total$log10DALY,gamma=tune_svm$best.parameters[1],cost=tune_svm$best.parameters[2])

#cluster-then-predict approaches(RF, SVM and PLS)
  rownames(gpe1)<-gpe1$Row.names
  Gpe1_train<-merge(gpe1,data_nipals,by="row.names")
  colnames(Gpe1_train)[43:44]<-c("V1","V2")
  tune.RF_gpe1<-tune.randomForest(x=Gpe1_train[,8:47],y=Gpe1_train$log10DALY,mtry=c(3:30))
  tune.RF_gpe1$best.parameters 
  model_RF_DALY_gpe1<-randomForest(x=Gpe1_train[,8:47],y=Gpe1_train$log10DALY,importance=TRUE,na.action=na.roughfix,mtry=as.integer(tune.RF_gpe1$best.parameters))
  tune_svm1<-tune.svm(x=Gpe1_train[,8:47],y=Gpe1_train$log10DALY, gamma = 10^(-3:3), cost = 10^(-3:3))
  model_svm1_DALY<-svm(x=Gpe1_train[,8:47],y=Gpe1_train$log10DALY,gamma=tune_svm1$best.parameters[1],cost=tune_svm1$best.parameters[2])
  model_pre_pls1_DALY<-pls(X=Gpe1_train[,8:47],Y=Gpe1_train$log10DALY,ncomp=8)
  tune.pls1<- perf(model_pre_pls1_DALY, validation = 'Mfold', folds=10,criterion = 'all', test.keepX=40,measure="MSE")
  model_pls1_DALY<-pls(X=Gpe1_train[,8:47],Y=Gpe1_train$log10DALY,ncomp=which.min(tune.pls1$MSEP))

  rownames(gpe2)<-gpe2$Row.names
  Gpe2_train<-merge(gpe2,data_nipals,by="row.names")
  colnames(Gpe2_train)[43:44]<-c("V1","V2")
  tune.RF_gpe2<-tune.randomForest(x=Gpe2_train[,8:47],y=Gpe2_train$log10DALY,mtry=c(3:30))
  tune.RF_gpe2$best.parameters 
  model_RF_DALY_gpe2<-randomForest(x=Gpe2_train[,8:47],y=Gpe2_train$log10DALY,importance=TRUE,na.action=na.roughfix,mtry=as.integer(tune.RF_gpe2$best.parameters))
  tune_svm2<-tune.svm(x=Gpe2_train[,8:47],y=Gpe2_train$log10DALY, gamma = 10^(-3:3), cost = 10^(-3:3))
  model_svm2_DALY<-svm(x=Gpe2_train[,8:47],y=Gpe2_train$log10DALY,gamma=tune_svm2$best.parameters[1],cost=tune_svm2$best.parameters[2])
  model_pre_pls2_DALY<-pls(X=Gpe2_train[,8:47],Y=Gpe2_train$log10DALY,ncomp=8)
  tune.pls2<- perf(model_pre_pls2_DALY, validation = 'Mfold', folds=10,criterion = 'all', test.keepX=40,measure="MSE")
  model_pls2_DALY<-pls(X=Gpe2_train[,8:47],Y=Gpe2_train$log10DALY,ncomp=which.min(tune.pls2$MSEP))

  rownames(gpe3)<-gpe3$Row.names
  Gpe3_train<-merge(gpe3,data_nipals,by="row.names")
  colnames(Gpe3_train)[43:44]<-c("V1","V2")
  tune.RF_gpe3<-tune.randomForest(x=Gpe3_train[,8:47],y=Gpe3_train$log10DALY,mtry=c(3:30))
  tune.RF_gpe3$best.parameters 
  model_RF_DALY_gpe3<-randomForest(x=Gpe3_train[,8:47],y=Gpe3_train$log10DALY,importance=TRUE,na.action=na.roughfix,mtry=as.integer(tune.RF_gpe3$best.parameters))
  tune_svm3<-tune.svm(x=Gpe3_train[,8:47],y=Gpe3_train$log10DALY, gamma = 10^(-3:3), cost = 10^(-3:3))
  model_svm3_DALY<-svm(x=Gpe3_train[,8:47],y=Gpe3_train$log10DALY,gamma=tune_svm3$best.parameters[1],cost=tune_svm3$best.parameters[2])
  model_pre_pls3_DALY<-pls(X=Gpe3_train[,8:47],Y=Gpe3_train$log10DALY,ncomp=8)
  tune.pls3<- perf(model_pre_pls3_DALY, validation = 'Mfold', folds=10,criterion = 'all', test.keepX=40,measure="MSE")
  model_pls3_DALY<-pls(X=Gpe3_train[,8:47],Y=Gpe3_train$log10DALY,ncomp=which.min(tune.pls3$MSEP))

  rownames(gpe4)<-gpe4$Row.names
  Gpe4_train<-merge(gpe4,data_nipals,by="row.names")
  colnames(Gpe4_train)[43:44]<-c("V1","V2")
  tune.RF_gpe4<-tune.randomForest(x=Gpe4_train[,8:47],y=Gpe4_train$log10DALY,mtry=c(3:30))
  tune.RF_gpe4$best.parameters 
  model_RF_DALY_gpe4<-randomForest(x=Gpe4_train[,8:47],y=Gpe4_train$log10DALY,importance=TRUE,na.action=na.roughfix,mtry=as.integer(tune.RF_gpe4$best.parameters))
  tune_svm4<-tune.svm(x=Gpe4_train[,8:47],y=Gpe4_train$log10DALY, gamma = 10^(-3:3), cost = 10^(-3:3))
  model_svm4_DALY<-svm(x=Gpe4_train[,8:47],y=Gpe4_train$log10DALY,gamma=tune_svm4$best.parameters[1],cost=tune_svm4$best.parameters[2])
  model_pre_pls4_DALY<-pls(X=Gpe4_train[,8:47],Y=Gpe4_train$log10DALY,ncomp=8)
  tune.pls4<- perf(model_pre_pls4_DALY, validation = 'Mfold', folds=10,criterion = 'all', test.keepX=40,measure="MSE")
  model_pls4_DALY<-pls(X=Gpe4_train[,8:47],Y=Gpe4_train$log10DALY,ncomp=which.min(tune.pls4$MSEP))

#prediction
  results_DALY<-matrix(NA,ncol=9,nrow=nrow(test_total))
  rownames(results_DALY)<-test_total$Row.names
  colnames(results_DALY)<-c("RF_pred","PLS_pred","SVM_pred","Gpe_RF_pred","Gpe_SVM_pred","Gpe_PLS_pred","Combination_pred","Gpe","Real_value")
  results_DALY[,ncol(results_DALY)]<-test_total$log10DALY

#nipals for test set
  test_nipals<-nipals(X=rbind(train_total[,5:44],test_total[,6:45]), ncomp = 10, reconst = TRUE, max.iter = 500, tol = 1e-09)
  test_total[,6:45]<-test_nipals$rec[(nrow(test_nipals$rec)-nrow(test_total)+1):nrow(test_nipals$rec),]

  results_DALY[,1]<-predict(model_RF_DALY,test_total[,6:45])
  results_DALY[,2]<-predict(model_pls_DALY,test_total[,6:45])$predict[,,which.min(tune.pls$MSEP)]
  results_DALY[,3]<-predict(model_svm_DALY,test_total[,6:45])

#prediction per cluster
  results_DALY[,8]<-merge(results$analyse$classes_mol,results_DALY,by="row.names")$V2
  l4<-which(results_DALY[,8]==4)
  results_DALY[l4,4]<-predict(model_RF_DALY_gpe4,test_total[l4,6:45]) 
  results_DALY[l4,5]<-predict(model_svm4_DALY,test_total[l4,6:45]) 
  results_DALY[l4,6]<-predict(model_pls4_DALY,test_total[l4,6:45])$predict[,,which.min(tune.pls4$MSEP)] 
  l3<-which(results_DALY[,8]==3)
  results_DALY[l3,4]<-predict(model_RF_DALY_gpe3,test_total[l3,6:45])
  results_DALY[l3,5]<-predict(model_svm3_DALY,test_total[l3,6:45]) 
  results_DALY[l3,6]<-predict(model_pls3_DALY,test_total[l3,6:45])$predict[,,which.min(tune.pls3$MSEP)] 
  l2<-which(results_DALY[,8]==2)
  results_DALY[l2,4]<-predict(model_RF_DALY_gpe2,test_total[l2,6:45]) 
  results_DALY[l2,5]<-predict(model_svm2_DALY,test_total[l2,6:45]) 
  results_DALY[l2,6]<-predict(model_pls2_DALY,test_total[l2,6:45])$predict[,,which.min(tune.pls2$MSEP)] 
  l1<-which(results_DALY[,8]==1)
  results_DALY[l1,4]<-predict(model_RF_DALY_gpe1,test_total[l1,6:45]) 
  results_DALY[l1,5]<-predict(model_svm1_DALY,test_total[l1,6:45]) 
  results_DALY[l1,6]<-predict(model_pls1_DALY,test_total[l1,6:45])$predict[,,which.min(tune.pls1$MSEP)] 

#Results for each method
  Error_bcle<-matrix(NA,ncol=7,nrow=nrow(test_total))
  colnames(Error_bcle)<-c("RF_pred","PLS_pred","SVM_pred","Gpe_RF_pred","Gpe_SVM_pred","Gpe_PLS_pred","Gpe")
  Error_bcle[,1]<-results_DALY[,1]-results_DALY[,9]
  Error_bcle[,2]<-results_DALY[,2]-results_DALY[,9]
  Error_bcle[,3]<-results_DALY[,3]-results_DALY[,9]
  Error_bcle[,4]<-results_DALY[,4]-results_DALY[,9]
  Error_bcle[,5]<-results_DALY[,5]-results_DALY[,9]
  Error_bcle[,6]<-results_DALY[,6]-results_DALY[,9]
  Error_bcle[,7]<-results_DALY[,8]
  Error<-rbind(Error,Error_bcle)
}
Error<-Error[-1,]
save(Error,file="Error_DALY.rds")
}
#scatterplot
nbscatterboot<-500
scatter_DALY<-cbind(usetox_keep_DALY_sans_NA,matrix(NA,ncol=nbscatterboot,nrow=nrow(usetox_keep_DALY_sans_NA)))
if (!file.exists("scatter_DALY.rds"))
{for (tt in 1:nbscatterboot)
{ print(tt)
  #Train/test
  gpe1<-usetox_keep_DALY_sans_NA_c[usetox_keep_DALY_sans_NA_c$V2=="1",]
  gpe2<-usetox_keep_DALY_sans_NA_c[usetox_keep_DALY_sans_NA_c$V2=="2",]
  gpe3<-usetox_keep_DALY_sans_NA_c[usetox_keep_DALY_sans_NA_c$V2=="3",]
  gpe4<-usetox_keep_DALY_sans_NA_c[usetox_keep_DALY_sans_NA_c$V2=="4",]
  sample1<-sample(c(1:nrow(gpe1)),size=1)
  sample2<-sample(c(1:nrow(gpe2)),size=12)
  sample3<-sample(c(1:nrow(gpe3)),size=9)
  sample4<-sample(c(1:nrow(gpe4)),size=3)
  Valid1<-gpe1[sample1,]
  Valid2<-gpe2[sample2,]
  Valid3<-gpe3[sample3,]
  Valid4<-gpe4[sample4,]
  
  #test set
  usetox_keep_DALY_test<-rbind(Valid1,Valid2,Valid3,Valid4)
  rownames(usetox_keep_DALY_test)<-usetox_keep_DALY_test$CAS
  usetox_keep_DALY_test<-usetox_keep_DALY_test[,c(2,3,4,6)]
  colnames(usetox_keep_DALY_test)[4]<-"Cluster"
  
  usetox_keep_DALY_train<-usetox_keep_DALY_sans_NA
  for (i in (1:nrow(usetox_keep_DALY_test)))
  {usetox_keep_DALY_train<-usetox_keep_DALY_train[-which(usetox_keep_DALY_train$CAS==usetox_keep_DALY_test$CAS[i]),]}
  
  train_total<-merge(usetox_keep_DALY_train,descriptors,by="row.names")
  test_total<-merge(usetox_keep_DALY_test,descriptors,by="row.names")
  train_total$log10DALY<-log10(train_total$DALY+epsilon)
  test_total$log10DALY<-log10(test_total$DALY+epsilon)
  
  #modelisation
  #1. Nipals
  pls_nipals_train<-nipals(X=train_total[,5:44], ncomp = 10, reconst = TRUE, max.iter = 500, tol = 1e-09)
  data_nipals<-cbind(pls_nipals_train$rec,train_total$log10DALY)
  colnames(data_nipals)[ncol(data_nipals)]<-"log10DALY"
  rownames(data_nipals)<-train_total$Row.names
  
  ff<-as.formula(log10DALY~C0+C1+C2+C3+C4+C5+Elec_dip_mom+HOMO+LUMO+Molecular_mass+Molec_surface_area+Nb_C+Nb_Chl+Nb_Fluo+Nb_H+Nb_Nitro+Nb_O+Nb_Phos+Nb_S+Nb_aromatic_bonds+Nb_atoms+Nb_bonds+Nb_circuits+Nb_double_bonds+Nb_halogen+Nb_multiple_bonds+Nb_non_H_atoms+Nb_non_H_bonds+Nb_rings+Nb_rotable_bonds+Nb_triple_bonds+Pola+Sum_of_conv+Total_energy+V0+V1+V2+V3+V4+V5)
  
  #2. SVM global for cluster 2 3 4, Cluster then RF for cluster 1
  #svm
  tune_svm<-tune.svm(x=pls_nipals_train$rec,y=train_total$log10DALY, gamma = 10^(-3:3), cost = 10^(-3:3))
  model_svm_DALY<-svm(x=pls_nipals_train$rec,y=train_total$log10DALY,gamma=tune_svm$best.parameters[1],cost=tune_svm$best.parameters[2])
  
  #clustering then RF gpe 1
  rownames(gpe1)<-gpe1$Row.names
  Gpe1_train<-merge(gpe1,data_nipals,by="row.names")
  colnames(Gpe1_train)[43:44]<-c("V1","V2")
  tune.RF_gpe1<-tune.randomForest(x=Gpe1_train[,8:47],y=Gpe1_train$log10DALY,mtry=c(3:30))
  tune.RF_gpe1$best.parameters 
  model_RF_DALY_gpe1<-randomForest(x=Gpe1_train[,8:47],y=Gpe1_train$log10DALY,importance=TRUE,na.action=na.roughfix,mtry=as.integer(tune.RF_gpe1$best.parameters))
  
  #3.prediction
  results_DALY<-matrix(NA,ncol=9,nrow=nrow(test_total))
  rownames(results_DALY)<-test_total$Row.names
  colnames(results_DALY)<-c("RF_pred","PLS_pred","SVM_pred","Gpe_RF_pred","Gpe_SVM_pred","Gpe_PLS_pred","Combination_pred","Gpe","Real_value")
  results_DALY[,ncol(results_DALY)]<-test_total$log10DALY
  results_DALY[,8]<-merge(results$analyse$classes_mol,results_DALY,by="row.names")$V2
  #nipals for test
  test_nipals<-nipals(X=rbind(train_total[,5:44],test_total[,6:45]), ncomp = 10, reconst = TRUE, max.iter = 500, tol = 1e-09)
  test_total[,6:45]<-test_nipals$rec[(nrow(test_nipals$rec)-nrow(test_total)+1):nrow(test_nipals$rec),]
  
  results_DALY[,3]<-predict(model_svm_DALY,test_total[,6:45]) ##prediction SVM global
  l1<-which(results_DALY[,8]==1)
  results_DALY[l1,4]<-predict(model_RF_DALY_gpe1,test_total[l1,6:45]) ##cluster RF gpe1
  
  for (jj in 1:nrow(results_DALY))
  {
    if (results_DALY[jj,8]==1) ##cluster-RF
    { scatter_DALY[rownames(results_DALY)[jj],tt+3]<-results_DALY[jj,4]}
    else ##cluster-SVM
    {scatter_DALY[rownames(results_DALY)[jj],tt+3]<-results_DALY[jj,3]}
  }  
save(scatter_DALY,file="scatter_DALY.rds")  
}  
}
load("scatter_DALY.rds")
scatter_DALY<-as.data.frame(scatter_DALY)
scatter_DALY$med_est<-apply(scatter_DALY[4:(nbboot+3)],1,median,na.rm=T)
scatter_DALY$med_est05<-apply(scatter_DALY[4:(nbboot+3)],1,quantile,probs=0.05,na.rm=T)
scatter_DALY$med_est95<-apply(scatter_DALY[4:(nbboot+3)],1,quantile,probs=0.95,na.rm=T)
scatter_DALY$log10DALY<-log10(scatter_DALY$DALY+epsilon)
ggplot(data = scatter_DALY,aes(x=log10DALY, y=med_est,ymin=med_est05,ymax=med_est95))+geom_point()+geom_abline(slope=1,intercept=0)+geom_errorbar()+xlab(expression(CF[HT]))+ylab("Estimation")
dev.print(png, file = "scatter_DALY.png", width = 1600, height = 1500, res = 150)

load("Error_DALY.rds")
colnames(Error)[1:7]<-c("RF","PLS","SVM","Cluster-RF","Cluster-SVM","Cluster-PLS","Gpe")

#general performances
colMeans(abs(Error),na.rm=T)

col_boxplot<-apply(abs(Error),2,median,na.rm=T)

library("RColorBrewer")
palette_col<-rev(brewer.pal(n = 6, name = 'RdYlGn'))
boxplot(abs(Error[,c(1,2,3,4,6,5)]),outline=F,ylab="Absolute Error",col=palette_col[rank(col_boxplot[c(1,2,3,4,6,5)])])
dev.print(png, file = "boxplot_DALY_AE.png", width = 1600, height = 1500, res = 150)

#performances per cluster
col_gpe1<-apply(abs(Error[Error[,7]==1,]),2,median,na.rm=T)
col_gpe2<-apply(abs(Error[Error[,7]==2,]),2,median,na.rm=T)
col_gpe3<-apply(abs(Error[Error[,7]==3,]),2,median,na.rm=T)
col_gpe4<-apply(abs(Error[Error[,7]==4,]),2,median,na.rm=T)

par(mfrow=c(2,2))
boxplot(abs(Error[Error[,7]==1,c(1,2,3,4,6,5)]),ylim=c(0,8),outline=F,ylab="Absolute Error",main="Cluster 1",col=palette_col[rank(col_gpe1[c(1,2,3,4,6,5)])])
boxplot(abs(Error[Error[,7]==2,c(1,2,3,4,6,5)]),ylim=c(0,8),outline=F,ylab="Absolute Error",main="Cluster 2",col=palette_col[rank(col_gpe2[c(1,2,3,4,6,5)])])
boxplot(abs(Error[Error[,7]==3,c(1,2,3,4,6,5)]),ylim=c(0,8),outline=F,ylab="Absolute Error",main="Cluster 3",col=palette_col[rank(col_gpe3[c(1,2,3,4,6,5)])])
boxplot(abs(Error[Error[,7]==4,c(1,2,3,4,6,5)]),ylim=c(0,8),outline=F,ylab="Absolute Error",main="Cluster 4",col=palette_col[rank(col_gpe4[c(1,2,3,4,6,5)])])
dev.print(png, file = "boxplot_DALY_AE_gpe.png", width = 2500, height = 1500, res = 150)

#best local model
usetox_keep_DALY_sans_NA_c<-usetox_keep_DALY_sans_NA_c[,-c(1,5)]
colnames(usetox_keep_DALY_sans_NA_c)[4]<-"Gpe"
rownames(usetox_keep_DALY_sans_NA_c)<-usetox_keep_DALY_sans_NA_c$CAS
total<-merge(usetox_keep_DALY_sans_NA_c,descriptors,by="row.names")
total$log10DALY<-log10(total$DALY+epsilon)
 
# Nipals
library(mixOmics)
nipals_total<-nipals(X=total[,6:45], ncomp = 10, reconst = TRUE, max.iter = 500, tol = 1e-09)
total[,6:45]<-nipals_total$rec

#svm global 
tune_svm_total<-tune.svm(x=total[,6:45],y=total$log10DALY, gamma = 10^(-3:3), cost = 10^(-3:3))
model_svm_DALY_final_for_gpe12345<-svm(x=total[,6:45],y=total$log10DALY,gamma=tune_svm_total$best.parameters[1],cost=tune_svm_total$best.parameters[2])

#SVM description
model_svm_DALY_final_for_gpe12345$index ##point support
length(model_svm_DALY_final_for_gpe12345$index)

#W
t(model_svm_DALY_final_for_gpe12345$coefs)%*%model_svm_DALY_final_for_gpe12345$SV

#by importance order
(t(model_svm_DALY_final_for_gpe12345$coefs)%*%model_svm_DALY_final_for_gpe12345$SV)[,order(abs(t(model_svm_DALY_final_for_gpe12345$coefs)%*%model_svm_DALY_final_for_gpe12345$SV),decreasing = TRUE)]

#10 most importants
(t(model_svm_DALY_final_for_gpe12345$coefs)%*%model_svm_DALY_final_for_gpe12345$SV)[,order(abs(t(model_svm_DALY_final_for_gpe12345$coefs)%*%model_svm_DALY_final_for_gpe12345$SV),decreasing = TRUE)][1:10]

#b
model_svm_DALY_final_for_gpe12345$rho

#prediction on 102 NA from USEtox
usetox_DALY_NA<-merge(usetox_keep[is.na(usetox_keep$DALY),],descriptors,by="row.names")
rownames(usetox_DALY_NA)<-usetox_DALY_NA$Row.names
nrow(usetox_DALY_NA)
usetox_DALY_NA<-usetox_DALY_NA[,-1]
colnames(classes_commun)<-c("CAS","Gpe")
usetox_DALY_NA_gpe<-merge(usetox_DALY_NA,classes_commun,by="CAS")
usetox_DALY_NA_gpe$DALY_predict_global<-0
usetox_DALY_NA_gpe$DALY_predict_local<-0
#nipals
nipals_NA_100<-nipals(X=rbind(total[,6:45],usetox_DALY_NA_gpe[,4:43]), ncomp = 10, reconst = TRUE, max.iter = 500, tol = 1e-09)
usetox_DALY_NA_gpe[,4:43]<-nipals_NA_100$rec[c((nrow(total)+1):nrow(nipals_NA_100$rec)),]

#prediction best global model
usetox_DALY_NA_gpe$DALY_predict_global<-predict(model_svm_DALY_final_for_gpe12345,usetox_DALY_NA_gpe[,4:43])

#prediction best local model (just for cluster 1)
usetox_DALY_NA_gpe$DALY_predict_local<-usetox_DALY_NA_gpe$DALY_predict_global
#gpe 1 RF gpe
tune.RF_gpe1_final<-tune.randomForest(x=total[total$Gpe==1,6:45],y=total[total$Gpe==1,]$log10DALY,mtry=c(3:30))
tune.RF_gpe1_final$best.parameters
model_RFgpe1_DALY_final_for_gpe1<-randomForest(x=total[total$Gpe==1,6:45],y=total[total$Gpe==1,]$log10DALY,importance=TRUE,na.action=na.roughfix,mtry=as.integer(tune.RF_gpe1_final$best.parameters))

varImpPlot(model_RFgpe1_DALY_final_for_gpe1,n.var=10)

usetox_DALY_NA_gpe[usetox_DALY_NA_gpe$Gpe==1,]$DALY_predict_local<-predict(model_RFgpe1_DALY_final_for_gpe1,usetox_DALY_NA_gpe[usetox_DALY_NA_gpe$Gpe==1,4:43])

#computation of the prediction intervals (leave one out)
#for global SVM 
int_prediction<-matrix(NA,nrow=nrow(usetox_DALY_NA_gpe),ncol=nrow(total))
for (i in 1:nrow(total))
{model_svm_DALY_final_for_gpe12345_loo<-svm(x=total[-i,6:45],y=total$log10DALY[-i],gamma=tune_svm_total$best.parameters[1],cost=tune_svm_total$best.parameters[2])
int_prediction[,i]<-predict(model_svm_DALY_final_for_gpe12345_loo,usetox_DALY_NA_gpe[,4:43])
}

usetox_DALY_NA_gpe$int_prediction_global_lower<-apply(int_prediction,1,quantile,probs=c(0.025))
usetox_DALY_NA_gpe$int_prediction_global_upper<-apply(int_prediction,1,quantile,probs=c(0.975))

#for cluster then RF (cluster 1)
total_gpe1<-total[total$Gpe==1,]
int_prediction_loc<-matrix(NA,nrow=nrow(usetox_DALY_NA_gpe[usetox_DALY_NA_gpe$Gpe==1,]),ncol=nrow(total_gpe1))
for (i in 1:nrow(total_gpe1))
{ model_RFgpe1_DALY_final_for_gpe1_loo<-randomForest(x=total_gpe1[-i,6:45],y=total_gpe1$log10DALY[-i],importance=TRUE,na.action=na.roughfix,mtry=as.integer(tune.RF_gpe1_final$best.parameters))
  int_prediction_loc[,i]<-predict(model_RFgpe1_DALY_final_for_gpe1_loo,usetox_DALY_NA_gpe[usetox_DALY_NA_gpe$Gpe==1,4:43])
}

usetox_DALY_NA_gpe$int_prediction_local_lower<-usetox_DALY_NA_gpe$int_prediction_global_lower
usetox_DALY_NA_gpe$int_prediction_local_upper<-usetox_DALY_NA_gpe$int_prediction_global_upper

usetox_DALY_NA_gpe[usetox_DALY_NA_gpe$Gpe==1,]$int_prediction_local_lower<-apply(int_prediction_loc,1,quantile,probs=c(0.025))
usetox_DALY_NA_gpe[usetox_DALY_NA_gpe$Gpe==1,]$int_prediction_local_upper<-apply(int_prediction_loc,1,quantile,probs=c(0.975))

#results outputs
usetox_name<-read.csv("usetox_name.csv", header=T,sep=";",na.strings =c("n/a"))
colnames(usetox_name)[1]<-"CAS"
DALY_prediction<-merge(usetox_name,usetox_DALY_NA_gpe,by="CAS")
DALY_prediction$DALY_predict_global<-signif(10**(DALY_prediction$DALY_predict_global)-epsilon,digits=2)
DALY_prediction$DALY_predict_local<-signif(10**(DALY_prediction$DALY_predict_local)-epsilon,digits=2)
DALY_prediction$int_prediction_local_lower<-signif(10**(DALY_prediction$int_prediction_local_lower)-epsilon,digits=2)
DALY_prediction$int_prediction_local_upper<-signif(10**(DALY_prediction$int_prediction_local_upper)-epsilon,digits=2)
write.csv(DALY_prediction[,c(1,2,46,50,51)],file="Prediction_DALY.csv")

#The same for the PDF###############################################
#independant script 
rm(list = ls())
setwd("C:/Users/Remi/Dropbox/ACV Usetox/Code R et rapports/Rstudio")

set.seed(150720)
#upload the data
usetox<-read.csv("usetox.csv", header=F,sep=";",na.strings =c("n/a"))
usetox<-usetox[1:3077,]
colnames(usetox)<-c("CAS","DALY","PDF")
rownames(usetox)<-usetox$CAS

load("TyPol_2020_07_23.Rdata")
descriptors<-results$X
colnames(descriptors)<-c("C0","C1","C2","C3","C4","C5","Elec_dip_mom","HOMO","LUMO",  "Molecular_mass" ,"Molec_surface_area", "Nb_C", "Nb_Chl"   ,       "Nb_Fluo"     , "Nb_H"    ,      "Nb_Nitro"         , "Nb_O"           , "Nb_Phos"       ,"Nb_S"            ,"Nb_aromatic_bonds"         , "Nb_atoms"                   ,"Nb_bonds"                  ,"Nb_circuits"                ,"Nb_double_bonds"           , "Nb_halogen"           ,"Nb_multiple_bonds"         ,"Nb_non_H_atoms"             ,"Nb_non_H_bonds"            , "Nb_rings"                   ,"Nb_rotable_bonds"          ,"Nb_triple_bonds"            ,"Pola"                   ,"Sum_of_conv"    ,"Total_energy"                     ,"V0"  ,"V1" , "V2"  ,"V3", "V4"  ,"V5" )

#keep in common
common_U_X_2101<-intersect(rownames(usetox),rownames(descriptors)) 
usetox_keep<-usetox[common_U_X_2101,]
descriptors_keep<-descriptors[common_U_X_2101,]

#upload of TyPol
head(results$analyse$classes_mol) ##classif
rownames(results$analyse$classes_mol)<-results$analyse$classes_mol[,1]
classes_commun<-results$analyse$classes_mol[common_U_X_2101,]

#check of NA
length(usetox_keep$PDF)##274
sum(is.na(usetox_keep$PDF)==F)##259 
usetox_keep_PDF_sans_NA<-usetox_keep[is.na(usetox_keep$PDF)==F,]

table(classes_commun[,2]) 
usetox_keep_PDF_sans_NA_c<-merge(usetox_keep_PDF_sans_NA,classes_commun,by="row.names")
table(usetox_keep_PDF_sans_NA_c$V2) 

#train/test procedure
library(mixOmics)
library(randomForest)
library(e1071)

Error<-matrix(NA,ncol=7)
colnames(Error)<-c("RF_pred","PLS_pred","SVM_pred","Gpe_RF_pred","Gpe_SVM_pred","Gpe_PLS_pred","Gpe")
nbboot<-200
if (!file.exists("Error_PDF.rds"))
{for (tt in (1:nbboot))
{ print(tt)  
  #Train/test
  #15% in each cluster : 5 in cluster 1, 17 in 2, 12 in 3, 3 in 4
  gpe1<-usetox_keep_PDF_sans_NA_c[usetox_keep_PDF_sans_NA_c$V2=="1",]
  gpe2<-usetox_keep_PDF_sans_NA_c[usetox_keep_PDF_sans_NA_c$V2=="2",]
  gpe3<-usetox_keep_PDF_sans_NA_c[usetox_keep_PDF_sans_NA_c$V2=="3",]
  gpe4<-usetox_keep_PDF_sans_NA_c[usetox_keep_PDF_sans_NA_c$V2=="4",]
  gpe5<-usetox_keep_PDF_sans_NA_c[usetox_keep_PDF_sans_NA_c$V2=="5",]
  sample1<-sample(c(1:nrow(gpe1)),size=5)
  sample2<-sample(c(1:nrow(gpe2)),size=17)
  sample3<-sample(c(1:nrow(gpe3)),size=12)
  sample4<-sample(c(1:nrow(gpe4)),size=3)
  Valid1<-gpe1[sample1,]
  Valid2<-gpe2[sample2,]
  Valid3<-gpe3[sample3,]
  Valid4<-gpe4[sample4,]
  
  #test set (with the only compound of cluster 5)
  usetox_keep_PDF_test<-rbind(Valid1,Valid2,Valid3,Valid4,gpe5)
  rownames(usetox_keep_PDF_test)<-usetox_keep_PDF_test$CAS
  usetox_keep_PDF_test<-usetox_keep_PDF_test[,c(2,3,4,6)]
  colnames(usetox_keep_PDF_test)[4]<-"Cluster"
  
  #train set
   usetox_keep_PDF_train<-usetox_keep_PDF_sans_NA
  for (i in (1:nrow(usetox_keep_PDF_test)))
  {usetox_keep_PDF_train<-usetox_keep_PDF_train[-which(usetox_keep_PDF_train$CAS==usetox_keep_PDF_test$CAS[i]),]}
  
  train_total<-merge(usetox_keep_PDF_train,descriptors,by="row.names")
  test_total<-merge(usetox_keep_PDF_test,descriptors,by="row.names")
  train_total$log10PDF<-log10(train_total$PDF)
  test_total$log10PDF<-log10(test_total$PDF)
  
  #modelisation
  #1. Nipals for train set
  pls_nipals_train<-nipals(X=train_total[,5:44], ncomp = 10, reconst = TRUE, max.iter = 500, tol = 1e-09)
  data_nipals<-cbind(pls_nipals_train$rec,train_total$log10PDF)
  colnames(data_nipals)[ncol(data_nipals)]<-"log10PDF"
  rownames(data_nipals)<-train_total$Row.names
  
  ff<-as.formula(log10PDF~C0+C1+C2+C3+C4+C5+Elec_dip_mom+HOMO+LUMO+Molecular_mass+Molec_surface_area+Nb_C+Nb_Chl+Nb_Fluo+Nb_H+Nb_Nitro+Nb_O+Nb_Phos+Nb_S+Nb_aromatic_bonds+Nb_atoms+Nb_bonds+Nb_circuits+Nb_double_bonds+Nb_halogen+Nb_multiple_bonds+Nb_non_H_atoms+Nb_non_H_bonds+Nb_rings+Nb_rotable_bonds+Nb_triple_bonds+Pola+Sum_of_conv+Total_energy+V0+V1+V2+V3+V4+V5)
  #2. rf
  tune.RF<-tune.randomForest(x=pls_nipals_train$rec,y=train_total$log10PDF,mtry=c(3:30))
  tune.RF$best.parameters 
  model_RF_PDF<-randomForest(ff,data=data_nipals,importance=TRUE,na.action=na.roughfix,mtry=as.integer(tune.RF$best.parameters))
  
  #pls
  model_pre_pls_PDF<-pls(X=pls_nipals_train$rec,Y=train_total$log10PDF,ncomp=10)
  tune.pls <- perf(model_pre_pls_PDF, validation = 'Mfold', folds=10,criterion = 'all', test.keepX=40,measure="MSE")
  model_pls_PDF<-pls(X=pls_nipals_train$rec,Y=train_total$log10PDF,ncomp=which.min(tune.pls$MSEP))
  
  #svm
  tune_svm<-tune.svm(x=pls_nipals_train$rec,y=train_total$log10PDF, gamma = 10^(-3:3), cost = 10^(-3:3))
  model_svm_PDF<-svm(x=pls_nipals_train$rec,y=train_total$log10PDF,gamma=tune_svm$best.parameters[1],cost=tune_svm$best.parameters[2])
  
  #clustering then RF (then SVM and then PLS)
  rownames(gpe1)<-gpe1$Row.names
  Gpe1_train<-merge(gpe1,data_nipals,by="row.names")
  colnames(Gpe1_train)[43:44]<-c("V1","V2")
  tune.RF_gpe1<-tune.randomForest(x=Gpe1_train[,8:47],y=Gpe1_train$log10PDF,mtry=c(3:30))
  tune.RF_gpe1$best.parameters 
  model_RF_PDF_gpe1<-randomForest(x=Gpe1_train[,8:47],y=Gpe1_train$log10PDF,importance=TRUE,na.action=na.roughfix,mtry=as.integer(tune.RF_gpe1$best.parameters))
  tune_svm1<-tune.svm(x=Gpe1_train[,8:47],y=Gpe1_train$log10PDF, gamma = 10^(-3:3), cost = 10^(-3:3))
  model_svm1_PDF<-svm(x=Gpe1_train[,8:47],y=Gpe1_train$log10PDF,gamma=tune_svm1$best.parameters[1],cost=tune_svm1$best.parameters[2])
  model_pre_pls1_PDF<-pls(X=Gpe1_train[,8:47],Y=Gpe1_train$log10PDF,ncomp=8)
  tune.pls1<- perf(model_pre_pls1_PDF, validation = 'Mfold', folds=10,criterion = 'all', test.keepX=40,measure="MSE")
  model_pls1_PDF<-pls(X=Gpe1_train[,8:47],Y=Gpe1_train$log10PDF,ncomp=which.min(tune.pls1$MSEP))
  
  rownames(gpe2)<-gpe2$Row.names
  Gpe2_train<-merge(gpe2,data_nipals,by="row.names")
  colnames(Gpe2_train)[43:44]<-c("V1","V2")
  tune.RF_gpe2<-tune.randomForest(x=Gpe2_train[,8:47],y=Gpe2_train$log10PDF,mtry=c(3:30))
  tune.RF_gpe2$best.parameters 
  model_RF_PDF_gpe2<-randomForest(x=Gpe2_train[,8:47],y=Gpe2_train$log10PDF,importance=TRUE,na.action=na.roughfix,mtry=as.integer(tune.RF_gpe2$best.parameters))
  tune_svm2<-tune.svm(x=Gpe2_train[,8:47],y=Gpe2_train$log10PDF, gamma = 10^(-3:3), cost = 10^(-3:3))
  model_svm2_PDF<-svm(x=Gpe2_train[,8:47],y=Gpe2_train$log10PDF,gamma=tune_svm2$best.parameters[1],cost=tune_svm2$best.parameters[2])
  model_pre_pls2_PDF<-pls(X=Gpe2_train[,8:47],Y=Gpe2_train$log10PDF,ncomp=8)
  tune.pls2<- perf(model_pre_pls2_PDF, validation = 'Mfold', folds=10,criterion = 'all', test.keepX=40,measure="MSE")
  model_pls2_PDF<-pls(X=Gpe2_train[,8:47],Y=Gpe2_train$log10PDF,ncomp=which.min(tune.pls2$MSEP))
  
  rownames(gpe3)<-gpe3$Row.names
  Gpe3_train<-merge(gpe3,data_nipals,by="row.names")
  colnames(Gpe3_train)[43:44]<-c("V1","V2")
  tune.RF_gpe3<-tune.randomForest(x=Gpe3_train[,8:47],y=Gpe3_train$log10PDF,mtry=c(3:30))
  tune.RF_gpe3$best.parameters 
  model_RF_PDF_gpe3<-randomForest(x=Gpe3_train[,8:47],y=Gpe3_train$log10PDF,importance=TRUE,na.action=na.roughfix,mtry=as.integer(tune.RF_gpe3$best.parameters))
  tune_svm3<-tune.svm(x=Gpe3_train[,8:47],y=Gpe3_train$log10PDF, gamma = 10^(-3:3), cost = 10^(-3:3))
  model_svm3_PDF<-svm(x=Gpe3_train[,8:47],y=Gpe3_train$log10PDF,gamma=tune_svm3$best.parameters[1],cost=tune_svm3$best.parameters[2])
  model_pre_pls3_PDF<-pls(X=Gpe3_train[,8:47],Y=Gpe3_train$log10PDF,ncomp=8)
  tune.pls3<- perf(model_pre_pls3_PDF, validation = 'Mfold', folds=10,criterion = 'all', test.keepX=40,measure="MSE")
  model_pls3_PDF<-pls(X=Gpe3_train[,8:47],Y=Gpe3_train$log10PDF,ncomp=which.min(tune.pls3$MSEP))
  
  rownames(gpe4)<-gpe4$Row.names
  Gpe4_train<-merge(gpe4,data_nipals,by="row.names")
  colnames(Gpe4_train)[43:44]<-c("V1","V2")
  tune.RF_gpe4<-tune.randomForest(x=Gpe4_train[,8:47],y=Gpe4_train$log10PDF,mtry=c(3:30))
  tune.RF_gpe4$best.parameters 
  model_RF_PDF_gpe4<-randomForest(x=Gpe4_train[,8:47],y=Gpe4_train$log10PDF,importance=TRUE,na.action=na.roughfix,mtry=as.integer(tune.RF_gpe4$best.parameters))
  tune_svm4<-tune.svm(x=Gpe4_train[,8:47],y=Gpe4_train$log10PDF, gamma = 10^(-3:3), cost = 10^(-3:3))
  model_svm4_PDF<-svm(x=Gpe4_train[,8:47],y=Gpe4_train$log10PDF,gamma=tune_svm4$best.parameters[1],cost=tune_svm4$best.parameters[2])
  model_pre_pls4_PDF<-pls(X=Gpe4_train[,8:47],Y=Gpe4_train$log10PDF,ncomp=8)
  tune.pls4<- perf(model_pre_pls4_PDF, validation = 'Mfold', folds=10,criterion = 'all', test.keepX=40,measure="MSE")
  model_pls4_PDF<-pls(X=Gpe4_train[,8:47],Y=Gpe4_train$log10PDF,ncomp=which.min(tune.pls4$MSEP))
  
  #3.prediction
  results_PDF<-matrix(NA,ncol=9,nrow=nrow(test_total))
  rownames(results_PDF)<-test_total$Row.names
  colnames(results_PDF)<-c("RF_pred","PLS_pred","SVM_pred","Gpe_RF_pred","Gpe_SVM_pred","Gpe_PLS_pred","Combination_pred","Gpe","Real_value")
  results_PDF[,ncol(results_PDF)]<-test_total$log10PDF
  
  #nipals for test set
  test_nipals<-nipals(X=rbind(train_total[,5:44],test_total[,6:45]), ncomp = 10, reconst = TRUE, max.iter = 500, tol = 1e-09)
  test_total[,6:45]<-test_nipals$rec[(nrow(test_nipals$rec)-nrow(test_total)+1):nrow(test_nipals$rec),]
  
  results_PDF[,1]<-predict(model_RF_PDF,test_total[,6:45])
  results_PDF[,2]<-predict(model_pls_PDF,test_total[,6:45])$predict[,,which.min(tune.pls$MSEP)]
  results_PDF[,3]<-predict(model_svm_PDF,test_total[,6:45])
  
  #per cluster
  results_PDF[,8]<-merge(results$analyse$classes_mol,results_PDF,by="row.names")$V2
  l4<-which(results_PDF[,8]==4)
  results_PDF[l4,4]<-predict(model_RF_PDF_gpe4,test_total[l4,6:45]) 
  results_PDF[l4,5]<-predict(model_svm4_PDF,test_total[l4,6:45]) 
  results_PDF[l4,6]<-predict(model_pls4_PDF,test_total[l4,6:45])$predict[,,which.min(tune.pls4$MSEP)] 
  l3<-which(results_PDF[,8]==3)
  results_PDF[l3,4]<-predict(model_RF_PDF_gpe3,test_total[l3,6:45])
  results_PDF[l3,5]<-predict(model_svm3_PDF,test_total[l3,6:45]) 
  results_PDF[l3,6]<-predict(model_pls3_PDF,test_total[l3,6:45])$predict[,,which.min(tune.pls3$MSEP)] 
  l2<-which(results_PDF[,8]==2)
  results_PDF[l2,4]<-predict(model_RF_PDF_gpe2,test_total[l2,6:45]) 
  results_PDF[l2,5]<-predict(model_svm2_PDF,test_total[l2,6:45]) 
  results_PDF[l2,6]<-predict(model_pls2_PDF,test_total[l2,6:45])$predict[,,which.min(tune.pls2$MSEP)] 
  l1<-which(results_PDF[,8]==1)
  results_PDF[l1,4]<-predict(model_RF_PDF_gpe1,test_total[l1,6:45]) 
  results_PDF[l1,5]<-predict(model_svm1_PDF,test_total[l1,6:45]) 
  results_PDF[l1,6]<-predict(model_pls1_PDF,test_total[l1,6:45])$predict[,,which.min(tune.pls1$MSEP)] 
  
  #Results per method
  Error_bcle<-matrix(NA,ncol=7,nrow=nrow(test_total))
  colnames(Error_bcle)<-c("RF_pred","PLS_pred","SVM_pred","Gpe_RF_pred","Gpe_SVM_pred","Gpe_PLS_pred","Gpe")
  Error_bcle[,1]<-results_PDF[,1]-results_PDF[,9]
  Error_bcle[,2]<-results_PDF[,2]-results_PDF[,9]
  Error_bcle[,3]<-results_PDF[,3]-results_PDF[,9]
  Error_bcle[,4]<-results_PDF[,4]-results_PDF[,9]
  Error_bcle[,5]<-results_PDF[,5]-results_PDF[,9]
  Error_bcle[,6]<-results_PDF[,6]-results_PDF[,9]
  Error_bcle[,7]<-results_PDF[,8]
  Error<-rbind(Error,Error_bcle)
}
  Error<-Error[-1,]
  save(Error,file="Error_PDF.rds")
}
#scatterplot
nbscatterboot<-500
scatter_PDF<-cbind(usetox_keep_PDF_sans_NA,matrix(NA,ncol=nbscatterboot,nrow=nrow(usetox_keep_PDF_sans_NA)))
if (!file.exists("scatter_PDF.rds"))
{for (tt in (1:nbscatterboot))
{ print(tt)  
  #Train/test
  gpe1<-usetox_keep_PDF_sans_NA_c[usetox_keep_PDF_sans_NA_c$V2=="1",]
  gpe2<-usetox_keep_PDF_sans_NA_c[usetox_keep_PDF_sans_NA_c$V2=="2",]
  gpe3<-usetox_keep_PDF_sans_NA_c[usetox_keep_PDF_sans_NA_c$V2=="3",]
  gpe4<-usetox_keep_PDF_sans_NA_c[usetox_keep_PDF_sans_NA_c$V2=="4",]
  gpe5<-usetox_keep_PDF_sans_NA_c[usetox_keep_PDF_sans_NA_c$V2=="5",]
  sample1<-sample(c(1:nrow(gpe1)),size=5)
  sample2<-sample(c(1:nrow(gpe2)),size=17)
  sample3<-sample(c(1:nrow(gpe3)),size=12)
  sample4<-sample(c(1:nrow(gpe4)),size=3)
  Valid1<-gpe1[sample1,]
  Valid2<-gpe2[sample2,]
  Valid3<-gpe3[sample3,]
  Valid4<-gpe4[sample4,]
  
  #test set
  usetox_keep_PDF_test<-rbind(Valid1,Valid2,Valid3,Valid4,gpe5)
  rownames(usetox_keep_PDF_test)<-usetox_keep_PDF_test$CAS
  usetox_keep_PDF_test<-usetox_keep_PDF_test[,c(2,3,4,6)]
  colnames(usetox_keep_PDF_test)[4]<-"Cluster"
  
  #train set
  usetox_keep_PDF_train<-usetox_keep_PDF_sans_NA
  for (i in (1:nrow(usetox_keep_PDF_test)))
  {usetox_keep_PDF_train<-usetox_keep_PDF_train[-which(usetox_keep_PDF_train$CAS==usetox_keep_PDF_test$CAS[i]),]}
  
  train_total<-merge(usetox_keep_PDF_train,descriptors,by="row.names")
  test_total<-merge(usetox_keep_PDF_test,descriptors,by="row.names")
  train_total$log10PDF<-log10(train_total$PDF)
  test_total$log10PDF<-log10(test_total$PDF)
  
  #modelisation
  #1. Nipals for train
  pls_nipals_train<-nipals(X=train_total[,5:44], ncomp = 10, reconst = TRUE, max.iter = 500, tol = 1e-09)
  data_nipals<-cbind(pls_nipals_train$rec,train_total$log10PDF)
  colnames(data_nipals)[ncol(data_nipals)]<-"log10PDF"
  rownames(data_nipals)<-train_total$Row.names
  
  ff<-as.formula(log10PDF~C0+C1+C2+C3+C4+C5+Elec_dip_mom+HOMO+LUMO+Molecular_mass+Molec_surface_area+Nb_C+Nb_Chl+Nb_Fluo+Nb_H+Nb_Nitro+Nb_O+Nb_Phos+Nb_S+Nb_aromatic_bonds+Nb_atoms+Nb_bonds+Nb_circuits+Nb_double_bonds+Nb_halogen+Nb_multiple_bonds+Nb_non_H_atoms+Nb_non_H_bonds+Nb_rings+Nb_rotable_bonds+Nb_triple_bonds+Pola+Sum_of_conv+Total_energy+V0+V1+V2+V3+V4+V5)
  #2. modelisation
  #cluster then SVM for cluster 1 and 4 and cluster then RF for cluster 2 and 3, SVM global pour 5  
  rownames(gpe1)<-gpe1$Row.names
  Gpe1_train<-merge(gpe1,data_nipals,by="row.names")
  colnames(Gpe1_train)[43:44]<-c("V1","V2")
  tune_svm1<-tune.svm(x=Gpe1_train[,8:47],y=Gpe1_train$log10PDF, gamma = 10^(-3:3), cost = 10^(-3:3))
  model_svm1_PDF<-svm(x=Gpe1_train[,8:47],y=Gpe1_train$log10PDF,gamma=tune_svm1$best.parameters[1],cost=tune_svm1$best.parameters[2])
  
  rownames(gpe4)<-gpe4$Row.names
  Gpe4_train<-merge(gpe4,data_nipals,by="row.names")
  colnames(Gpe4_train)[43:44]<-c("V1","V2")
  tune_svm4<-tune.svm(x=Gpe4_train[,8:47],y=Gpe4_train$log10PDF, gamma = 10^(-3:3), cost = 10^(-3:3))
  model_svm4_PDF<-svm(x=Gpe4_train[,8:47],y=Gpe4_train$log10PDF,gamma=tune_svm4$best.parameters[1],cost=tune_svm4$best.parameters[2])
  
  rownames(gpe2)<-gpe2$Row.names
  Gpe2_train<-merge(gpe2,data_nipals,by="row.names")
  colnames(Gpe2_train)[43:44]<-c("V1","V2")
  tune.RF_gpe2<-tune.randomForest(x=Gpe2_train[,8:47],y=Gpe2_train$log10PDF,mtry=c(3:30))
  tune.RF_gpe2$best.parameters 
  model_RF_PDF_gpe2<-randomForest(x=Gpe2_train[,8:47],y=Gpe2_train$log10PDF,importance=TRUE,na.action=na.roughfix,mtry=as.integer(tune.RF_gpe2$best.parameters))
 
  rownames(gpe3)<-gpe3$Row.names
  Gpe3_train<-merge(gpe3,data_nipals,by="row.names")
  colnames(Gpe3_train)[43:44]<-c("V1","V2")
  tune.RF_gpe3<-tune.randomForest(x=Gpe3_train[,8:47],y=Gpe3_train$log10PDF,mtry=c(3:30))
  tune.RF_gpe3$best.parameters 
  model_RF_PDF_gpe3<-randomForest(x=Gpe3_train[,8:47],y=Gpe3_train$log10PDF,importance=TRUE,na.action=na.roughfix,mtry=as.integer(tune.RF_gpe3$best.parameters))
  
  #svm global
  tune_svm<-tune.svm(x=pls_nipals_train$rec,y=train_total$log10PDF, gamma = 10^(-3:3), cost = 10^(-3:3))
  model_svm_PDF<-svm(x=pls_nipals_train$rec,y=train_total$log10PDF,gamma=tune_svm$best.parameters[1],cost=tune_svm$best.parameters[2])
  
  #3. prediction
  results_PDF<-matrix(NA,ncol=9,nrow=nrow(test_total))
  rownames(results_PDF)<-test_total$Row.names
  colnames(results_PDF)<-c("RF_pred","PLS_pred","SVM_pred","Gpe_RF_pred","Gpe_SVM_pred","Gpe_PLS_pred","Combination_pred","Gpe","Real_value")
  results_PDF[,ncol(results_PDF)]<-test_total$log10PDF
  results_PDF[,8]<-merge(results$analyse$classes_mol,results_PDF,by="row.names")$V2
  
  #nipals for test 
  test_nipals<-nipals(X=rbind(train_total[,5:44],test_total[,6:45]), ncomp = 10, reconst = TRUE, max.iter = 500, tol = 1e-09)
  test_total[,6:45]<-test_nipals$rec[(nrow(test_nipals$rec)-nrow(test_total)+1):nrow(test_nipals$rec),]
  
  results_PDF[,3]<-predict(model_svm_PDF,test_total[,6:45])
  l4<-which(results_PDF[,8]==4)
  results_PDF[l4,5]<-predict(model_svm4_PDF,test_total[l4,6:45]) 
  l3<-which(results_PDF[,8]==3)
  results_PDF[l3,4]<-predict(model_RF_PDF_gpe3,test_total[l3,6:45])
  l2<-which(results_PDF[,8]==2)
  results_PDF[l2,4]<-predict(model_RF_PDF_gpe2,test_total[l2,6:45]) 
  l1<-which(results_PDF[,8]==1)
  results_PDF[l1,5]<-predict(model_svm1_PDF,test_total[l1,6:45]) 
  
  #cluster then SVM for cluster 1 and 4 and cluster then RF for cluster 2 and 3, SVM pour 5  
  for (jj in 1:nrow(results_PDF))
  { if (results_PDF[jj,8]==5) {scatter_PDF[rownames(results_PDF)[jj],tt+3]<-results_PDF[jj,3]}
    {if ((results_PDF[jj,8]==2)|(results_PDF[jj,8]==3)) ##cluster-RF
      { scatter_PDF[rownames(results_PDF)[jj],tt+3]<-results_PDF[jj,4]}
      else ##cluster-SVM
      {scatter_PDF[rownames(results_PDF)[jj],tt+3]<-results_PDF[jj,5]}
    }
  }  
  save(scatter_PDF,file="scatter_PDF.rds")  
}  
}

load("scatter_PDF.rds")
scatter_PDF<-as.data.frame(scatter_PDF)
scatter_PDF$med_est<-apply(scatter_PDF[4:(nbboot+3)],1,median,na.rm=T)
scatter_PDF$med_est05<-apply(scatter_PDF[4:(nbboot+3)],1,quantile,probs=0.05,na.rm=T)
scatter_PDF$med_est95<-apply(scatter_PDF[4:(nbboot+3)],1,quantile,probs=0.95,na.rm=T)
scatter_PDF$log10PDF<-log10(scatter_PDF$PDF)
scatter_PDF_plot<-scatter_PDF[-185,]
ggplot(data = scatter_PDF_plot,aes(x=log10PDF, y=med_est,ymin=med_est05,ymax=med_est95))+geom_point()+geom_abline(slope=1,intercept=0)+geom_errorbar()+xlab(expression(CF[ET]))+ylab("Estimation")
dev.print(png, file = "scatter_PDF.png", width = 1600, height = 1500, res = 150)

load("Error_PDF.rds")
colnames(Error)[1:7]<-c("RF","PLS","SVM","Cluster-RF","Cluster-SVM","Cluster-PLS","Gpe")

#performances on all the data
colMeans(abs(Error),na.rm=T)

col_boxplot<-apply(abs(Error),2,median,na.rm=T)

library("RColorBrewer")
palette_col<-rev(brewer.pal(n = 6, name = 'RdYlGn'))
boxplot(abs(Error[,c(1,2,3,4,6,5)]),outline=F,ylab="Absolute Error",col=palette_col[rank(col_boxplot[c(1,2,3,4,6,5)])])
dev.print(png, file = "boxplot_PDF_AE.png", width = 1600, height = 1500, res = 150)

#Performances per cluster
col_gpe1<-apply(abs(Error[Error[,7]==1,]),2,median,na.rm=T)
col_gpe2<-apply(abs(Error[Error[,7]==2,]),2,median,na.rm=T)
col_gpe3<-apply(abs(Error[Error[,7]==3,]),2,median,na.rm=T)
col_gpe4<-apply(abs(Error[Error[,7]==4,]),2,median,na.rm=T)

par(mfrow=c(2,2))
boxplot(abs(Error[Error[,7]==1,c(1,2,3,4,6,5)]),ylim=c(0,5),outline=F,ylab="Absolute Error",main="Cluster 1",col=palette_col[rank(col_gpe1[c(1,2,3,4,6,5)])])
boxplot(abs(Error[Error[,7]==2,c(1,2,3,4,6,5)]),ylim=c(0,5),outline=F,ylab="Absolute Error",main="Cluster 2",col=palette_col[rank(col_gpe2[c(1,2,3,4,6,5)])])
boxplot(abs(Error[Error[,7]==3,c(1,2,3,4,6,5)]),ylim=c(0,5),outline=F,ylab="Absolute Error",main="Cluster 3",col=palette_col[rank(col_gpe3[c(1,2,3,4,6,5)])])
boxplot(abs(Error[Error[,7]==4,c(1,2,3,4,6,5)]),ylim=c(0,5),outline=F,ylab="Absolute Error",main="Cluster 4",col=palette_col[rank(col_gpe4[c(1,2,3,4,6,5)])])
dev.print(png, file = "boxplot_PDF_AE_gpe.png", width = 2500, height = 1500, res = 150)

#best model on all the data
usetox_keep_PDF_sans_NA_c<-usetox_keep_PDF_sans_NA_c[,-c(1,5)]
colnames(usetox_keep_PDF_sans_NA_c)[4]<-"Gpe"
rownames(usetox_keep_PDF_sans_NA_c)<-usetox_keep_PDF_sans_NA_c$CAS
total<-merge(usetox_keep_PDF_sans_NA_c,descriptors,by="row.names")
total$log10PDF<-log10(total$PDF)

#Nipals
nipals_total<-nipals(X=total[,6:45], ncomp = 10, reconst = TRUE, max.iter = 500, tol = 1e-09)
total[,6:45]<-nipals_total$rec

#best model selected on all the data (cluster then RF)
#gpe 1
tune.RFgpe1_for_gpe1<-tune.randomForest(x=total[total$Gpe==1,6:45],y=total[total$Gpe==1,]$log10PDF,mtry=c(3:30))
model_RFgpe1_PDF_final_for_gpe1<-randomForest(x=total[total$Gpe==1,6:45],y=total[total$Gpe==1,]$log10PDF,importance=TRUE,na.action=na.roughfix,mtry=as.integer(tune.RFgpe1_for_gpe1$best.parameters))

varImpPlot(model_RFgpe1_PDF_final_for_gpe1,n.var=10)

#gpe 2
tune.RFgpe2_for_gpe2<-tune.randomForest(x=total[total$Gpe==2,6:45],y=total[total$Gpe==2,]$log10PDF,mtry=c(3:30))
model_RFgpe2_PDF_final_for_gpe2<-randomForest(x=total[total$Gpe==2,6:45],y=total[total$Gpe==2,]$log10PDF,importance=TRUE,na.action=na.roughfix,mtry=as.integer(tune.RFgpe2_for_gpe2$best.parameters))

varImpPlot(model_RFgpe2_PDF_final_for_gpe2,n.var=10)

#gpe 3
tune.RFgpe3_for_gpe3<-tune.randomForest(x=total[total$Gpe==3,6:45],y=total[total$Gpe==3,]$log10PDF,mtry=c(3:30))
model_RFgpe3_PDF_final_for_gpe3<-randomForest(x=total[total$Gpe==3,6:45],y=total[total$Gpe==3,]$log10PDF,importance=TRUE,na.action=na.roughfix,mtry=as.integer(tune.RFgpe3_for_gpe3$best.parameters))

varImpPlot(model_RFgpe3_PDF_final_for_gpe3,n.var=10)

#gpe 4
tune.RFgpe4_for_gpe4<-tune.randomForest(x=total[total$Gpe==4,6:45],y=total[total$Gpe==4,]$log10PDF,mtry=c(3:30))
model_RFgpe4_PDF_final_for_gpe4<-randomForest(x=total[total$Gpe==4,6:45],y=total[total$Gpe==4,]$log10PDF,importance=TRUE,na.action=na.roughfix,mtry=as.integer(tune.RFgpe4_for_gpe4$best.parameters))

varImpPlot(model_RFgpe4_PDF_final_for_gpe4,n.var=10)

#important variables
namesRFg1<-names(sort(model_RFgpe1_PDF_final_for_gpe1$importance[,1],decreasing = T)[1:10])
namesRFg2<-names(sort(model_RFgpe2_PDF_final_for_gpe2$importance[,1],decreasing = T)[1:10])
namesRFg3<-names(sort(model_RFgpe3_PDF_final_for_gpe3$importance[,1],decreasing = T)[1:10])
namesRFg4<-names(sort(model_RFgpe4_PDF_final_for_gpe4$importance[,1],decreasing = T)[1:10])

namestot<-data.frame("Cluster 1"=namesRFg1,"Cluster 2"=namesRFg2,"Cluster 3"=namesRFg3,"Cluster 4"=namesRFg4)

write.csv(namestot,file="Names_PDF.csv")

# prediction on 15 NA of USEtox
usetox_PDF_NA<-merge(usetox_keep[is.na(usetox_keep$PDF),],descriptors,by="row.names")
rownames(usetox_PDF_NA)<-usetox_PDF_NA$Row.names
nrow(usetox_PDF_NA)
usetox_PDF_NA<-usetox_PDF_NA[,-1]
colnames(classes_commun)<-c("CAS","Gpe")
usetox_PDF_NA_gpe<-merge(usetox_PDF_NA,classes_commun,by="CAS")
usetox_PDF_NA_gpe$PDF_predict_global<-0
#nipals
nipals_NA_15<-nipals(X=rbind(total[,6:45],usetox_PDF_NA_gpe[,4:43]), ncomp = 10, reconst = TRUE, max.iter = 500, tol = 1e-09)
usetox_PDF_NA_gpe[,4:43]<-nipals_NA_15$rec[c((nrow(total)+1):nrow(nipals_NA_15$rec)),]

#usetox_PDF_NA_gpe[usetox_PDF_NA_gpe$Gpe==1,]$PDF_predict<-predict(model_RFgpe1_PDF_final_for_gpe1,usetox_PDF_NA_gpe[usetox_PDF_NA_gpe$Gpe==1,4:43])
#no compound to predict in cluster 1
usetox_PDF_NA_gpe[usetox_PDF_NA_gpe$Gpe==2,]$PDF_predict_global<-predict(model_RFgpe2_PDF_final_for_gpe2,usetox_PDF_NA_gpe[usetox_PDF_NA_gpe$Gpe==2,4:43])
usetox_PDF_NA_gpe[usetox_PDF_NA_gpe$Gpe==3,]$PDF_predict_global<-predict(model_RFgpe3_PDF_final_for_gpe3,usetox_PDF_NA_gpe[usetox_PDF_NA_gpe$Gpe==3,4:43])
usetox_PDF_NA_gpe[usetox_PDF_NA_gpe$Gpe==4,]$PDF_predict_global<-predict(model_RFgpe4_PDF_final_for_gpe4,usetox_PDF_NA_gpe[usetox_PDF_NA_gpe$Gpe==4,4:43])
#usetox_PDF_NA_gpe[usetox_PDF_NA_gpe$Gpe==5,]$PDF_predict<-predict(model_svm_PDF_final_for_gpe5,usetox_PDF_NA_gpe[usetox_PDF_NA_gpe$Gpe==5,4:43])
# no compound to predict in cluster 5

#using best local models (cluster then SVM for cluster 1 and 4, cluster theb RF gpe (= best global model, computed above) for cluster 2 and 3)
#cluster then SVM cluster 4
usetox_PDF_NA_gpe$PDF_predict_local<-usetox_PDF_NA_gpe$PDF_predict_global
set.seed(47)
tune_svmgpe4_for_gpe4<-tune.svm(x=total[total$Gpe==4,6:45],y=total[total$Gpe==4,]$log10PDF, gamma = 10^(-3:3), cost = 10^(-3:3))
model_svmgpe4_PDF_final_for_gpe4<-svm(x=total[total$Gpe==4,6:45],y=total[total$Gpe==4,]$log10PDF,gamma=tune_svmgpe4_for_gpe4$best.parameters[1],cost=tune_svmgpe4_for_gpe4$best.parameters[2])

#SVM description
length(model_svmgpe4_PDF_final_for_gpe4$index)

(t(model_svmgpe4_PDF_final_for_gpe4$coefs)%*%model_svmgpe4_PDF_final_for_gpe4$SV)[,order(abs(t(model_svmgpe4_PDF_final_for_gpe4$coefs)%*%model_svmgpe4_PDF_final_for_gpe4$SV))]

#10 more important
names((t(model_svmgpe4_PDF_final_for_gpe4$coefs)%*%model_svmgpe4_PDF_final_for_gpe4$SV)[,order(abs(t(model_svmgpe4_PDF_final_for_gpe4$coefs)%*%model_svmgpe4_PDF_final_for_gpe4$SV))][40:31])

#cluster then SVM cluster 1
tune_svmgpe1_for_gpe1<-tune.svm(x=total[total$Gpe==1,6:45],y=total[total$Gpe==1,]$log10PDF, gamma = 10^(-3:3), cost = 10^(-3:3))
model_svmgpe1_PDF_final_for_gpe1<-svm(x=total[total$Gpe==1,6:45],y=total[total$Gpe==1,]$log10PDF,gamma=tune_svmgpe1_for_gpe1$best.parameters[1],cost=tune_svmgpe1_for_gpe1$best.parameters[2])

#SVM description
length(model_svmgpe1_PDF_final_for_gpe1$index)

#10 more important
names((t(model_svmgpe1_PDF_final_for_gpe1$coefs)%*%model_svmgpe1_PDF_final_for_gpe1$SV)[,order(abs(t(model_svmgpe1_PDF_final_for_gpe1$coefs)%*%model_svmgpe1_PDF_final_for_gpe1$SV))][40:31])

usetox_PDF_NA_gpe[usetox_PDF_NA_gpe$Gpe==4,]$PDF_predict_local<-predict(model_svmgpe4_PDF_final_for_gpe4,usetox_PDF_NA_gpe[usetox_PDF_NA_gpe$Gpe==4,4:43])

usetox_name<-read.csv("usetox_name.csv", header=T,sep=";",na.strings =c("n/a"))
colnames(usetox_name)[1]<-"CAS"

#compute prediction interval (leave one out)
usetox_PDF_NA_gpe$int_prediction_local_lower<-NA
usetox_PDF_NA_gpe$int_prediction_local_upper<-NA

#RF (cluster 2)
total_gpe2<-total[total$Gpe==2,]
int_prediction_loc<-matrix(NA,nrow=nrow(usetox_PDF_NA_gpe[usetox_PDF_NA_gpe$Gpe==2,]),ncol=nrow(total_gpe2))
for (i in 1:nrow(total_gpe2))
{ model_RFgpe2_PDF_final_for_gpe2_loo<-randomForest(x=total_gpe2[-i,6:45],y=total_gpe2$log10PDF[-i],importance=TRUE,na.action=na.roughfix,mtry=as.integer(tune.RFgpe2_for_gpe2$best.parameters))
int_prediction_loc[,i]<-predict(model_RFgpe2_PDF_final_for_gpe2_loo,usetox_PDF_NA_gpe[usetox_PDF_NA_gpe$Gpe==2,4:43])
}

usetox_PDF_NA_gpe[usetox_PDF_NA_gpe$Gpe==2,]$int_prediction_local_lower<-apply(int_prediction_loc,1,quantile,probs=c(0.025))
usetox_PDF_NA_gpe[usetox_PDF_NA_gpe$Gpe==2,]$int_prediction_local_upper<-apply(int_prediction_loc,1,quantile,probs=c(0.975))

#RF cluster 3
total_gpe3<-total[total$Gpe==3,]
int_prediction_loc3<-matrix(NA,nrow=nrow(usetox_PDF_NA_gpe[usetox_PDF_NA_gpe$Gpe==3,]),ncol=nrow(total_gpe3))
for (i in 1:nrow(total_gpe3))
{ model_RFgpe3_PDF_final_for_gpe3_loo<-randomForest(x=total_gpe3[-i,6:45],y=total_gpe3$log10PDF[-i],importance=TRUE,na.action=na.roughfix,mtry=as.integer(tune.RFgpe3_for_gpe3$best.parameters))
int_prediction_loc3[,i]<-predict(model_RFgpe3_PDF_final_for_gpe3_loo,usetox_PDF_NA_gpe[usetox_PDF_NA_gpe$Gpe==3,4:43])
}

usetox_PDF_NA_gpe[usetox_PDF_NA_gpe$Gpe==3,]$int_prediction_local_lower<-apply(int_prediction_loc3,1,quantile,probs=c(0.025))
usetox_PDF_NA_gpe[usetox_PDF_NA_gpe$Gpe==3,]$int_prediction_local_upper<-apply(int_prediction_loc3,1,quantile,probs=c(0.975))

#SVM cluster 4
total_gpe4<-total[total$Gpe==4,]
int_prediction_loc4<-matrix(NA,nrow=nrow(usetox_PDF_NA_gpe[usetox_PDF_NA_gpe$Gpe==4,]),ncol=nrow(total_gpe4))
for (i in 1:nrow(total_gpe4))
{ model_svmgpe4_PDF_final_for_gpe4_loo<-svm(x=total_gpe4[-i,6:45],y=total_gpe4$log10PDF[-i],gamma=tune_svmgpe4_for_gpe4$best.parameters[1],cost=tune_svmgpe4_for_gpe4$best.parameters[2])
int_prediction_loc4[,i]<-predict(model_svmgpe4_PDF_final_for_gpe4_loo,usetox_PDF_NA_gpe[usetox_PDF_NA_gpe$Gpe==4,4:43])
}

usetox_PDF_NA_gpe[usetox_PDF_NA_gpe$Gpe==4,]$int_prediction_local_lower<-apply(int_prediction_loc4,1,quantile,probs=c(0.025))
usetox_PDF_NA_gpe[usetox_PDF_NA_gpe$Gpe==4,]$int_prediction_local_upper<-apply(int_prediction_loc4,1,quantile,probs=c(0.975))

#result outputs
PDF_predict<-merge(usetox_name,usetox_PDF_NA_gpe,by="CAS")
PDF_predict$PDF_predict_global<-round(10**PDF_predict$PDF_predict_global,digits=0)
PDF_predict$PDF_predict_local<-round(10**PDF_predict$PDF_predict_local,digits=0)
PDF_predict$int_prediction_local_lower<-round(10**PDF_predict$int_prediction_local_lower,digits=0)
PDF_predict$int_prediction_local_upper<-round(10**PDF_predict$int_prediction_local_upper,digits=0)
write.csv(PDF_predict[,c(1,2,45,47,48,49)],file="Prediction_PDF.csv")
```
